# Temporal dynamics of collateral RNA cleavage by LbuCas13a in human cells

**DOI:** 10.1101/2023.01.19.524716

**Authors:** Jorik F. Bot, Zhihan Zhao, Mengyuan Li, Darnell Kammeron, Peng Shang, Niels Geijsen

## Abstract

CRISPR-Cas13 exclusively targets RNA. In prokaryotic cells, Cas13 cleaves both target and non-target RNA indiscriminately upon activation by a specific target RNA, but in eukaryotic cells collateral cleavage activity has been limited. To investigate collateral cleavage by Cas13 in eukaryotic cells, we first compared various Cas13 orthologs and found that specifically LbuCas13a exhibits strong collateral RNA cleavage activity in human cells when delivered as ribonucleoprotein, independent of cell line and targeting both exogenous and endogenous transcripts. Collateral RNA cleavage started within 50 minutes of ribonucleoprotein delivery resulting in major alterations to the total RNA profile. In response to the collateral RNA cleavage, cells upregulated genes associated with stress and innate immune response, ultimately leading to apoptotic cell death. This enabled us to use LbuCas13a as a flexible and repeatable target-RNA-specific cell elimination tool. Finally, we used Nanopore sequencing to explore the identity of collaterally cleaved RNAs, the nucleotide position at which they are cleaved, and the temporal dynamics of collateral RNA cleavage. This revealed that LbuCas13a activation leads to rapid and global cleavage of cytoplasmic RNAs at specific nucleotide positions. In conclusion, we here report that LbuCas13a has high collateral activity in human cells and describe the temporal dynamics of the collateral RNA cleavage, the cellular responses ultimately leading to apoptosis, how this can be exploited as a cell elimination tool, and the collateral cleavage preferences of LbuCas13a.

## Introduction

CRISPR-Cas is a prokaryotic adaptive immune system that protects bacteria and archaea from invading viruses and plasmids (Marraffini, 2015; Mohanraju et al., 2016; Oost et al., 2014). Since its initial discovery, an impressive diversity of CRISPR-Cas systems has been revealed (Makarova et al., 2020). Especially the Class 2 DNA targeting effector proteins, Cas9 and Cas12, have been extensively studied and have found widespread adoption for biotechnological and clinical use (Liu et al., 2022; Sharma et al., 2021). A recently discovered Class 2 system, CRISPR-Cas13, targets and cleaves RNA. The signature element of CRISPR-Cas13 systems are the two Higher Eukaryotes and Prokaryotes Nucleotide–binding (HEPN) RNase domains of the effector protein (Koonin et al., 2017; Makarova et al., 2015). When a Cas13 protein is activated by a guide-matching target RNA, these domains cooperate to form a single active RNA nuclease site (Abudayyeh et al., 2016; Cox et al., 2017; East-Seletsky et al., 2016; Konermann et al., 2018; Smargon et al., 2017; Xu et al., 2021; Yan et al., 2018). This catalytic site is located on the outside of the protein, facing away from the guide RNA-target RNA complex (Knott et al., 2017; Liu et al., 2017b, 2017a; Slaymaker et al., 2019; Zhang et al., 2018). As a consequence, Cas13 cleaves both target and non-target RNA indiscriminately *in vitro* (Abudayyeh et al., 2016; East-Seletsky et al., 2017, 2016; Konermann et al., 2018; Smargon et al., 2017; Xu et al., 2021; Yan et al., 2018) and in bacteria (Abudayyeh et al., 2016; Meeske et al., 2019; Smargon et al., 2017; Yan et al., 2018) when activated by a specific target RNA. *In vitro,* this collateral RNA cleavage has been employed to develop portable, rapid, and highly sensitive nucleic acid detection methods (East-Seletsky et al., 2017; Gootenberg et al., 2018, 2017; Myhrvold et al., 2018), including for Coronavirus disease 2019 (COVID-19) diagnostics (Arizti-sanz et al., 2020; Fozouni et al., 2021; Patchsung et al., 2020). In bacteria, collateral RNA cleavage probably serves to protect bacterial communities by inducing dormancy after phage infection (Meeske et al., 2019; Mohanraju et al., 2022).

Initially, no evidence of collateral cleavage was found when Cas13 was first tested in eukaryotic cells (Abudayyeh et al., 2017; Cox et al., 2017; Konermann et al., 2018). Instead, Cas13 induced efficient and specific target RNA knockdown. Because Cas13-based knockdown was shown to have higher specificity and efficiency than RNAi (Abudayyeh et al., 2017; Cox et al., 2017; He et al., 2020; Huynh et al., 2020; Konermann et al., 2018; Li et al., 2021; Zhang et al., 2021) and CRISPRi (Huynh et al., 2020; Konermann et al., 2018), it has seen enthusiastic adoption as a specific RNA knockdown tool. For example, Cas13 has been adapted for use in mice (He et al., 2020; Kushawah et al., 2020; Powell et al., 2022; Zhou et al., 2020), fish (Kushawah et al., 2020; Wang et al., 2021), yeast (Jing et al., 2018), fly (Huynh et al., 2020), plants (Aman et al., 2018; Mahas et al., 2019), and as an anti-viral tool (Blanchard et al., 2021; Freije et al., 2019; Tng et al., 2020). However, recently several groups have reported that Cas13 is capable of collateral cleavage in eukaryotes in certain conditions, in some cases causing cytotoxicity (Ai et al., 2022; Buchman et al., 2020; Y. Li et al., 2023; Özcan et al., 2021; Shi et al., 2023; Wang et al., 2019; Xu et al., 2021; Zhang et al., 2019). To further advance CRISPR-Cas13 as a specific RNA knockdown method, some groups have made progress to minimize the Cas13 collateral cleavage effects (Ai et al., 2022; Tong et al., 2022). We took the opposite approach and set out to exploit Cas13’s collateral cleavage activity as a target-RNA-specific cell elimination tool. A comparison of the collateral cleavage activity of different Cas13 orthologs identified LbuCas13a as a particularly active Cas13 variant. In this set out to explore the on-target and collateral RNA cleavage activity of LbuCas13a in cells. We observed that specifically ribonucleoprotein complex (RNP) delivery of LbuCas13a induces robust collateral RNA cleavage in human cells. The collateral activity is cell line and RNP delivery method independent, and is induced by targeting both exogenous and endogenous transcripts. The resulting collateral RNA cleavage leads to transcriptomic changes associated with the activation of an innate immune response. We demonstrate that collateral RNA cleavage triggers target RNA-specific apoptosis, which is strongly correlated with the expression level of the target RNA. Consequently, LbuCas13a can be used as a target RNA-specific cell selection tool which eliminates specific cell types from a heterogenous cell population. We demonstrate that this can be used to enrich for cells that have undergone gene editing. Finally, using Nanopore sequencing, we observed that upon activation by the target RNA, LbuCas13a indiscriminately cleaves cytoplasmic RNAs. This cleavage occurs at very specific positions, often within the loops of stem-loop structures. In conclusion, here we describe the highly active LbuCas13a, the temporal dynamics of target and collateral RNA cleavage, the cellular response to collateral RNA cleavage, the application of LbuCas13a as a negative cell selection tool, and the identity and cleavage positions of the RNAs subject to collateral RNA cleavage by LbuCas13a.

## Results

### LbuCas13a exhibits strong collateral RNA cleavage activity in human cells

The enzymatic activity of different Cas13 orthologs *in vitro* is known to range over seven orders of magnitude (East-Seletsky et al., 2017). We hypothesized that the Cas13 orthologs with the strongest collateral cleavage activity *in vitro* are the most likely to show any collateral cleavage in human cells. Therefore, we compared the collateral RNA cleavage activity of six different CRISPR-Cas13 effectors (LbuCas13a, LwaCas13a, BzCas13b, PspCas13b, RspCas13d and RfxCas13d) *in vitro*, as well as in cells. We produced and purified all six recombinant proteins (Supplementary figure 1A). Next, the *in vitro* collateral RNA cleavage activity of these proteins was assessed using the RNaseAlert® assay (IDT), which quantitatively detects non-specific RNA cleavage. LbuCas13a exhibited the strongest *in vitro* collateral cleavage activity (Supplementary figure 1B). To test these orthologs in human cells, we transfected them as recombinant protein together with guide RNAs into HAP1 cells constitutively overexpressing a fluorescence-inactivated EGFP^Y66S^ (Zhao et al., 2022), referred to as HAP1-dEGFP from hereon. As transfection method we used ‘induced Transduction by Osmocytosis and Propanebetaine’ (iTOP), a previously reported method for the intracellular delivery of recombinant proteins (D’Astolfo et al., 2015). Next, the total RNA integrity was assessed using a Bioanalyzer. The total RNA profile of dEGFP expressing cells transfected with LbuCas13a and a dEGFP targeting guide RNA clearly revealed a degradation pattern mainly consisting of several novel fragments preceding the 18S ribosomal RNA (rRNA) peak (Figure 1A), while the total RNA of wildtype HAP1 cells transfected with LbuCas13a and a dEGFP targeting guide RNA was intact at all timepoints. This collateral RNA cleavage pattern was time-dependent and peaked at 100 to 200 minutes after transfection, with some cleavage fragments remaining at 24 hours after transfection. None of the other Cas13 orthologs were able to generate such a distinct change to the total RNA pattern for any of the target RNAs tested (Supplementary figures 2 and 3). Next we performed an LbuCas13a protein and guide RNA concentration gradient, ranging from 1 µM to 15 µM and using an equimolar ratio of protein and guide RNA (Supplementary figure 4). Collateral cleavage was evident in the total RNA profile from 1 µM and increased logarithmically with the concentration. Because 15 µM yielded the strongest collateral effect and further increases in concentration probably would only give a minimal improvement, we used 15 µM of protein and guide RNA in all further experiments.

**Figure 1.**
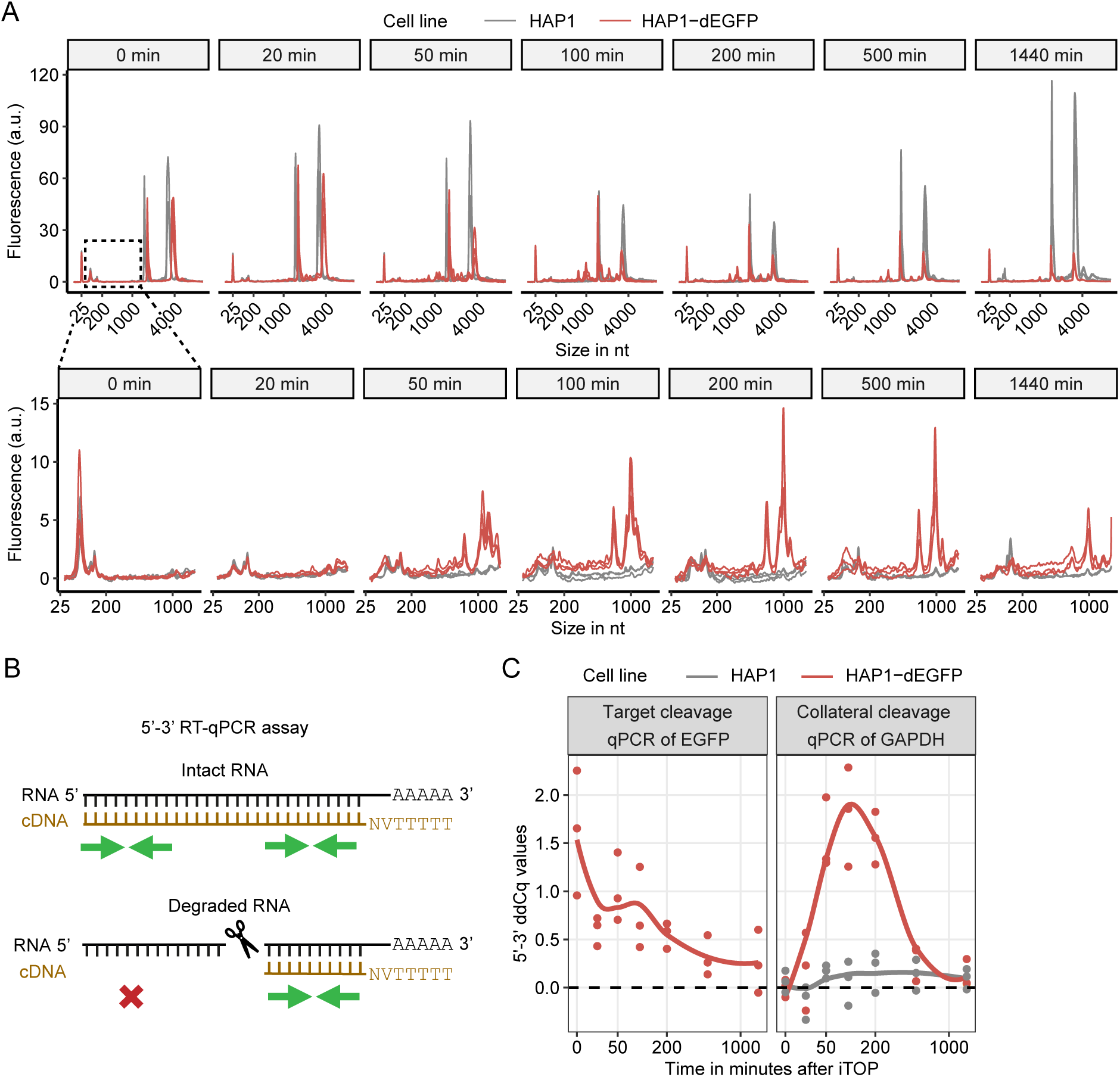
LbuCas13a exhibits strong collateral cleavage activity *in vitro* and in human cells. **A** Total RNA profiles at various timepoints after transfection of the LbuCas13a RNP (n=3). Both HAP1 and HAP1-dEGFP cells were transfected with LbuCas13a and a dEGFP targeting guide RNA. The bottom row is zoomed in to the region indicated by the dotted box in the 0 minute panel. Fluorescence values are corrected for differences in amount of loaded RNA. **B** Schematic depiction of the 5’-3’ RT-qPCR assay. Green arrows indicate the 5’ and 3’ primer pairs used for qPCR. Red cross indicates no amplification is possible when there is a break between the poly-A tail and the 5’ primer pair. RNA is depicted in black, cDNA in brown. **C** 5’-3’ RT-qPCR assay (n=3) on samples from A. Line shows ggplot2’s default LOESS regression (Wickham, 2009). Since wild type HAP1 control cells do not contain dEGFP, no 5’-3’ RT-qPCR assay on dEGFP was performed for these cells.

To specifically assess the cleavage of the target transcript and a collateral transcript, we used a modified RT-qPCR strategy designed to quantify transcript integrity (Auer and Lyianarachchi, 2003; Nolan et al., 2006; Swift et al., 2000), here named 5’-3’ RT-qPCR. In the 5’-3’ RT-qPCR assay, cDNA is created by reverse transcription using oligo-dT primers, driving cDNA synthesis from the 3’ end of the mRNA transcript (Figure 1B). Thus, transcript degradation results in a 3’ bias of the cDNA. Consequently, a 5’ located primer pair is expected to return higher Cq values than a primer pair located at the 3’ end of the transcript. To validate this assay, we first transfected cells with RNase A protein. This indeed resulted in the expected Cq value increase of the 5’ primer pairs relative to the 3’ primer pairs (Supplementary figure 5A). Using the 5’-3’ RT-qPCR assay, we observed that LbuCas13a was able to degrade both target RNA (dEGFP) and collateral RNA (GAPDH) (Figure 1C). While target RNA cleavage seemingly started almost immediately, the collateral GAPDH cleavage started later and peaked at around 100 minutes after transfection. Both target and collateral RNA integrity returned to normal from about 500 minutes after transfection. dEGFP targeting with LwaCas13a, PspCas13b and BzCas13b also resulted in a significant relative increase of the 5’ Cq values of the dEGFP transcript, indicating that these orthologs degraded the dEGFP target RNA in HAP1-dEGFP cells (Supplementary figure 5B). While RfxCas13d did increase the relative 5’ Cq values of the target transcript when targeting dEGFP (Supplementary figure 5C), this increase was not significant (P = 0.076). RspCas13d also did not show significant target RNA cleavage. Although LwaCas13a, BzCas13b and PspCas13b showed a minimal increase in non-target RNA 5’ Cq values compared to the non-targeting control (Supplementary figure 5B), suggesting low levels of collateral cleavage, these increases were not significant. RspCas13d and RfxCas13d did not show an increase of collateral RNA 5’ Cq values (Supplementary figure 5B and C).

Taken together, we observed that under these specific conditions (I.e. iTOP RNP delivery, HAP1 cells, for these specific guide RNAs), LbuCas13a exhibited the highest collateral RNA cleavage activity both *in vitro* and in cells. Therefore, we used LbuCas13a for all further experiments. While collateral cleavage inside cells has also been reported for some of the other orthologs compared here (Ai et al., 2022; Y. Li et al., 2023; Shi et al., 2023), we could not find strong evidence for collateral cleavage with these orthologs. This further highlights the situational nature of collateral cleavage by Cas13 in human cells (Bot et al., 2022), which we further discuss in supplementary note 1.

### Different RNP delivery methods support LbuCas13a collateral cleavage

We next examined whether collateral cleavage could also be induced using different delivery methods, and whether it depended on targeting exogenous RNA. First, we electroporated HAP1 cells with LbuCas13a and a guide RNA targeting one of the highly abundant endogenous transcripts RPS19, GAPDH or 18S rRNA. Delivery by electroporation resulted in strong total RNA degradation for all three guide RNAs, demonstrating that collateral cleavage is not the result of the iTOP delivery method per se, and does not require targeting an overexpressed exogenous transcript (Supplementary figure 6A).

To explore if the collateral RNA cleavage also occurs when LbuCas13a is overexpressed in the cell, we transiently transfected an LbuCas13a expression plasmid into HAP1-dEGFP cells. The transiently transfected cells showed a clear LbuCas13a band on western blot (supplementary figure 7A), despite an average transfection efficiency of only 9.2%. We enriched successfully transfected cells by FACS two days after plasmid transfection. Both the unsorted and enriched cells were transfected with a dEGFP-targeting guide RNA after allowing the enriched cells to recover overnight from sorting. No collateral cleavage pattern was observed in the protein expressing cells transfected with guide RNA (Supplementary figure 7B). Altogether, our data demonstrate that LbuCas13a is capable of collateral cleavage, targeting both exogenous and endogenous transcripts, and using two different RNP delivery methods. However, no collateral cleavage was observed when the LbuCas13a protein was expressed.

### Collateral cleavage occurs in all tested cell lines

Since previous reports have demonstrated that Cas13 activity can be cell-line dependent (Bot et al., 2022), we next explored whether collateral RNA cleavage by LbuCas13a RNP delivery was also cell-type dependent. We used iTOP to transfect the chronic myeloid leukemia derived line HAP1, the rhabdomyosarcoma line RD, the osteosarcoma line U2OS, and the retinal pigment epithelia line ARPE19 with LbuCas13a targeting the highly abundant endogenous transcripts RPS19, GAPDH, or 18S rRNA. All four cell lines demonstrated robust collateral cleavage after targeting any of these target transcripts (Figure 2A). Across all four cell lines, 18S rRNA targeting resulted in the most robust collateral RNA cleavage activity, followed by GAPDH and RPS19 respectively. Notably, the degradation pattern was exactly the same in all conditions and does not depend on the target RNA or cell line. This consistent RNA degradation pattern allowed us to quantify the total RNA degradation by taking the area under the curve of the region where most of the cleaved RNA fragments occur (See Figure 2A). HAP1 cells consistently demonstrated significantly more total RNA degradation than the other three cell lines (Figure 2B). Next, we utilized the 5’-3’ RT-qPCR assay to further quantify the collateral cleavage effect in these samples. RNA degradation was presented by both an overall increase in Cq values compared to untreated cells as well as an increased difference between the 5’ and 3’ primer pair (Figure 2C), with HAP1 again showing the biggest effect. HAP1 cells are known as near haploid cells, however they diploidize quickly in just several passages (Yaguchi et al, 2018). The cells we used in this study are mostly diploid. A possible reason for the higher collateral activity in HAP1 cells could be a higher RNP delivery efficiency, as iTOP was developed using this cell type (D’Astolfo et al, 2015).

**Figure 2.**
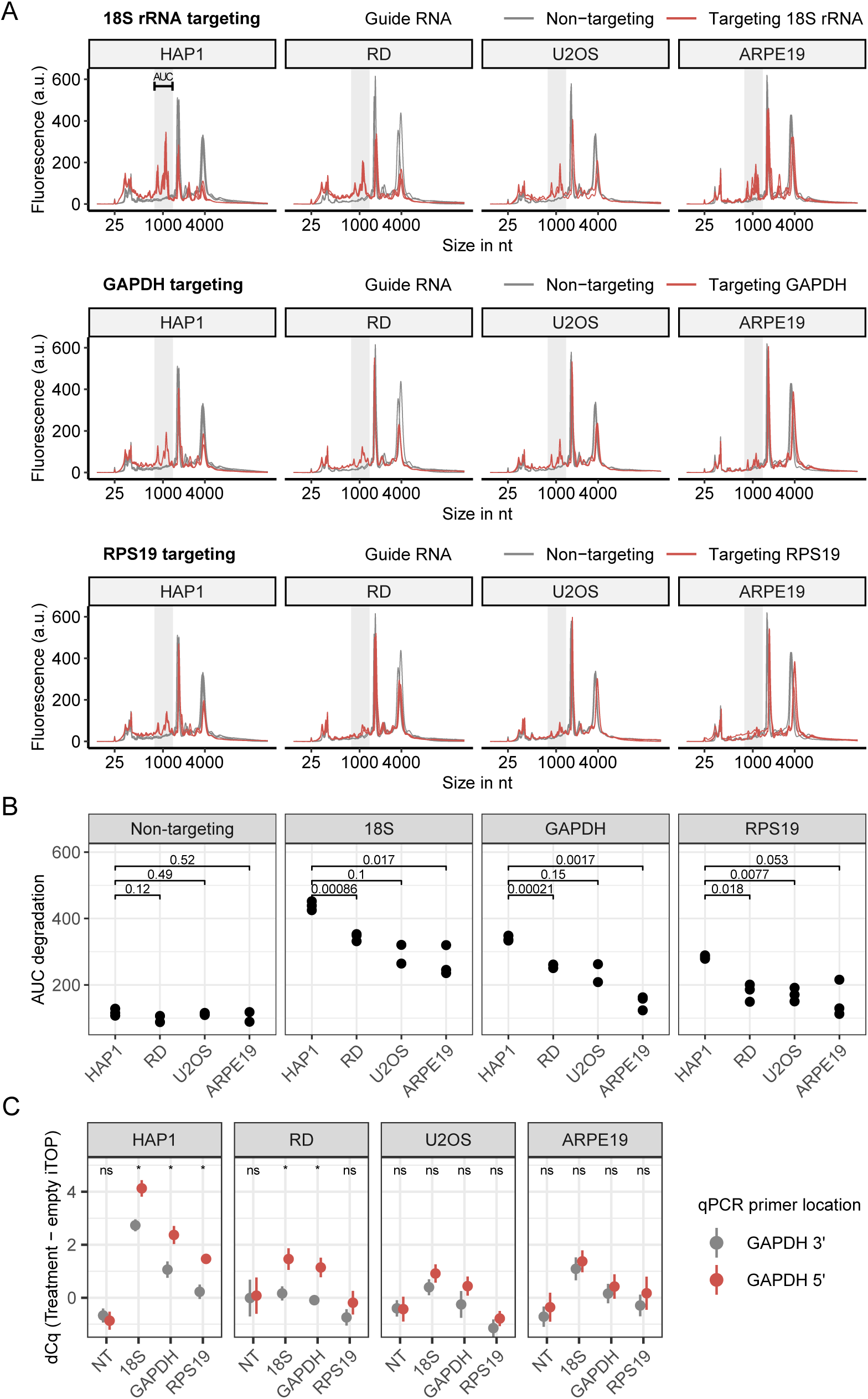
LbuCas13a displays collateral cleavage in different cell lines. **A** Total RNA profiles after targeting the endogenous transcripts 18S rRNA, GAPDH and RPS19 respectively. Total RNA isolated 100 minutes after RNP transfection. Fluorescence values from the Bioanalyzer were normalized to the total area under the curve of each sample. n=3, except for the U2OS 18S rRNA, U2OS GAPDH and ARPE19 non-targeting samples, where n=2. **B** Quantification of total RNA cleavage in different cell lines, by taking the area under the curve of the region outlined by the grey boxes in A. P-values are calculated by unpaired Student’s t-test. **C** 5’-3’ RT-qPCR assays on samples from A (n=3). Dot and line show mean and standard error of the mean. P-values calculated by comparing GAPDH 5’ with GAPDH 3’ using an unpaired Student’s t-test (ns = not significant, * = p-value < 0.05).

Several reports showing specific RNA knockdown with Cas13 used HEK293T cells (Abudayyeh et al., 2017; Cox et al., 2017; Konermann et al., 2018), while others explicitly report low or even the absence of collateral cleavage in this cell type (Ai et al., 2022; Wang et al., 2019), suggesting that Cas13 may be less active in HEK293T cells. Using electroporation to deliver LbuCas13a protein and guide RNA into HEK293T cells, we observed the characteristic RNA degradation pattern upon targeting GAPDH and 18S rRNA (Supplemental figure 6B), indicating that HEK293T cells are not completely void of LbuCas13a collateral cleavage activity. However, in agreement with above mentioned publications, the extent of the collateral cleavage was decreased compared to HAP1 when targeting GAPDH and even undetectable when targeting the highly abundant RPS19 transcript.

### RNA targeting with LbuCas13a induces cell death via apoptosis

Over the course of our experiments we noticed that a day after targeting dEGFP with LbuCas13a cells were rounding up and detaching from the bottom of the plate (Figure 3A), implying loss of viability. To examine this further, we performed live cell imaging with Annexin V as a marker for early apoptosis and CellTox Green to stain for dead cells (Wallberg et al., 2016). When cells die by necrosis they become positive for both these dyes simultaneously. However, if they go into apoptosis Annexin V staining precedes CellTox green staining. We transfected LbuCas13a and one of three different guide RNAs targeting dEGFP into either wildtype or dEGFP expressing cells HAP1. In wildtype cells the number of Annexin V and CellTox green positive cells was similar to the empty transfection control for all three dEGFP targeting guide RNAs (Figure 3B). However, in HAP1-dEGFP the number of Annexin V positive cells rapidly increased from approximately 7.5 hours after transfection, followed by an increase in CellTox Green positive cells approximately 15 hours after transfection (Figure 3B and Supplementary movies 1 and 2). This shows that LbuCas13a is capable of inducing cell death, and that this is specific to cell expressing the target RNA. Furthermore, it demonstrates that the targeted cells go into apoptosis, not necrosis.

**Figure 3.**
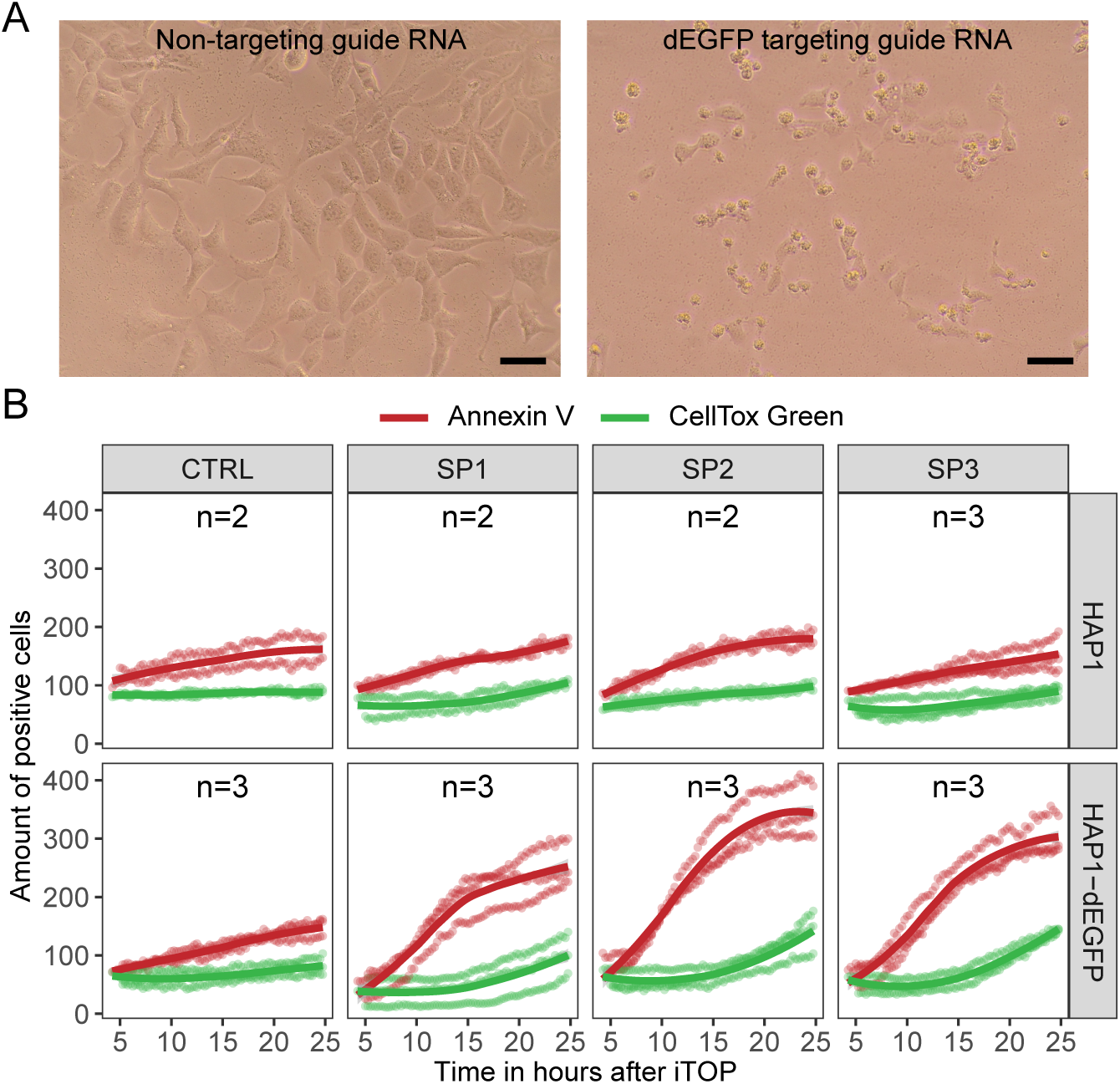
Cells enter apoptosis in response to collateral RNA cleavage. **A** Loss of cell viability when treated with LbuCas13a RNP targeting dEGFP RNA. Images taken 24 hours after transfection. Scale bar is 50 µm. **B** Target RNA specific induction of apoptosis in HAP1 cells by LbuCas13a. Total amount of Annexin V (early apoptosis marker) and CellTox Green (dead cell marker) positive cells during live imaging after transfection. CTRL is an empty transfection. SP1, SP2 and SP3 are transfections with LbuCas13a and three different guide RNAs all targeting dEGFP. Quantified using CellProfiler (McQuin et al., 2018).

### Application of LbuCas13a as a cell selection tool

Given that target-activated collateral RNA cleavage by LbuCas13a can induce cell death, we wondered whether we could exploit this property to eliminate cells that express a specific target RNA from a heterogeneous cell population (Figure 4A). To test this, we mixed fluorescent EGFP expressing HAP1-EGFP cells with wildtype HAP1 cells and transfected this mixed population with LbuCas13a and a guide RNA targeting EGFP. It is important to note that this EGFP targeting guide does not cause collateral RNA cleavage or reduced viability in wildtype cells (Figure 1B and Figure 3B). We then measured the percentage of EGFP positive cells by FACS over time. The percentage EGFP positive cells rapidly decreased when targeting EGFP (Figure 4B), with 36.3% EGFP-positive cells remaining at day 6 compared to more than 60% EGFP positive cells when using a non-targeting guide. The decrease in GFP positive cells was maintained over the entire six-day assay, showing that it was not due to temporary knockdown of the GFP transcript itself, as the GFP transcript integrity is largely restored after 500 minutes to 24 hours after RNP transfection (see Figure 1C). Next, we examined whether multiple sequential selection events could further eliminate the EGFP positive cell population (Figure 4C). We performed several serial transfections at 2-day intervals, with up to 3 subsequent selection rounds. Every additional selection round decreased the percentage of EGFP positive cells in the population by approximately 20 percentage points. From the above data we conclude that Cas13-based cell-selection is able to reduce the target cell population and is repeatable.

**Figure 4.**
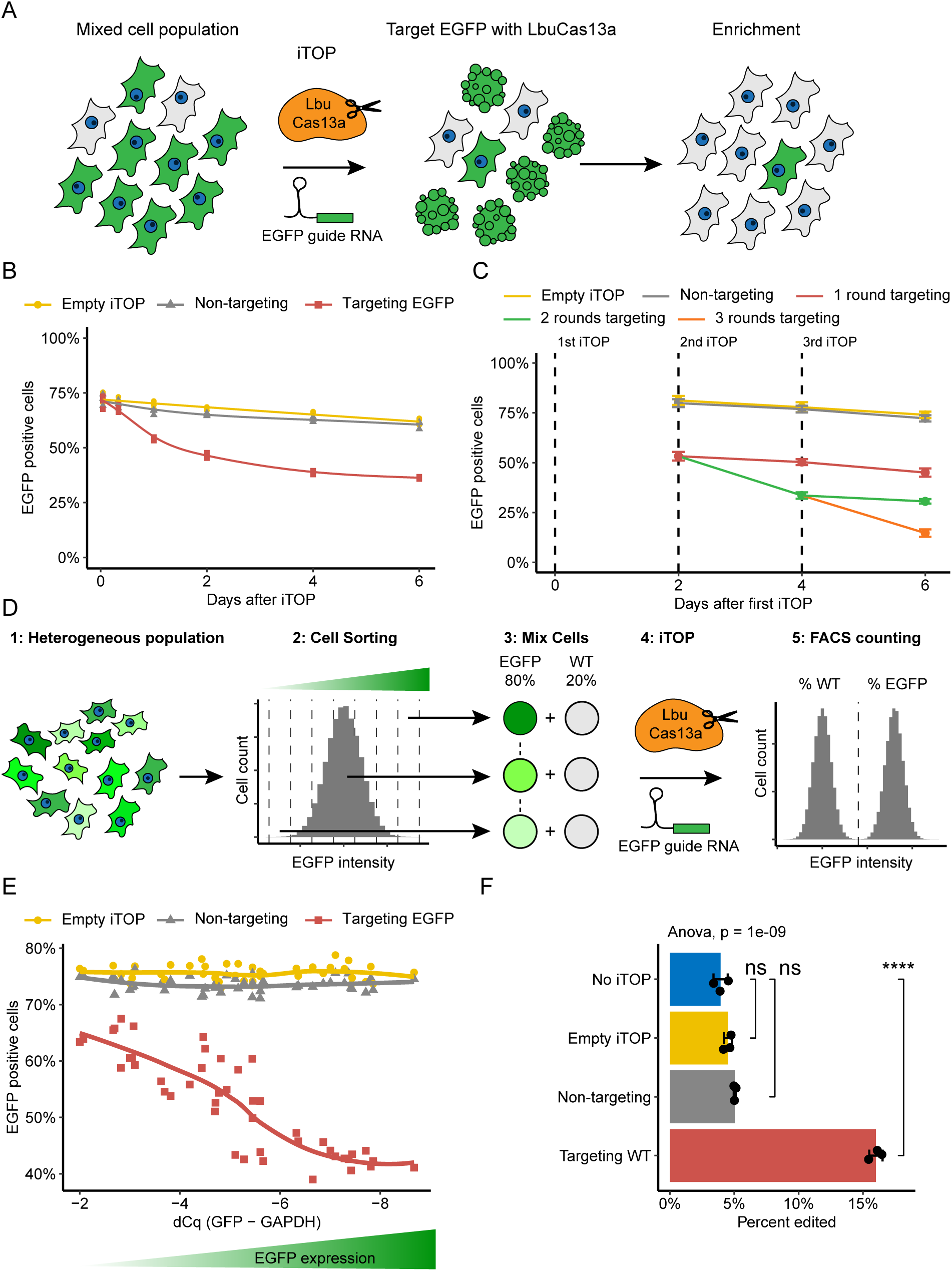
LbuCas13a as a cell selection tool. **A** Schematic of the Cas13-based cell selection. **B** Depletion of EGFP positive HAP1 cells after targeting the EGFP transcript, as measured by FACS (n=3). Trendlines show a generalized additive model using formula y ∼ s(x, k = 6). **C** Multiple rounds of targeting improve depletion of EGFP positive HAP1 cells, as measured by FACS (n=3). Error bars show standard deviation. **D** Schematic showing creation of cell lines with different EGFP expression levels and subsequent selection experiments. **E** Target RNA expression level determines cell selection efficiency (n=3). Target RNA expression level was determined using RT-qPCR, and the percentage of GFP positive cells was determined using FACS. Trendlines created by ggplot2’s default LOESS regression. **F** Enrichment of edited cells by targeting unedited cells with LbuCas13a. Error bars show standard deviation. P-values calculated by unpaired Welch’s t-test (ns = not significant, **** = P-value < 0.0001).

Target RNA expression level theoretically limits the number of active LbuCas13a molecules and therefore the amount of collateral cleavage, which could influence the efficiency of cell selection. To test this, we FACS sorted a heterogeneous population of EGFP expressing cells into cell populations with different levels of EGFP intensity (Figure 4D and Supplementary figure 8A). RT-qPCR confirmed EGFP expression levels increased with EGFP intensity (Supplementary figure 8B). We then tested the depletion efficiency of these cells by mixing them with wildtype HAP1 cells and targeting EGFP with LbuCas13a. In parallel we also performed RT-qPCR on these cell lines to quantify their EGFP expression at the time of the experiment. We found a strong correlation between the expression level of the target RNA and the depletion of EGFP positive cells (Figure 4E), suggesting that activating more LbuCas13a proteins results in more collateral RNA cleavage and a high probability of inducing cell death. However, depletion efficiency seems to plateau at the highest levels of target RNA expression. At this point the RNP delivery efficiency may have become the limiting factor.

We next examined whether the same strategy could be used to positively select for cells that underwent successful gene editing. For these experiments we used a HAP1-EGFP^Δfluor^ cell line containing a 9 nt deletion in EGFP, rendering it non-fluorescent (Supplementary figure 8C) (Zhao et al., 2022). Using this system, we previously reported Cas12a-mediated correction of EGFP fluorescence, with an efficiency of about 5% (Zhao et al., 2022). When we employed LbuCas13a and a guide RNA that targets the unedited cells as a cell selection tool, the percentage of EGFP expressing cells increased to around 15% (Figure 4F). Thus, a single round of selection using LbuCas13a resulted in a 3-fold enrichment. Altogether, these proof-of-principle studies show the potential of Cas13 as a potent cell selection tool, including the enrichment of successfully edited cells.

### Genes involved in the innate immune system are upregulated after collateral RNA cleavage

To gain further insight into the cellular response to the collateral RNA cleavage and the mechanism by which it triggers apoptosis, we performed RNA sequencing at 16 hours after transfection. Based on the 5’-3’ RT-qPCR data (Figure 1C), at this timepoint most mRNA cleavage has been resolved, allowing us to focus on the cellular response following the collateral cleavage. STAR (Dobin et al., 2013) aligned sequences at similar rates across all conditions, however, a higher percentage of the reads from the dEGFP targeting samples mapped to introns compared to the controls, consistent with a relative increase in immature nuclear transcripts generated as a result of cytoplasmic RNA depletion in the cell (Supplementary figure 9A). First, we performed principal component analysis (PCA) (Supplementary figure 9B). The PCA plot shows a clear separation between the dEGFP targeting samples and the controls, suggesting that collateral cleavage by LbuCas13a leads to substantial changes in the transcriptome. Indeed, we found 721 significantly upregulated and 36 significantly downregulated genes (adjusted P-value < 0.05, |Log2 FC| > 1) in response to LbuCas13a activation (Figure 5A). The relatively low number of downregulated genes and high number of upregulated genes is perhaps somewhat counterintuitive in the context of collateral cleavage, but consistent with another report documenting the cellular response to Cas13 collateral cleavage (Y. Li et al., 2023). In agreement with Li et al., we observed that many of the upregulated genes we find are immediate early response genes, such as JUN, FOS, EGR1, EGR2, EGR3, EGR4 and ARC. GO enrichment analysis of the significantly upregulated genes revealed an enrichment of many biological processes related to the innate cellular immune response (Figure 5B), and molecular functions related to chemokine and cytokine activity (Supplementary figure 9C). KEGG pathway analysis also showed enrichment for immune related pathways, such as viral protein interaction with cytokine and cytokine receptor, IL17 signaling pathway, TNF signaling pathway and NF-kappa B signaling pathway (Supplementary figure 9D). These are known factors of the cellular immune response that are triggered, for example, by intracellular recognition of viral RNA and suggests that collateral RNA cleavage by LbuCas13a and subsequent increase in 5’ uncapped RNA and 3’ ends without poly-A tail triggers a similar immune response which ultimately leads to apoptosis.

**Figure 5.**
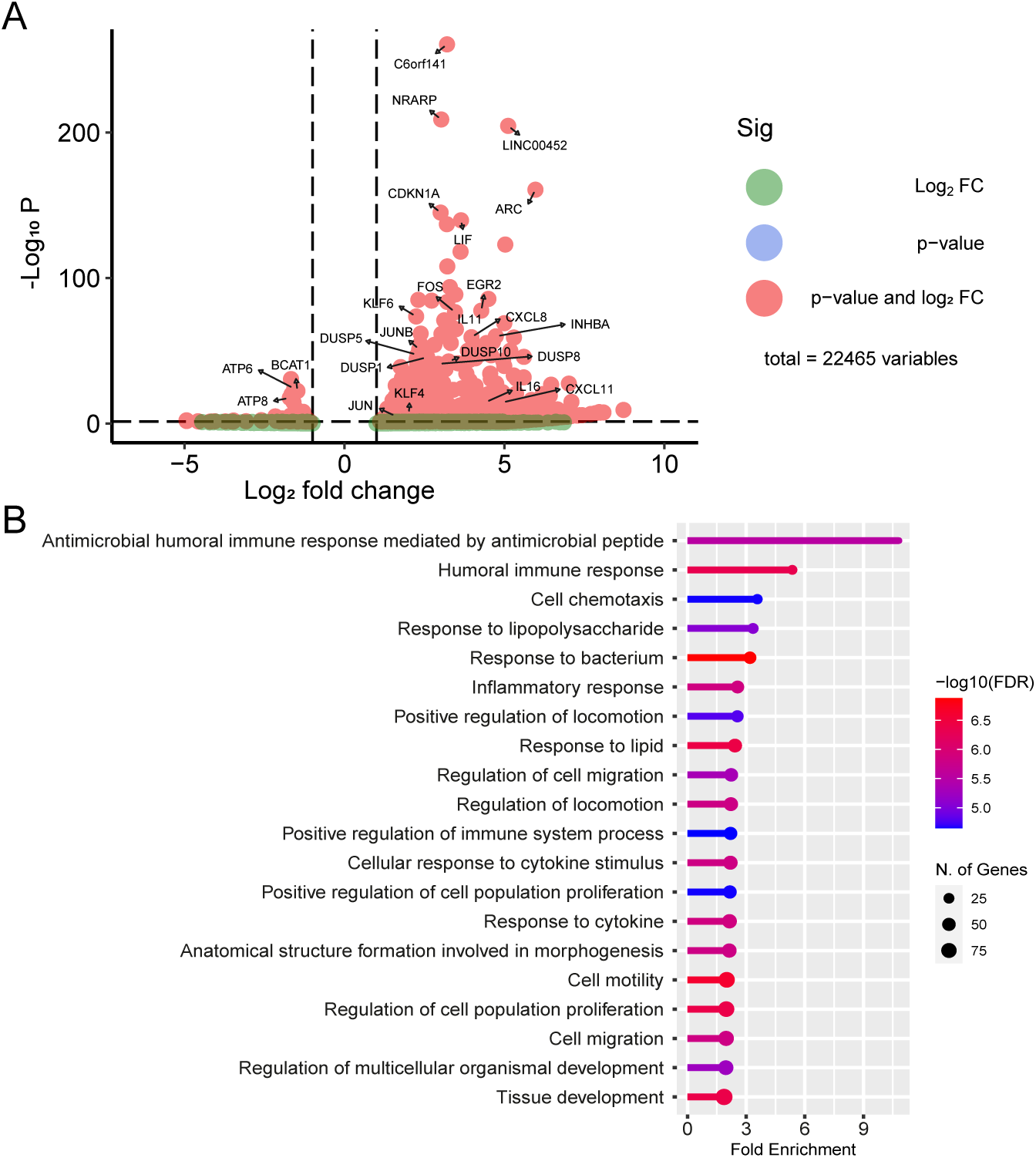
Innate immune response genes are upregulated in response to collateral RNA cleavage. **A** Many genes are upregulated in response to collateral RNA cleavage. RNA-Seq EnhancedVolcano (Blighe et al., 2022) plot of dEGFP targeting samples vs controls, n=3. Fold change and P-values were calculated using DESeq2, see methods. Dotted lines indicate adjusted P-value = 0.05 and |Log2 FC| = 1. **B** A ranked list of enriched GO biological processes using the significantly upregulated genes (adjusted P-value < 0.05 & Log2 FC > 1). Plot generated using ShinyGo (Ge et al., 2020).

### Characterizing collateral RNA cleavage using nanopore sequencing

We decided to use Nanopore sequencing to explore the identity of the RNAs that are subject to collateral cleavage, where RNAs get cleaved by LbuCas13a, and the dynamics of the collateral cleavage process. HAP1-dEGFP cells were transfected with LbuCas13a protein and either a non-targeting or a dEGFP targeting guide RNA. Nanopore read lengths of dEGFP targeting samples were much shorter than control samples (Supplementary figure 10C), containing markedly fewer reads longer than 1000 nt, suggesting widespread collateral RNA cleavage. We found that when targeting dEGFP, the number of protein coding transcripts decreased relative to all other transcripts biotypes (Supplementary figure 11A). On the other hand, the percentage of reads mapping to lncRNAs, snoRNAs, and (in replicate 1) snRNAs increased, suggesting that nuclear RNAs were relatively protected from collateral cleavage. The abundance and read length of reads mapped to protein coding transcripts greatly decreased at 50 and 200 minutes after transfection (Figure 6A and Supplementary figure 11B). At these timepoints, almost all reads were much shorter than the reference they mapped to, whereas most reads in the empty transfection and non-targeting controls were full length transcripts. This suggests unbiased cleavage of all mRNAs after LbuCas13a activation by dEGFP.

**Figure 6.**
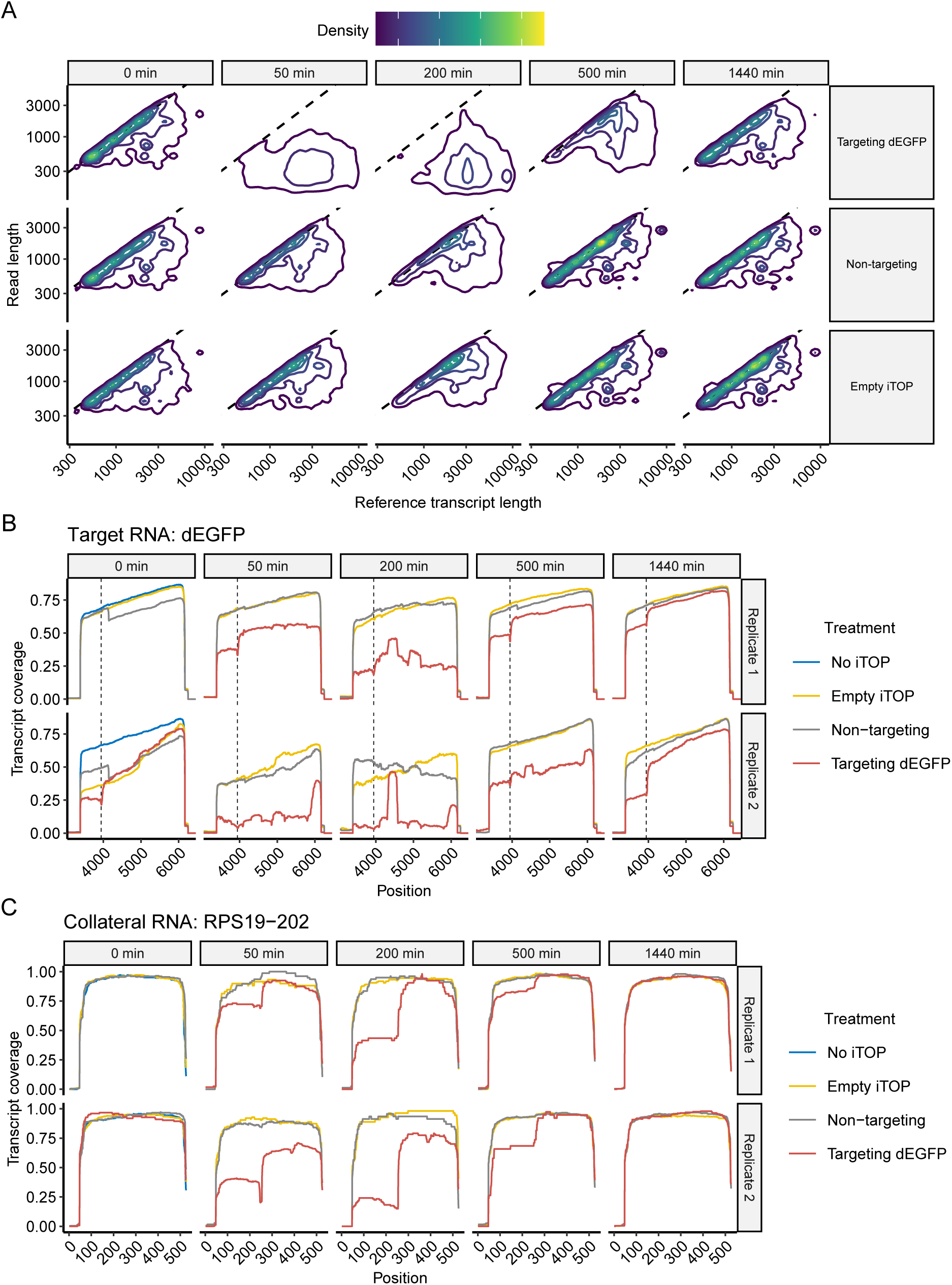
Nanopore sequencing after collateral cleavage by LbuCas13a. **A** Density plot of read length versus reference transcript length of all reads mapped to protein coding transcripts, excluding histone genes and WDR74. Columns show minutes after transfection. Data from replicate 2, see Supplementary figure 5A for replicate 1. **B** Transcript coverage or the target transcript dEGFP. Reads were aligned to the entire plasmid sequence used to make this cell line (Supplementary file 1, pEF-1a_dEGFP-Y66S-IRIS-PuroR-WPRE). The start and stop codon are at position 3415 and 5316 respectively. The dotted lines indicate the protospacer position. Columns show minutes after transfection. The dEGFP targeting sample at 0 minutes after transfection of replicate 1 was excluded due to failed library preparation. **C** Transcript coverage of RPS19-202, a collateral RNA. RPS19-202 appears to have one major cleavage site.

Next we explored the dynamics of cleavage of individual transcripts by calculating their relative transcript coverage. We found that at 50, 200 and 500 minutes after transfection, the overall coverage of dEGFP-IRIS-PuroR is much lower for the dEGFP targeting samples than the control conditions (Figure 6B). Additionally, the target RNA shows a decrease in coverage at the exact location of the protospacer at all timepoints (Figure 6B, dotted lines), which fits with the 5’-3’ RT-qPCR data that showed target RNA cleavage immediately after transfection. This suggest LbuCas13a cleaves in, or very near the protospacer, which is surprising since others have shown that Cas13 cleavage sites do not depend on protospacer position (Abudayyeh et al., 2016; Konermann et al., 2018; Smargon et al., 2017). Alternatively, the protospacer specific dip in coverage may be a technical artifact, for example, due to LbuCas13a still being bound to the target RNA after purification, preventing reverse transcription at this location. Besides the coverage dip around the protospacer, additional dips in coverage were observed, which were consistent between timepoints and replicates. We suspect that these dips are LbuCas13a cleavage sites. Non-target mRNAs and cytoplasmic non-coding RNAs also show these dips in coverage at specific positions, indicating collateral cleavage of these transcripts (Figure 6C, Supplementary figures 12 and 13). We did not observe any differences in coverage between targeting and control samples for small nuclear or mitochondrial RNAs (Supplementary figure 14). The dips in transcript coverage tend to be 10-20 nucleotides wide, so we cannot discern the exact cleavage position (Figure 7A). *In vitro*, LbuCas13a was shown to prefer cleaving at uracil residues in or near single stranded loop structures (East-Seletsky et al., 2017). In our data, transcript coverage dips often map to predicted stem-loop structures, with uracil containing single stranded loops on the 5’ side of the coverage dip (Figure 7B), suggesting that the 5’ side of the dip is the actual cleavage location. These results indicate that LbuCas13a cleavage preferences follow the same pattern in human cells as observed *in vitro*. Next, we manually annotated 100 potential cleavage sites (Supplementary table 4), assuming that the Uracil closest to the 5’ side of the coverage dip was the cleavage position. This revealed a US (S = G or C) di-nucleotide preference (Figure 7C). Taken together, the Nanopore sequencing provides further evidence of collateral RNA cleavage in human cells by LbuCas13a and suggests that both mRNAs and cytoplasmic ncRNAs are cleaved without bias. Additionally, LbuCas13a cleaves RNA at specific positions, showing a preference for uracil-containing single stranded loop structures.

**Figure 7.**
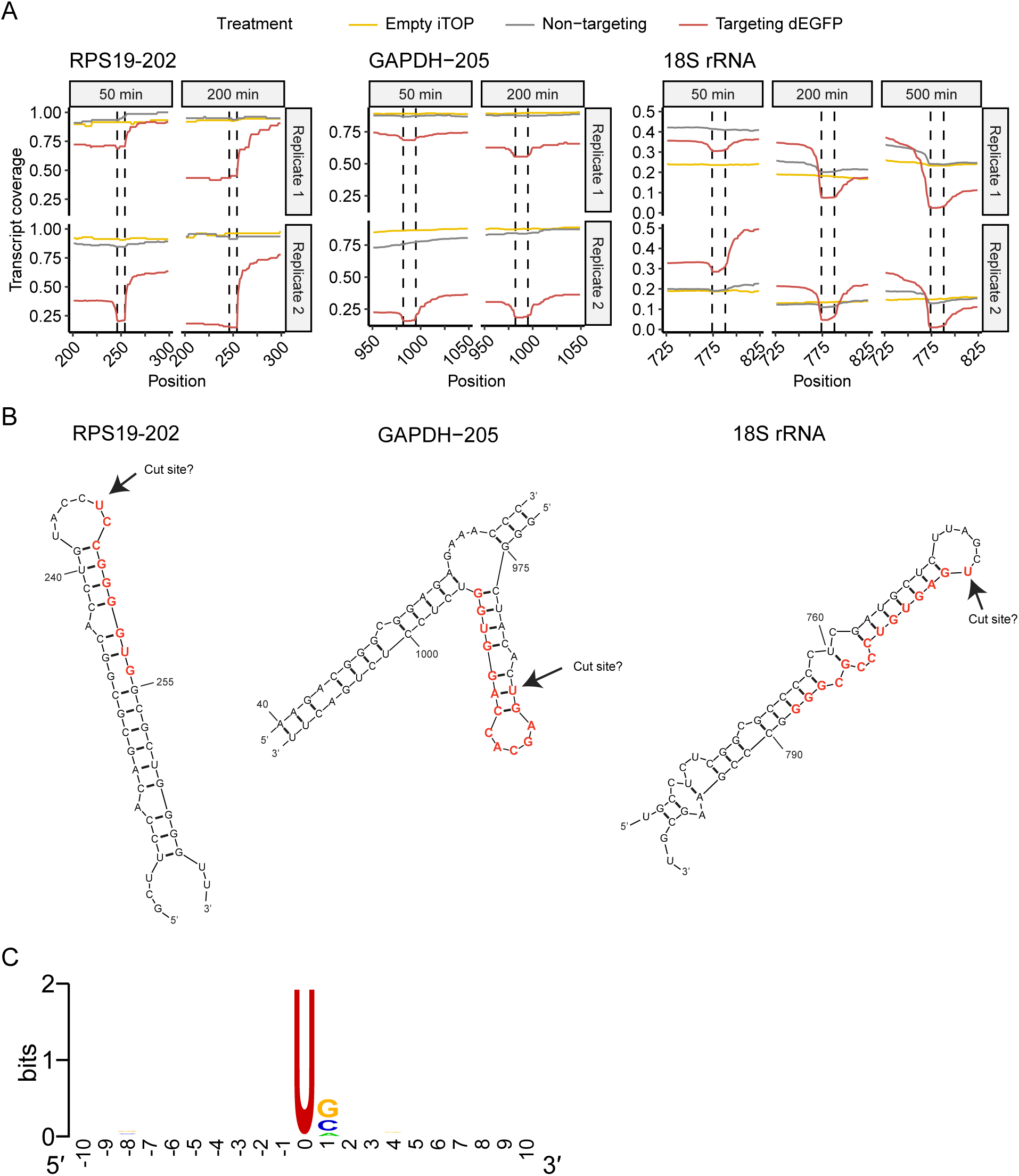
LbuCas13a collateral RNA cleavage sites in human cells. **A** Transcript coverage of non-target RNAs, zoomed in at putative cleavage sites. **B** The predicted RNA structures at the putative cleavage sites from A. Red letters are the approximate location of the coverage dips, corresponding to the dashed lines in A. RNA structures for RPS19-202 and GAPDH-205 are the top predictions of Mfold (Zuker, 2003) using the default setting. 18S rRNA structure provided by RNAcentral.org (Consortium, 2021) (URS0000726FAB_9606). **C** Sequence logo around 100 manually annotated cleavage sites. The U at position 0 was enforced during potential cleavage site selection. Logo made using WebLogo (Crooks et al., 2004).

## Discussion

In this work, we revealed the temporal dynamics of both target and collateral RNA cleavage by LbuCas13a, as well as the cellular responses to this collateral RNA cleavage (Figure 8). Target and collateral RNA cleavage start within 50 minutes after RNP delivery, with target cleavage preceding collateral RNA cleavage. All cytoplasmic RNAs seems to be cleaved in an unbiased manner, with a cleavage preference for Uracil containing stem loop structures. Target and collateral mRNA integrity mostly returns to normal at around 500 minutes after RNP delivery, while ribosomal RNA takes longer to recover, with some cleavage fragments still being present at 24 hours after delivery. The recovery of mRNA integrity coincides with the cells starting to become positive for Annexin V, indicating early apoptosis, at about 6-8 hours after delivery. Stress and innate immune response genes are upregulated, probably contributing to the induction of apoptosis. From about 15 hours after delivery, cell death starts occurring. As such, LbuCas13a can be used as a negative cell selection tool.

**Figure 8.**
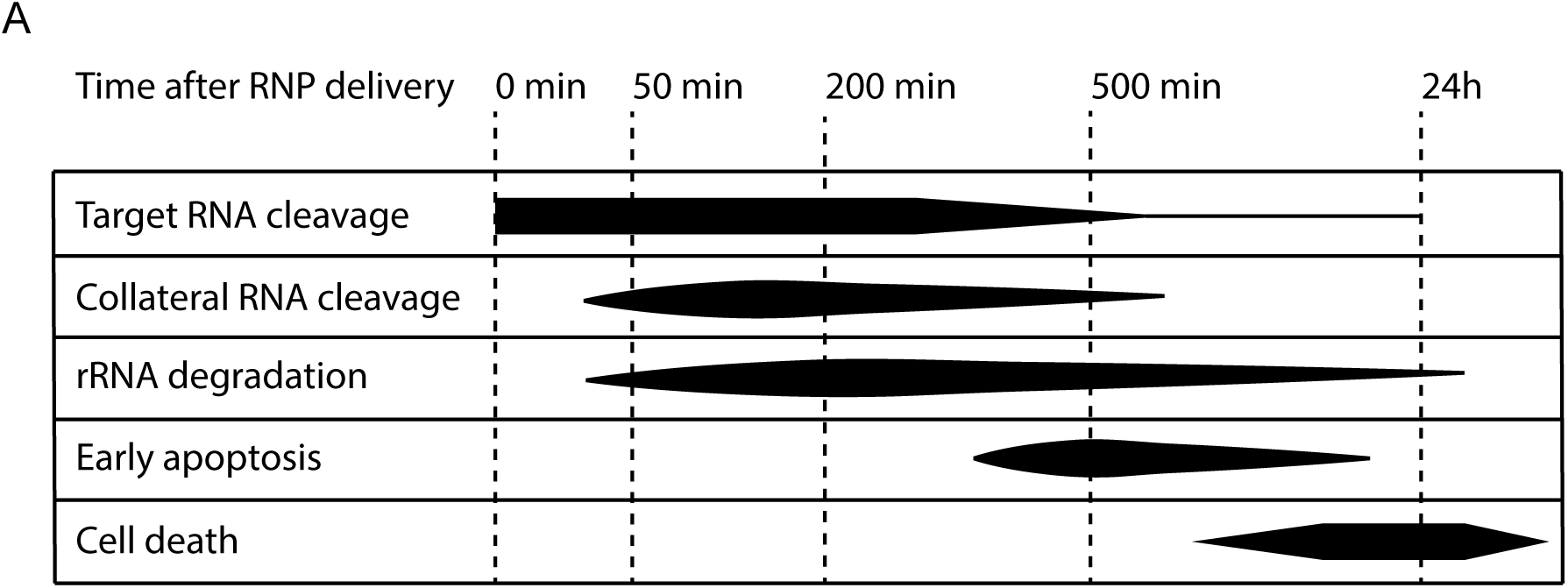
A timeline of events after RNA targeting with LbuCas13a in cells. **A** This plot provides an summary of the timing of events after RNA targeting with LbuCas13a. The target RNA cleavage timing takes into account both figures 1C and 6B. While figure 1C suggests target cleavage peaks at 0 minutes after delivery, figure 6B suggests it peaks at 50-200 minutes after delivery. Collateral RNA cleavage timing is based on figures 1C, 6A and 6C. For ribosomal RNA degradation see figure 1A. The early apoptosis timeline is based on figure 3B. The cell death timeline is based on figure 3B and 4B.

The collateral RNA cleavage activity of Cas13 is well established *in vitro* (Abudayyeh et al., 2016; East-Seletsky et al., 2017, 2016; Konermann et al., 2018; Smargon et al., 2017; Xu et al., 2021; Yan et al., 2018) and in bacteria (Abudayyeh et al., 2016; Meeske et al., 2019; Smargon et al., 2017; Yan et al., 2018), however in eukaryotes the situation is less clear. While many groups report that no collateral cleavage seems to occur (Abudayyeh et al., 2017; Cox et al., 2017; Konermann et al., 2018; Tieu et al., 2024; Wessels et al., 2024), others have recently run into unexpected cytotoxicity and/or apparent collateral RNA cleavage when using CRISPR-Cas13 in eukaryotic cells (Ai et al., 2022; Buchman et al., 2020; Y. Li et al., 2023; Özcan et al., 2021; Shi et al., 2021; Tong et al., 2021; Wang et al., 2019). From these reports and our work, several factors have emerged that are likely important in determining if and how much collateral cleavage occurs, namely the collateral activity of the ortholog, the intracellular RNP concentration, the target RNA expression level and cell line specific differences (Bot et al., 2022).

When comparing the collateral activity of several orthologs *in vitro*, in our hands LbuCas13a was the most active ortholog. However, different orthologs may have different preferences for factors such as the assay buffer, the protospacer sequence (Wessels et al., 2024, 2020) and the cleavage substrate (East-Seletsky et al., 2017). When comparing the collateral RNA cleavage activities of these Cas13 orthologs in cells, only LbuCas13a exhibited robust collateral cleavage across a range of different guide RNAs. While other reports have documented collateral cleavage or toxicity when using LwaCas13a (Özcan et al., 2021; Wang et al., 2019), or RfxCas13d (Ai et al., 2022; Y. Li et al., 2023; Shi et al., 2023), we did not find collateral RNA cleavage activity in human cells with these orthologs (see supplementary note 1).

We detected collateral cleavage when delivering LbuCas13a as RNP, but not when LbuCas13a was expressed as a transgene. Cas13 expression and protein levels may be limited by guide RNA independent RNA cleavage activity of Cas13 (Z. Li et al., 2023), potentially explaining why only RNP delivery allowed us to detect collateral RNA cleavage. Here we showed that by combining the highly active LbuCas13a with RNP delivery, apoptosis can be induced in target RNA expressing cells. This allowed us to specifically deplete target cells from a mixed population. With just one round of selection, we reach a similar efficiency as Shi and colleagues (Shi et al., 2023). However, unlike Shi et al., we show that multiple selection rounds are possible, achieving much greater negative selection efficiency, namely reducing target cells from 80% to just 13% of the population. In addition, Shi and colleagues use cells stably expressing RfxCas13d, which requires the generation of such an RfxCas13s expression cell line before cell selection is possible. By transfecting Cas13 as RNP, our system can be immediately applied to any cell type without the need for introducing any transgenes. Finally, to our knowledge we are the first to show that Cas13 can be applied to enrich for cells that have undergone a rare gene editing event, achieving a three times enrichment. Enriching for edited cells with CRISPR-Cas13 has major advantages over other methods that select for genome editing events, as it does not require the co-insertion of a selection marker (Shy et al., 2016) or a second selectable edit (Ewen-Campen and Perrimon, 2018). Furthermore, it does not depend on co-expression of gene editing components with a selection marker, which risks selection for permanent genomic integration of the gene editing plasmid.

Although we were able to deplete cells from a mixed population, we were not able to completely eliminate all target cells. In addition, high target RNA expression is required to enable the induction of cell death, which limits the potential target RNAs that can be used. Nevertheless, we are optimistic that future iterations of the CRISPR-Cas13 cell selection system will overcome these limitations. There are several potential strategies to improve the CRISPR-Cas13 cell selection systems. For example, the number of selection rounds could be increased, since we observed no indication of reduced efficiency with subsequent selection rounds. Furthermore, the sensitivity to collateral cleavage of cells might differ between different phases of the cell cycle, as it does for many anti-cancer drugs (Johnson et al., 2021), so cell cycle synchronization may be a factor to consider as well. Finally, orthologs with even higher collateral RNA cleavage activity than LbuCas13a could be discovered or engineered through directed protein evolution approaches.

In conclusion, here we demonstrated that LbuCas13a is capable of high collateral cleavage activity in human cell lines. We described in detail the events occurring following activation of LbuCas13a by a target RNA. Finally, we showed how CRISPR-Cas13 can be employed as a specific and effective cell elimination tool. We envision that this new system will find broad application in both basic and clinical research, enabling the selective elimination of unedited cells, specific cell types to improve stem cell differentiation yields or perhaps the specific elimination of cancer cells.

## Materials and Methods

### Contact for reagent and resource sharing

Recombinant LbuCas13a and AsCas12a proteins, the HAP1-dEGFPY66S and the HAP1-dEGFPΔfluor cell lines are available through Divvly (https://divvly.com/geijsenlab).

### Molecular cloning

For recombinant protein production, all Cas13 orthologs were cloned into a pET15b expression vector (Addgene: 62731). LbuCas13a and LshCas13a were a gift from Jennifer Doudna (Addgene: 83482 and 83487). LwaCas13a, BzCas13b and PspCas13b were a gift from Feng Zhang (Addgene: 91924, 89898 and 103862). RspCas13d was a gift from Arbor Biotechnologies (Addgene: 108304). pET-28b-RfxCas13d-His was a gift from Ariel Bazzini & Miguel Angel Moreno-Mateos (Addgene: 141322). Cloning was done using standard molecular biology techniques using In-Fusion cloning according to the manufacturer’s instruction. For the construction of the lentiviral LbuCas13a and RfxCas13d protein and guide expression vectors, all fragments were ordered from IDT as gBlocks. The fragments were cloned into the pEF-1a_dEGFP^Y66S^-IRIS-PuroR-WPRE lentiviral plasmid backbone. Cloning was done using standard molecular biology techniques using In-Fusion cloning according to the manufacturer’s instruction. All plasmid sequences are available in supplementary file 1. Plasmids will be made available on Addgene.

### Cell culture

HAP1 cells were a gift from Dr. Brummelkamp, NKI Amsterdam. RD cells were a gift from Dr. Gerben Schaaf, Erasmus MC. HEK293T cells were ordered from Takara (#632180). U2OS and ARPE19 cells were ordered from ATCC (HTB-96 and CRL-2302 respectively). All cell lines were tested for mycoplasma monthly.

HAP1 cells were cultured in Iscove’s Modified Dulbecco’s Medium (IMDM) supplemented with 10% Fetal Bovine Serum (FBS). HEK293T cells were cultured in Dulbecco’s Modified Eagle Medium (DMEM) with 10% FBS. U2OS and RD cells were cultured in DMEM supplemented with 10% FBS and 2 mM L-Glutamine. ARPE19 cells were cultured in DMEM/F12 with 10% FBS. All culture medium was purchased from Gibco. All cells were grown at 37°C in a humidified atmosphere containing 5% CO_2_.

To generate the HAP1-dEGFP^Y66S^, HAP1-dEGPF^△fluor^ and HAP1-EGPF lines (Zhao et al., 2022), HAP1 cells were transduced with the pEF-1a_dEGFP^Y66S^-IRIS-PuroR-WPRE, pEF-1a_EGFP^△fluor^-IRIS-PuroR-WPRE or pEF-1a_EGFP-IRIS-PuroR-WPRE plasmids (Supplementary file 1) using lentivirus. Two days after transduction, puromycin selection was started at 0.5 µg/mL to select for integration of the plasmid. After resistant cells had established themselves, the puromycin concentration was increased every 2 days to 1, 2, 5 and finally 10 µg/mL. HAP1-dEGFP^Y66S^, HAP1-dEGFP^△fluor^ and HAP1-EGFP cells were maintained in HAP1 medium supplemented with 10 µg/mL puromycin.

To generate the LbuCas13a and RfxCas13d protein or guide RNA expressing cells, HAP1-dEGFP cells were transduced with lentivirus (See Supplementary file 1 for plasmid sequences). Two days after transduction, blasticidin selection was started at 10 µg/mL. Because the percentage of BFP positive cells remained low, cells were sorted for being BFP or mScarlet positive at 14 after infection on a CytoFLEX SRT benchtop cell sorter (Beckman Coulter). Cells were maintained in HAP1 medium supplemented with 10 µg/mL blasticidin.

### Lentivirus production

Lentivirus was produced in HEK293T cells. Briefly, 4 packaging plasmids pHDM-Hgpm2, pHDM-Rev1b, pHDM-Tat1b, pHDM-G and the transfer plasmid (in the ratio of 2:1:1:2:4) were transfected to HEK293T cells with FuGENE HD transfection reagent (Promega, E2311) according to the manufacturer’s protocol. The FuGENE to DNA ratio was 2.5:1. One day after transfection, the cell culture medium was supplemented with medium containing caffeine (Sigma C0750) and HEPES (Gibco 15630-080), to a final concentration of 3 mM and 25 mM respectively. The supernatant containing the lentivirus was harvested 48 hours after transfection, filtered with 0.22 µm filter (EMD Millipore, SCGP00525), and stored at 4°C.

### Expression and purification of recombinant Cas13 proteins

The sequence of the expression vectors is available in supplementary file 1. Expression vectors will be made available on Addgene. The different recombinant Cas13 orthologs were produced as described previously for Cas12 (Zhao et al., 2022). In brief, the proteins were expressed in *E. coli* BL21(DE3) and purified by Ni-NTA affinity purification followed by gel-filtration chromatography. The purified Cas13 proteins were concentrated to 75 μM using the Amicon ultra centrifugal filter MWCO 50 kDa (Merck), flash frozen in liquid nitrogen, and stored at -80°C.

### Preparation of guide RNAs and target RNA by in vitro transcription

Oligonucleotides carrying the T7 promoter and appropriate downstream sequence were synthesized by IDT and PCR-amplified to generate *in vitro* transcription (IVT) templates (see supplementary table 1 for template and primer sequences). The EF-1a_dEGFP^Y66S^-IRIS-PuroR-WPRE plasmid (supplementary file 1) was used as PCR template to generate the IVT template for the target RNA. Transcription reactions were performed at 37°C for 4 hours in a buffer containing 40 mM Tris-HCl (pH 8), 20 mM MgCl_2_, 5 mM DTT, 2 mM NTPs, 50 μg/mL T7 RNA polymerase, and 250 ng/μL DNA template. Next, to remove the DNA template, 1 μL Turbo DNase (Invitrogen: AM2238) per 100 μL IVT reaction was added and incubated at 37°C for 30 minutes. The RNA was then purified using the Zymo RNA Clean & Concentrator -25 kit (R1017). Typically, four different guide RNAs were tested for each target RNA, assessing collateral cleavage by comparing total RNA degradation using the Bioanalyzer (see below). The guide RNA with the highest amount of total RNA degradation was selected for use in experiments.

### In vitro collateral cleavage activity assay

Collateral RNA cleavage detection assays were performed with IDT RNaseAlert (IDT, 11-04-02-03) according to the manufacturer’s protocol. Reactions were performed in different cleavage buffers depending on the Cas13 ortholog. For LshCas13a, LwaCas13a, LbuCas13a and RfxCas13d the buffer (Abudayyeh et al., 2016) consisted of 40 mM Tris-HCl pH 7.3, 60 mM NaCl and 6 mM MgCl_2_. For Cas13b (BzCas13b and PspCas13b), the cleavage buffer contained 10 mM Tris-HCl pH 7.5, 50 mM NaCl and 0.5 mM MgCl_2_ (Smargon et al., 2017). For RspCas13d, the cleavage buffer used was 20 mM HEPES pH 7.1, 50 mM KCl, 5 mM MgCl_2_ and 5% glycerol (Yan et al., 2018). Before the RNaseAlert assay, 1 μM Cas13 and 500 nM guide RNA were preassembled in 10X cleavage buffer for 10 min at 37°C. The RNaseAlert assays were performed using 100 nM Cas13 protein and 50 nM guide RNA with the appropriate amount of target RNA, using a 50 µL reaction volume. Reactions were prepared on ice, in opaque 96-well plates (Greiner, 655076). Fluorescence was measured every 5 minutes using a Tecan Spark plate reader, while incubating at 37°C. Reactions were allowed to proceed for at least one hour. The collateral cleavage rate was calculated in Excel by applying the SLOPE function to the initial linear increase in fluorescence.

### Ribonucleoprotein transfection using iTOP

The Cas13 proteins and their guide RNAs were transfected into target cells using iTOP, using a protocol adapted from D’Astolfo et al. (D’Astolfo et al., 2015). The iTOP protocol consists of four steps: (1) seeding the cells, (2) iTOP transfection mixture preparation, (3) incubating the cells in the iTOP mixture, and (4) recovery. First, cells were plated the day before transfection in Matrigel coated pates (Corning: 356231, diluted 1:100 in PBS), such that the confluency was ∼70% the next day. Second, to prepare the iTOP transfection mixture, 2/5 of transfection supplement B (Opti-MEM media supplemented with 810 mM NaCl, 504 mM GABA, 1.875X concentrated N2 (Gibco: 17502048), 1.875X concentrated B27 (Gibco: 17504044), 1.875X concentrated MEM non-essential amino acids (Gibco: 11140050), 3.75 mM L-Glutamine (Gibco: 25030081), 2.5 µg/mL bFGF (Gibco: PHG0023), and 0.625 µg/mL EGF (Gibco: PHG0313)), 1/5 of Cas13 protein (75 µM) or empty protein storage buffer (Zhao et al., 2022), 1/5 of guide RNA (75 µM) or nuclease-free water, and finally 1/5 nuclease-free water were combined. iTOP mixture volumes used per well were: 50 µL for 96-well plates, 150 µL for 48-well plates, 200 µL for µ-slide 8-well (ibidi, 80827) and 1000 µL for 6-well plates. Third, the culture medium was aspirated and iTOP mixture was carefully added into the wells. Next the cells were returned to the cell culture incubator for 45 minutes. Fourth, immediately after incubation the iTOP mixture was aspirated and replaced by pre-warmed cell culture medium (96-well: 200 µL, 48-well: 600 µL, µ-slide 8-well: 300 µL, 6-well: 4000 µL). Finally cells were returned to the cell culture incubator or immediately harvested for downstream analysis.

### Ribonucleoprotein transfection using electroporation

A Lonza Nucleofector system including a Nucleofector 2b Device and the Cell Line Nucleofector Kit V (Lonza) were used for electroporation of HAP1-dEGFP and HEK293T cells targeting endogenous transcripts (Supplementary figure 6A and 6B), according to the manufactures instructions. One million cells were used per electroporation, resuspended in 100 µL Nucleofector solution V buffer containing 1 µM LbuCas13a and 1 µM guide RNA. The electroporation was performed in the Nucleofector 2b device using the ‘Cell-line T-020’ program. For the comparison of LbuCas13a with RfxCas13d (Supplementary figure 3 and 5C), HAP1-dEGFP cells were electroporated using a Bio-Rad Gene Pulser Xcell. One million cells were resuspended in Gene Pulser electroporation buffer (Bio-Rad #1652677) containing 1 µM Cas13 protein and 1 µM guide RNA in a total volume of 150 µl. Cells were electroporated in 4mm cuvettes using a single 400V, 5ms square wave. After electroporation, the cells were incubated at 37°C.

### Plasmid transfection

Cells were plated in 6-well plates such that the confluence was ∼80% at the time of transfection. Five µg plasmid DNA was mixed with 15 µg PEI (MW 25000, Polysciences #23966) in a total volume of 500 µl opti-MEM (Gibco #31985070). The mixture was incubated at room temperature for twenty minutes. After incubation, the mixture was added dropwise to the culture medium in the well. Next, the 6-well plate was spun at 200g for five minutes, after which the cells were return to the incubator.

### Total RNA isolation

Cells were collected in TRIzol reagent (Invitrogen: 15596018) and stored at -80°C until RNA isolation. To isolate total RNA, chloroform was added to the TRIzol at a ratio of 1:5 and samples were mixed by briefly vortexing. Next, samples were centrifuged for 15 minutes at 12000 g at 4°C. Total RNA was isolated from the colorless upper aqueous phase using the Zymo RNA Clean & Concentrator -5 kit, including an in-column DNase treatment, according to the manufacturer’s instructions.

### Bioanalyzer

Total RNA integrity was analyzed on the Bioanalyzer using either the RNA nano or RNA pico chips, depending on the available RNA amount. The area under the curve (AUC) was calculated in R with the “AUC” function from the DescTools package using the default settings (Signorell, 2021). To account for differences in the amount of RNA loaded, all samples were normalized to their total AUC. This was done by dividing the sample fluorescence by the sample AUC and multiplying by the mean AUC across all samples. In our hands, this method resulted in a more accurate normalization than using the sample concentration estimates supplied by the Bioanalyzer software.

### 5’-3’ RT-qPCR

RNP transfection by iTOP was performed in 96-well plates, as described above. Besides LbuCas13a and either a targeting or a non-targeting guide RNA, an empty transfection control was always included. All ddCq values are relative to this empty transfection control. Total RNA was isolated and DNase-treated as described above. An anchored oligo(dT) primer was used for reverse transcription using SuperScript III reverse transcriptase (Invitrogen, 18080093), according to the manufacturer’s protocol. qPCR was performed in triplicate using Bio-Rad iQ SYBR Green Supermix with 300 nM primers. The primer sequences used for reverse transcription and qPCR can be found in supplementary table 2.

Traditionally, the ratio between the Cq values of the 3’ and 5’ primer pair has been used to quantify transcript integrity (Nolan et al., 2006). However, this will results in different ratios for the same amount of degradation when expression levels are different (^20^ ≠ ^25^). Therefore, to quantify the transcript integrity we used a modified version of the ddCq method where the 3’ primer pair takes on the function of the housekeeping gene. First, to calculate the 5’-3’ dCq values, the 3’ Cq value was subtracted from the 5’ Cq value for each sample. Next, to calculate the 5’-3’ ddCq, the mean 5’-3’ dCq value of the empty transfection controls was subtracted from the 5’-3’ dCq values of the treatments. To avoid confusion with the method were the ratio between the 3’ and 5’ primer pair is used, we called the assay 5’-3’ RT-qPCR here instead.

### Western blot

First, the cells were put on ice and washed three times with ice cold PBS. Cells were collected in ice cold RIPA buffer and kept on ice for 20 minutes. The lysate was spun down at 16000g at 4°C for 20 minutes. The supernatant was transferred to a new tube and the protein concentration was measured using a BCA assay (Thermo Scientific, 23225). Before loading on gel, samples were mixed with Laemmli buffer (Bio-Rad, 1610747) and heat denatured at 95°C for 5 minutes. Twenty µg protein lysate was loaded per well, unless otherwise indicated.

Samples were run on 4-20% Mini-PROTEAN precast gels (Bio-Rad, 4568094). The protein was then transferred to a PVDF membrane using a wet transfer at 300 mA for 90 minutes. After transfer, the membrane was blocked in blocking buffer (PBS with 5% w/v skim milk powder, 1% w/v BSA, and 0.1% Tween20) at room temperature for twenty minutes. Next, the membrane was stained overnight in blocking buffer with an anti-HA tag (ThermoFisher, 26183) and a GAPDH loading control antibody (ThermoFisher, MA5-15738), diluted at 10000x and 1000x respectively. Next, the membrane was washed three times for five minutes in PBST (PBS with 0.1% Tween20). The membrane was stained for two hours at room temperature with Goat Anti-Mouse IgG H&L (HRP) (Abcam, ab6789), diluted 1:1000 in blocking buffer. After staining the membrane was again washed three times for five minutes in PBST. Finally, ECL substrate (ThermoFisher, 32106) was applied to the membrane and the signal was detected on a Bio-Rad GelDoc Go System.

### Nanopore cDNA sequencing

The experiment and subsequent library preparation consisted of seven steps: (1) RNP transfection, (2) RNA isolation, (3) rRNA depletion, (4) poly(A) tailing, (5) reverse transcription, (6) second strand synthesis and (7) barcode and sequencing adapter ligation. Manufacturer’s instructions were followed for all kits unless otherwise noted.

To ensure enough RNA for sequencing, this experiment was performed in 6-well plates, plating 600 000 HAP1-dEGFP^Y66S^ cells per well the day before the transfections. One well was reserved as a no transfection control. iTOP was performed as described above, using an iTOP mixture volume of 1 mL per well. Each timepoint had three conditions: an empty transfection control, a transfection with LbuCas13a and a non-targeting guide RNA, and a transfection with LbuCas13a and a dEGFP targeting guide RNA (SP2). Cells were collected in TRIzol at 0, 50, 100, 200, 400 and 1440 minutes after iTOP. Total RNA was isolated and DNase-treated as described above. RNA integrity was assessed on the Bioanalyzer (Supplementary figure 10A). Next, 5 µg total RNA of each sample was used for rRNA depletion with the RiboMinus Eukaryotic system v2 (Invitrogen, A15026). rRNA depletion was confirmed with the Bioanalyzer (Supplementary figure 10B). In order to also detect cleaved RNA fragments without a poly-A tail, we polyadenylated the rRNA depleted RNA with *E. coli* Poly(A) polymerase (NEB, M0276L) in the presence of 1 mM ATP and 1 unit/uL RNasin® (Promega, N2515) for 30 minutes at 37°C. Afterwards, the RNA was cleaned with the Zymo RNA Clean & Concentrator -5 Kit (R1013). Next, reverse transcription was performed using SuperScript IV RT (Invitrogen, 18090010) and anchored oligo dT_20_ primers (Supplementary table 2). The reverse transcription reaction was incubated at 50°C for 15 minutes followed by 80°C for 10 minutes. To remove RNA before second strand synthesis, 1 uL of RNase Cocktail (Thermo, AM2286) was added after reverse transcription and incubated at 37°C for 20 minutes. The NEBNext second strand synthesis kit (NEB, E6111) with 100ng random hexamer primers (Promega, C1181) per sample was used for second strand synthesis. Next, dsDNA was purified using AMPure XP beads (Beckman Coulter: A63882) at a ratio of 2 beads to 1 sample. The NEBNext Ultra II End Repair/dA-tailing Module (NEB, E7546S) was used before ligating Nanopore native barcodes (EXP-NBD104&114). Finally, samples were pooled at equal mass and the sequencing adapter (SQK-LSK109) was ligated. 1000 ng DNA was loaded on the Promethion flow cell (FLO-PRO002) and sequenced for 3 days. Two independent replicates were performed.

### Nanopore data analysis

During sequencing, the Nanopore MinKNOW software (version 21.11.7, Guppy version 5.1.13) called bases in real-time by the using the ‘Super-accurate’ base calling setting, demultiplexed samples and trimmed barcodes. For replicate 1, the 0 minute after transfection, dEGFP targeting sample had a very low DNA yield after library preparation and consequently much fewer reads than the other samples. Therefore, it was excluded from all analysis. Reads were mapped to the transcriptome using minimap2 (Li, 2018). Transcriptome alignment was done using the *-ax map-ont* arguments, against a merge of the Ensemble Homo_sapiens.GRCh38.cdna.all (release 104), Ensemble Homo_sapiens.GRCh38.ncrna (release 104), the entire pEF-1a_dEGFP^Y66S^-IRIS-PuroR-WPRE plasmid (Supplementary file 1) and the 18S and 28S ribosomal RNA. Mapping rates were similar across samples, with dEGFP targeting samples showing a small decrease at 500 and 1440 minutes after transfection (Supplementary figure 15A). Only the primary alignment was used for analysis. To generate the density plots in Figure 6A and Supplementary figure 11B, all histone transcripts and WDR74 were excluded. Histone transcripts and WDR74 were so abundant that they obscured all other mRNAs in the density plot. The extremely high abundance of WDR74 was caused by a small nuclear RNA mapping to WDR74-204. All analysis was performed in Rstudio, using data.table (Dowle and Srinivasan, 2022) and ShortRead (Morgan et al., 2009) for data loading, Tidyverse (Wickham et al., 2019) for data handling and ggplot2 (Wickham, 2010) for plotting. Raw sequencing reads are available online at https://www.ncbi.nlm.nih.gov/sra/PRJNA912090. The script used for calculating transcript coverages is available on https://github.com/Geijsenlab/cas13.

### RNA sequencing and analysis

For the RNA-seq experiment, HAP1-dEGFP^Y66S^ cells were transfected using iTOP, as described above. The day before transfection, 600 000 cells per well had been plated in 6-well plates. Treatments were: an empty transfection, a transfection with LbuCas13a but no guide RNA, a transfection with LbuCas13a and a non-targeting guide RNA, and finally a transfection with LbuCas13a and a dEGFP targeting guide RNA (SP2). All treatments were done in triplicates. After 16 hours, cells were lysed in TRIzol reagent. Total RNA was isolated and DNase-treated as described above. mRNA was then extracted using a NEBNext Poly(A) mRNA Magnetic Isolation Module (NEB, E7490S) and RNA-seq libraries were prepared using a NEBNext Ultra Directional RNA Library Prep Kit for Illumina (NEB, E7420L). RNA-seq libraries were sequenced on an Illumina NextSeq500 Instrument with at least 10 million reads per library. Quality control by FastQC (Andrews, 2010) and RSeQC (Wang et al., 2012) did not reveal any major issues. An index was generated using the RefSeq GRCh37 assembly and reads were aligned and quantified using STAR (Dobin et al., 2013) and featureCounts (Liao et al., 2014). Differentially expressed genes were determined using DESeq2 (Love et al., 2014). Genes with an adjusted P-value < 0.05 and log2(Fold Change) > |1| were regarded as differentially expressed genes, and they were subjected to subsequent enrichment analysis. GO term and pathway enrichment analysis was done using ShinyGO (Ge et al., 2020) V0.75, supplying all detected transcripts as background. RNA-seq data is available online at the NCBI Gene expression omnibus (GSE220759). The code of the differential gene expression analysis is available on https://github.com/Geijsenlab/cas13.

### Live cell imaging

The live cell imaging protocol used here was adapted from Wallberg *et al*. (Wallberg et al., 2016). HAP1 and HAP1-dEGFP^Y66S^ cells were imaged in imaging medium (FluoroBrite™ DMEM, 10% FBS, 2.5 mM CaCl_2_) supplemented with 2 µL CellTox™ Green (Promega, G8741) and 5 µL Annexin V-AV647 (Invitrogen, A23204) per milliliter. Cells were seeded in 8-well µ-Slides (ibidi, 80827) coated with a 1:100 dilution of Matrigel (Corning, 356231), plating 55 000 cells per well. The next day, iTOP was performed as described above, using an iTOP mixture volume of 200 µL per well. After removing the iTOP buffer, wells were washed with 300 µL PBS before adding 300 µL imaging medium. This extra PBS wash step prevents protein or salt precipitation during imaging. After iTOP, cells were allowed to recover in the cell culture incubator for up to four hours. Next, cells were moved to the Leica AF7000 and allowed to equilibrate for one hour (37°C, 5% CO_2_). During live imaging, pictures were taken every 15 minutes at four positions per well for 20 hours, using a 20x Dry N.A. 0.50 objective. Finally, the amount of Annexin V and CellTox™ Green positive cells were counted with CellProfiler (McQuin et al., 2018) using the IdentifyPrimaryObjects module.

### Cell selection experiments

For the cell selection experiments, HAP1-EGFP cells were mixed with HAP1 cells at a ratio of roughly 4:1. HAP1-EGFP were selected against by introducing LbuCas13a with an EGPF targeting guide RNA (SP2) using iTOP. The percentage of EGFP positive cells was measured by FACS on the BD FACSCanto II (BD Bioscience). To generate HAP1-EGFP lines with different levels of EGFP expression, cells were sorted into fifteen different populations with increasing EGFP intensities using the BD FACSJazz cell sorter (BD Bioscience). We sorted the cells in complete cell culture medium. After sorting, the three populations with the lowest EGFP intensity were cultured in medium without puromycin. The next three lowest populations were cultured at 2 µg/mL puromycin, the following groups of three at 5 µg/mL, 10 µg/mL and finally 20 µg/mL. For the selection experiment, the different EGFP lines were mixed with wildtype HAP1 cells and subjected to selection as described above. Additionally, in parallel they were also plated in a separate well. These parallel wells were harvested in TRIzol immediately after the mixed population had been subjected to iTOP, and used to determine their EGFP expression levels. Conventional RT-qPCR was performed as described above for 5’-3’ RT-qPCR. qPCR primers can be found in supplementary table 2.

To repair the dEGFP*^Δfluor^* mutation, cells were transfected with recombinant AsCas12a protein, guide RNA (Supplementary table 3) and a LAHR repair template (Supplementary table 3) using iTOP. For each well of a 96-well plate, 50 µL of iTOP mixture that contained 20 µL of transfection supplement B, 10 µL of AsCas12a protein (150 µM), 10 µL of guide RNA (300 µM) and 10 µL of 500 pmol repair template in nuclease-free water was used. Three days after iTOP transfection, the cells that had not been edited were selected against by delivering LbuCas13a and a dEGFP*^Δfluor^*targeting guide RNA (Supplementary table 1) using iTOP. Finally, the percentage of EGFP positive cells was assessed on the BD FACSCanto II three days after selection with LbuCas13a.

### Data availability

The datasets and computer code produced in this study are available in the following databases:

- Nanopore sequencing data: NCBI Sequencing Read Archive PRJNA912090 (https://www.ncbi.nlm.nih.gov/sra/PRJNA912090)
- RNA-seq data: Gene Expression Omnibus (https://www.ncbi.nlm.nih.gov/geo/query/acc.cgi?acc=GSE220759)
- Scripts for nanopore sequencing coverage and RNA-seq analysis: GitHub (https://github.com/Geijsenlab/cas13)

## Supporting information

Supplementary file 1 plasmid sequences

Supplementary note 1 on orthologs

Supplementary table 1 IVT oligos

Supplementary table 2 qPCR primers

Supplementary table 3 Cas12 oligo

Supplementary table 4 LbuCas13a cutsites

## Acknowledgements

The authors thank Stefan van der Elst of the Hubrecht Institute FACS facility, the LUMC FACS facility and the Utrecht sequencing facility (USEQ) for technical help. The novo Nordisk Foundation Center for Stem Cell Medicine (reNEW) is supported by a Novo Nordisk Foundation grant number NFF21CC0073729. This work was also supported by funding from NWO-TTW and the Princes Beatrix Muscle Foundation. Z.Z. was supported, in part, by the China Scholarship Council.

## Author contributions

JFB experimental design, execution and help writing the manuscript, ZZ experimental design, execution and help writing the manuscript, ML technical assistance, DK technical assistance, PS experimental design and help writing the manuscript, NG conceptual and experimental design, project supervision and help writing the manuscript.

