## Supplementary figure legends for "Temporal dynamics of collateral RNA cleavage by LbuCas13a in human cells"

**Supplementary figure 1. SDS-PAGE, RNase A 5’-3’ RT-qPCR and RNA yield after dEGFP targeting. A** SDS-PAGE gels of Cas13 orthologs after HIS-tag purification followed by fast protein liquid chromatography (FPLC) gel filtration. All proteins have the expected size. **B** IDT RNaseAlert assay. Cas13 proteins were incubated with an EGFP targeting crRNA and three different concentrations of EGFP target RNA. The cleavage rate was calculated by taking the slope of the initial linear increase in fluorescence intensity. RfxCas13d did not show any activity in this assay.

**Supplementary figure 2. Comparing the collateral activity in cells of different Cas13 Orthologs. A** Total RNA profiles after GAPDH, dEGFP and 18S rRNA targeting with LwaCas13a. **B** Total RNA profiles after GAPDH, dEGFP and 18S rRNA targeting with BzCas13b. **C** Total RNA profiles after GAPDH, dEGFP and 18S rRNA targeting with RspCas13d. RNA isolated 100 minutes after transfection using iTOP. Fluorescence was adjusted to the total area under the curve, to equalize for differences in loaded amount of RNA. n=3.

**Supplementary figure 3. Comparing the collateral activity of LbuCas13a with RfxCas13d. A** Total RNA profiles after electroporation of HAP1-dEGFP cells with 1 µM LbuCas13a (red) or RfxCas13d (blue), and either 1 µM of a non-targeting, dEGFP, GAPDH or 18S rRNA targeting guide RNA. RNA isolated 100 minutes after electroporation and analyzed on the bioanalyzer. Fluorescence was adjusted to the total area under the curve, to equalize for differences in loaded amount of RNA. n=3.

**Supplementary figure 4. Total RNA degradation increases with RNP concentration. A** Total RNA profiles 100 minutes after transduction with different concentrations of LbuCas13a recombinant protein and guide RNA, as determined by bioanalyzer (n=3). An equimolar ratio of protein to guide was used. Fluorescence intensity was adjusted to the total area under the curve of each sample. **B** The area under the curve of the degradation products (area between the dotted lines in **A**) increases with the protein and guide concentration (n=3). The trendline follows y = log(x).

**Supplementary figure 5. Comparing target and collateral RNA cleavage of different Cas13 orthologs using the 5’:3’ RT-qPCR assay. A** 5’-3’ RT-qPCR assay (n=3) on samples transfected with RNase A. RNA was isolated 50 minutes after transfection. P-values calculated by t-test (* < 0.05). RNase A transfections yield the expected increase in 5’ qPCR primer Cq values. **B** Cleavage of target (dEGFP) and collateral (GAPDH) RNA as assessed by the 5’-3’ RT-qPCR assay (n=3). RNA isolated 50 minutes after transfection. Only LbuCas13a shows significant collateral RNA cleavage. P-values calculated by unpaired Welch t-test (ns = not significant, * < 0.05, ** < 0.01). Small dots are independent replicates. Big dot and line show mean and standard error of the mean. **C** 5’-3’ ddCq values (n=3) from RNA isolated 100 minutes after electroporation with either 1 µM LbuCas13a or RfxCas13d RNP. All targeting treatments were compared with the non-targeting control using a t-test, numbers are p-values.

**Supplementary figure 6. RNP delivery by electroporation also results in collateral RNA cleavage. A** Total RNA profiles at 200 and 400 minutes after electroporation with LbuCas13a and a GAPDH, RPS19, 18S rRNA or non-targeting guide RNA. **B** Total RNA profiles 200 minutes after electroporation of HAP1 and HEK293T cells. For both **A** and **B**: fluorescence intensity was adjusted to the total area under the curve of each sample, and n=3.

**Supplementary figure 7. No collateral cleavage when the LbuCas13a expression vector is transiently transfected. A** Western blot after transient transfection of the LbuCas13a expression vector, using an HA-tag antibody. Western blot was performed on unsorted cells with an average transfection efficiency of 9.2%. Three independent transfections were performed. **B** Total RNA profiles 100 minutes after iTOP transfection, as determined by bioanalyzer (n=3). **C** Western blot after stable lentiviral integration of LbuCas13a or RfxCas13d expression plasmids. An HA-tag antibody was used for both Cas13 orthologs.

**Supplementary figure 8. Generating lines with various levels of EGFP expression and gene editing strategy. A** Gating strategy to sort different EGFP intensities. **B** qPCR on cells gated for different GFP intensities. A wide range of EGFP expression levels were recovered after sorting. **C** Schematic showing the LAHR editing strategy, see Zhao et al. for details (Zhao *et al*, 2022).

**Supplementary figure 9. RNA-seq data analysis. A** Targeting with LbuCas13a increase the abundance reads mapped to introns relative to reads mapped to exons. **B** PCA plot of the RNA-seq samples using the default setting of the plotPCA function in DESeq2 (Love *et al*, 2014). **C** Top enriched GO molecular functions using the significantly upregulated genes. Plot generated using ShinyGo (Ge *et al*, 2020). **D** KEGG pathway enrichment of the significantly upregulated genes. Plot generated using ShinyGo (Ge *et al*, 2020).

**Supplementary figure 10. Nanopore library preparation and read lengths. A** Bioanalyzer total RNA profiles of the samples send for Nanopore sequencing. **B** Bioanalyzer RNA profile after rRNA depletion. Legends same as **A.** **C** Rolling mean (k = 20) of the read lengths, corrected by the total amount of reads in each sample. The spike at ~4000 nt in the 50 minute replicate 1 sample is probably rRNA contamination. Legends same as **A.**

**Supplementary figure 11. Protein coding transcripts are affected by collateral cleavage. A** Percentage of all reads that mapped to various transcript biotypes. rRNA mapped reads were excluded due to variability in rRNA depletion efficiencies. **B** Density plot of read length versus reference transcript length of all reads mapped to protein coding transcripts, excluding histone genes and WDR74 (see M&M). Data from replicate 1.

**Supplementary figure 12. Relative transcript coverage of several mRNAs**. **A** Nanopore sequencing relative transcript coverage of GAPDH-205, **B** ACTB-217 and **C** PKM-202. Columns show minutes after transfection. Relative transcript coverage values range from 0 to 1. A value of 0 means none of the reads mapped to this transcript cover that specific nucleotide position. A value of 1 means all reads mapped to this transcript cover that specific nucleotide. Thus, the more reads covering only part of the transcript, the lower the overall relative coverage along the entire transcript will be. We suspect dips in transcript coverage are LbuCas13a cleavage sites.

**Supplementary figure 13. Relative transcript coverage of several cytoplasmic ncRNA. A** Nanopore relative transcript coverage of RN7SL1-201, **B** 18S rRNA and **C** 28S rRNA. Columns show minutes post transduction. Relative transcript coverage values range from 0 to 1. A value of 0 means none of the reads mapped to this transcript cover that specific nucleotide position. A value of 1 means all reads mapped to this transcript cover that specific nucleotide. Thus, the more reads covering only part of the transcript, the lower the overall relative coverage along the entire transcript will be. We suspect dips in transcript coverage are LbuCas13a cleavage sites.

**Supplementary figure 14. Relative transcript coverage of several nuclear and a mitochondrial RNA. A** Relative transcript coverage of U1, **B** RN7SK and **C** MT-RNR1. Columns show minutes post transduction. Relative transcript coverage values range from 0 to 1. A value of 0 means none of the reads mapped to this transcript cover that specific nucleotide position. A value of 1 means all reads mapped to this transcript cover that specific nucleotide. Thus, the more reads covering only part of the transcript, the lower the overall relative coverage along the entire transcript will be. The transcript coverage of these nuclear and mitochondrial RNAs does not seem to be affected by dEGFP targeting with LbuCas13a.

**Supplementary figure 15. Effects of Cas13 targeting on Nanopore mapping rates. A** The mapping rate of every sample submitted for Nanopore sequencing. Reads were mapped using minimap2, see methods. In replicate 1, the 0 minute after iTOP dEGFP Targeting sample had a low mapping rate. This sample was excluded from all further analysis.

**Supplementary figure 16. RfxCas13d does not show any activity in the RNaseAlert assay. A** RNaseAlert assay of RfxCas13d (n=3). RfxCas13d was incubated with a either a non-targeting (NT) or one of three different dEGFP targeting guide RNAs (SP1, SP2, SP3), and 0, 0.01, 0.1 or 1 nM dEGFP target RNA. Fluorescence was measured every minute. No increase in fluorescence is observed. The dotted horizontal line shows the maximum fluorescence of the RNaseA positive control.

**Supplementary figure 17. RfxCas13d does not exhibit collateral cleavage when the RfxCas13d protein is expressed in the cell. A** Total RNA profiles 100 minutes after electroporations, as determined by bioanalyzer (n=3). **B** Cq values from qPCR on samples from **A** using both a 5’ and a 3’ primer pair on GAPDH and dEGFP (n=3). **C** 5’-3’ ddCq values (n=3) from the qPCR on the samples from **A**. All targeting treatments were compared with the non-targeting control using a t-test (ns = not significant, * < 0.05). **D** Western blot after stable lentiviral integration of a RfxCas13d expression plasmids. An HA-tag antibody was used to detect RfxCas13d.

Ge SX, Jung D, Jung D & Yao R (2020) ShinyGO: A graphical gene-set enrichment tool for animals and plants. *Bioinformatics* 36: 2628–2629

Love MI, Huber W & Anders S (2014) Moderated estimation of fold change and dispersion for RNA-seq data with DESeq2. *Genome Biol* 15

Zhao Z, Shang P, Sage F & Geijsen N (2022) Ligation-assisted homologous recombination enables precise genome editing by deploying both MMEJ and HDR. *Nucleic Acids Res* 50: e62–e62
