## Supplementary file 1 plasmid sequences for "Temporal dynamics of collateral RNA cleavage by LbuCas13a in human cells"

### Plasmids used

#### **pEF-1a_dEGFP-Y66S-IRIS-PuroR-WPRE**

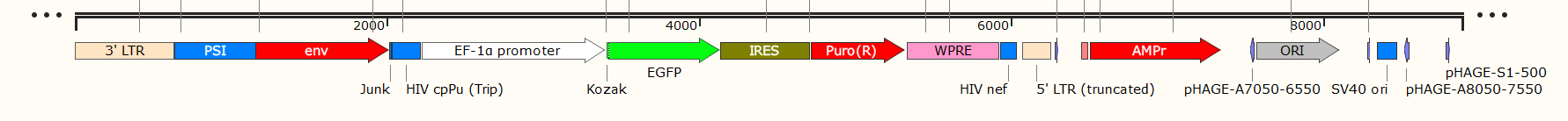

Sequence

tggaagggctaattcactcccaaagaagacaagatatccttgatctgtggatctaccacacacaaggctacttccctgattagcagaactacacaccagggccaggggtcagatatccactgacctttggatggtgctacaagctagtaccagttgagccagataaggtagaagaggccaataaaggagagaacaccagcttgttacaccctgtgagcctgcatgggatggatgacccggagagagaagtgttagagtggaggtttgacagccgcctagcatttcatcacgtggcccgagagctgcatccggagtacttcaagaactgctgatatcgagcttgctacaagggactttccgctggggactttccagggaggcgtggcctgggcgggactggggagtggcgagccctcagatcctgcatataagcagctgctttttgcctgtactgggtctctctggttagaccagatctgagcctgggagctctctggctaactagggaacccactgcttaagcctcaataaagcttgccttgagtgcttcaagtagtgtgtgcccgtctgttgtgtgactctggtaactagagatccctcagacccttttagtcagtgtggaaaatctctagcagtggcgcccgaacagggacttgaaagcgaaagggaaaccagaggagctctctcgacgcaggactcggcttgctgaagcgcgcacggcaagaggcgaggggcggcgactggtgagtacgccaaaaattttgactagcggaggctagaaggagagagatgggtgcgagagcgtcagtattaagcgggggagaattagatcgcgatgggaaaaaattcggttaaggccagggggaaagaaaaaatataaattaaaacatatagtatgggcaagcagggagctagaacgattcgcagttaatcctggcctgttagaaacatcagaaggctgtagacaaatactgggacagctacaaccatcccttcagacaggatcagaagaacttagatcattatataatacagtagcaaccctctattgtgtgcatcaaaggatagagataaaagacaccaaggaagctttagacaagatagaggaagagcaaaacaaaagtaagaccaccgcacagcaagcggccggccgctgatcttcagacctggaggaggagatatgagggacaattggagaagtgaattatataaatataaagtagtaaaaattgaaccattaggagtagcacccaccaaggcaaagagaagagtggtgcagagagaaaaaagagcagtgggaataggagctttgttccttgggttcttgggagcagcaggaagcactatgggcgcagcgtcaatgacgctgacggtacaggccagacaattattgtctggtatagtgcagcagcagaacaatttgctgagggctattgaggcgcaacagcatctgttgcaactcacagtctggggcatcaagcagctccaggcaagaatcctggctgtggaaagatacctaaaggatcaacagctcctggggatttggggttgctctggaaaactcatttgcaccactgctgtgccttggaatgctagttggagtaataaatctctggaacagatttggaatcacacgacctggatggagtgggacagagaaattaacaattacacaagcttaatacactccttaattgaagaatcgcaaaaccagcaagaaaagaatgaacaagaattattggaattagataaatgggcaagtttgtggaattggtttaacataacaaattggctgtggtatataaaattattcataatgatagtaggaggcttggtaggtttaagaatagtttttgctgtactttctatagtgaatagagttaggcagggatattcaccattatcgtttcagacccacctcccaaccccgaggggacccgacaggcccgaaggaatagaagaagaaggtggagagagagacagagacagatccattcgattagtgaacggatctcgacggtatcgccgaattcacaaatggcagtattcatccacaattttaaaagaaaaggggggattggggggtacagtgcaggggaaagaatagtagacataatagcaacagacatacaaactaaagaattacaaaaacaaattacaaaaattcaaaattttcgggtttattacagggacagcagagatccagtttggactagtcgtgaggctccggtgcccgtcagtgggcagagcgcacatcgcccacagtccccgagaagttggggggaggggtcggcaattgaaccggtgcctagagaaggtggcgcggggtaaactgggaaagtgatgtcgtgtactggctccgcctttttcccgagggtgggggagaaccgtatataagtgcagtagtcgccgtgaacgttctttttcgcaacgggtttgccgccagaacacaggtaagtgccgtgtgtggttcccgcgggcctggcctctttacgggttatggcccttgcgtgccttgaattacttccacctggctgcagtacgtgattcttgatcccgagcttcgggttggaagtgggtgggagagttcgaggccttgcgcttaaggagccccttcgcctcgtgcttgagttgaggcctggcctgggcgctggggccgccgcgtgcgaatctggtggcaccttcgcgcctgtctcgctgctttcgataagtctctagccatttaaaatttttgatgacctgctgcgacgctttttttctggcaagatagtcttgtaaatgcgggccaagatctgcacactggtatttcggtttttggggccgcgggcggcgacggggcccgtgcgtcccagcgcacatgttcggcgaggcggggcctgcgagcgcggccaccgagaatcggacgggggtagtctcaagctggccggcctgctctggtgcctggcctcgcgccgccgtgtatcgccccgccctgggcggcaaggctggcccggtcggcaccagttgcgtgagcggaaagatggccgcttcccggccctgctgcagggagctcaaaatggaggacgcggcgctcgggagagcgggcgggtgagtcacccacacaaaggaaaagggcctttccgtcctcagccgtcgcttcatgtgactccacggagtaccgggcgccgtccaggcacctcgattagttctcgagcttttggagtacgtcgtctttaggttggggggaggggttttatgcgatggagtttccccacactgagtgggtggagactgaagttaggccagcttggcacttgatgtaattctccttggaatttgccctttttgagtttggatcttggttcattctcaagcctcagacagtggttcaaagtttttttcttccatttcaggtgtcgtgaagcggccgccaccatggtgagcaagggcgaggagctgttcaccggggtggtgcccatcctggtcgagctggacggcgacgtaaacggccacaagttcagcgtgtccggcgagggcgagggcgatgccacctacggcaagctgacCCtgaagttcatctgcaccaccggcaagctgcccgtgccctggcccacTTtGgtgaccaccctgacctCcggcgtgcagtgcttcagccgctaccccgaccacatgaagcagcacgacttcttcaagtccgccatgcccgaaggctacgtccaggagcgcaccatcttcttcaaggacgacggcaactacaagacccgcgccgaggtgaagttcgagggcgacaccctggtgaaccgcatcgagctgaagggcatcgacttcaaggaggacggcaacatcctggggcacaagctggagtacaactacaacagccacaacgtctatatcatggccgacaagcagaagaacggcatcaaggtgaacttcaagatccgccacaacatcgaggacggcagcgtgcagctcgccgaccactaccagcagaacacccccatcggcgacggccccgtgctgctgcccgacaaccactacctgagcacccagtccgccctgagcaaagaccccaacgagaagcgcgatcacatggtcctgctggagttcgtgaccgccgccgggatcactctcggcatggacgagctgtacaagtaaggatccctcccccccccctaacgttactggccgaagccgcttggaataaggccggtgtgcgtttgtctatatgttattttccaccatattgccgtcttttggcaatgtgagggcccggaaacctggccctgtcttcttgacgagcattcctaggggtctttcccctctcgccaaaggaatgcaaggtctgttgaatgtcgtgaaggaagcagttcctctggaagcttcttgaagacaaacaacgtctgtagcgaccctttgcaggcagcggaaccccccacctggcgacaggtgcctctgcggccaaaagccacgtgtataagatacacctgcaaaggcggcacaaccccagtgccacgttgtgagttggatagttgtggaaagagtcaaatggctctcctcaagcgtattcaacaaggggctgaaggatgcccagaaggtaccccattgtatgggatctgatctggggcctcggtgcacatgctttacatgtgtttagtcgaggttaaaaaaacgtctaggccccccgaaccacggggacgtggttttcctttgaaaaacacgatgataatatggccacacatatgaccgagtacaagcccacggtgcgcctcgccacccgcgacgacgtccccagggccgtacgcaccctcgccgccgcgttcgccgactaccccgccacgcgccacaccgtcgatccggaccgccacatcgagcgggtcaccgagctgcaagaactcttcctcacgcgcgtcgggctcgacatcggcaaggtgtgggtcgcggacgacggcgccgcggtggcggtctggaccacgccggagagcgtcgaagcgggggcggtgttcgccgagatcggcccgcgcatggccgagttgagcggttcccggctggccgcgcagcaacagatggaaggcctcctggcgccgcaccggcccaaggagcccgcgtggttcctggccaccgtcggcgtctcgcccgaccaccagggcaagggtctgggcagcgccgtcgtgctccccggagtggaggcggccgagcgcgccggggtgcccgccttcctggagacctccgcgccccgcaacctccccttctacgagcggctcggcttcaccgtcaccgccgacgtcgaggtgcccgaaggaccgcgcacctggtgcatgacccgcaagcccggtgcctgaatcgatagatcctaatcaacctctggattacaaaatttgtgaaagattgactggtattcttaactatgttgctccttttacgctatgtggatacgctgctttaatgcctttgtatcatgctattgcttcccgtatggctttcattttctcctccttgtataaatcctggttgctgtctctttatgaggagttgtggcccgttgtcaggcaacgtggcgtggtgtgcactgtgtttgctgacgcaacccccactggttggggcattgccaccacctgtcagctcctttccgggactttcgctttccccctccctattgccacggcggaactcatcgccgcctgccttgcccgctgctggacaggggctcggctgttgggcactgacaattccgtggtgttgtcggggaaatcatcgtcctttccttggctgctcgcctgtgttgccacctggattctgcgcgggacgtccttctgctacgtcccttcggccctcaatccagcggaccttccttcccgcggcctgctgccggctctgcggcctcttccgcgtcttcgccttcgccctcagacgagtcggatctccctttgggccgcctccccgcctggtacctttaagaccaatgacttacaaggcagctgtagatcttagccactttttaaaagaaaaggggggactggaagggctaattcactcccaacgaagacaagatcacctgcaggacaggcgcgccctgctttttgcttgtactgggtctctctggttagaccagatctgagcctgggagctctctggctaactagggaacccactgcttaagcctcaataaagcttgccttgagtgcttcaagtagtgtgtgcccgtctgttgtgtgactctggtaactagagatccctcagacccttttagtcagtgtggaaaatctctagcacccgggcgattaaggaaagggctagatcattcttgaagacgaaagggcctcgtgatacgcctatttttataggttaatgtcatgataataatggtttcttagacgtcaggtggcacttttcggggaaatgtgcgcggaacccctatttgtttatttttctaaatacattcaaatatgtatccgctcatgagacaataaccctgataaatgcttcaataatattgaaaaaggaagagtatgagtattcaacatttccgtgtcgcccttattcccttttttgcggcattttgccttcctgtttttgctcacccagaaacgctggtgaaagtaaaagatgctgaagatcagttgggtgcacgagtgggttacatcgaactggatctcaacagcggtaagatccttgagagttttcgccccgaagaacgttttccaatgatgagcacttttaaagttctgctatgtggcgcggtattatcccgtgttgacgccgggcaagagcaactcggtcgccgcatacactattctcagaatgacttggttgagtactcaccagtcacagaaaagcatcttacggatggcatgacagtaagagaattatgcagtgctgccataaccatgagtgataacactgcggccaacttacttctgacaacgatcggaggaccgaaggagctaaccgcttttttgcacaacatgggggatcatgtaactcgccttgatcgttgggaaccggagctgaatgaagccataccaaacgacgagcgtgacaccacgatgcctgtagcaatggcaacaacgttgcgcaaactattaactggcgaactacttactctagcttcccggcaacaattaatagactggatggaggcggataaagttgcaggaccacttctgcgctcggcccttccggctggctggtttattgctgataaatctggagccggtgagcgtgggtctcgcggtatcattgcagcactggggccagatggtaagccctcccgtatcgtagttatctacacgacggggagtcaggcaactatggatgaacgaaatagacagatcgctgagataggtgcctcactgattaagcattggtaactgtcagaccaagtttactcatatatactttagattgatttaaaacttcatttttaatttaaaaggatctaggtgaagatcctttttgataatctcatgaccaaaatcccttaacgtgagttttcgttccactgagcgtcagaccccgtagaaaagatcaaaggatcttcttgagatcctttttttctgcgcgtaatctgctgcttgcaaacaaaaaaaccaccgctaccagcggtggtttgtttgccggatcaagagctaccaactctttttccgaaggtaactggcttcagcagagcgcagataccaaatactgttcttctagtgtagccgtagttaggccaccacttcaagaactctgtagcaccgcctacatacctcgctctgctaatcctgttaccagtggctgctgccagtggcgataagtcgtgtcttaccgggttggactcaagacgatagttaccggataaggcgcagcggtcgggctgaacggggggttcgtgcacacagcccagcttggagcgaacgacctacaccgaactgagatacctacagcgtgagctatgagaaagcgccacgcttcccgaagggagaaaggcggacaggtatccggtaagcggcagggtcggaacaggagagcgcacgagggagcttccagggggaaacgcctggtatctttatagtcctgtcgggtttcgccacctctgacttgagcgtcgatttttgtgatgctcgtcaggggggcggagcctatggaaaaacgccagcaacgcggcctttttacggttcctggccttttgctggccttttgctcacatgttctttcctgcgttatcccctgattctgtggataaccgtattaccgcctttgagtgagctgataccgctcgccgcagccgaacgaccgagcgcagcgagtcagtgagcgaggaagcggaagagcgcccaatacgcaaaccgcctctccccgcgcgttggccgattcattaatgcagcaagctcatggctgactaattttttttatttatgcagaggccgaggccgcctcggcctctgagctattccagaagtagtgaggaggcttttttggaggcctaggcttttgcaaaaagctccccgtggcacgacaggtttcccgactggaaagcgggcagtgagcgcaacgcaattaatgtgagttagctcactcattaggcaccccaggctttacactttatgcttccggctcgtatgttgtgtggaattgtgagcggataacaatttcacacaggaaacagctatgacatgattacgaatttcacaaataaagcatttttttcactgcattctagttgtggtttgtccaaactcatcaatgtatcttatcatgtctggatcaactggataactcaagctaaccaaaatcatcccaaacttcccaccccataccctattaccactgccaattacctgtggtttcatttactctaaacctgtgattcctctgaattattttcattttaaagaaattgtatttgttaaatatgtactacaaacttagtagt

#### **pEF-1a_EGFP-IRIS-PuroR-WPRE**

TggaagggctaattcactcccaaagaagacaagatatccttgatctgtggatctaccacacacaaggctacttccctgattagcagaactacacaccagggccaggggtcagatatccactgacctttggatggtgctacaagctagtaccagttgagccagataaggtagaagaggccaataaaggagagaacaccagcttgttacaccctgtgagcctgcatgggatggatgacccggagagagaagtgttagagtggaggtttgacagccgcctagcatttcatcacgtggcccgagagctgcatccggagtacttcaagaactgctgatatcgagcttgctacaagggactttccgctggggactttccagggaggcgtggcctgggcgggactggggagtggcgagccctcagatcctgcatataagcagctgctttttgcctgtactgggtctctctggttagaccagatctgagcctgggagctctctggctaactagggaacccactgcttaagcctcaataaagcttgccttgagtgcttcaagtagtgtgtgcccgtctgttgtgtgactctggtaactagagatccctcagacccttttagtcagtgtggaaaatctctagcagtggcgcccgaacagggacttgaaagcgaaagggaaaccagaggagctctctcgacgcaggactcggcttgctgaagcgcgcacggcaagaggcgaggggcggcgactggtgagtacgccaaaaattttgactagcggaggctagaaggagagagatgggtgcgagagcgtcagtattaagcgggggagaattagatcgcgatgggaaaaaattcggttaaggccagggggaaagaaaaaatataaattaaaacatatagtatgggcaagcagggagctagaacgattcgcagttaatcctggcctgttagaaacatcagaaggctgtagacaaatactgggacagctacaaccatcccttcagacaggatcagaagaacttagatcattatataatacagtagcaaccctctattgtgtgcatcaaaggatagagataaaagacaccaaggaagctttagacaagatagaggaagagcaaaacaaaagtaagaccaccgcacagcaagcggccggccgctgatcttcagacctggaggaggagatatgagggacaattggagaagtgaattatataaatataaagtagtaaaaattgaaccattaggagtagcacccaccaaggcaaagagaagagtggtgcagagagaaaaaagagcagtgggaataggagctttgttccttgggttcttgggagcagcaggaagcactatgggcgcagcgtcaatgacgctgacggtacaggccagacaattattgtctggtatagtgcagcagcagaacaatttgctgagggctattgaggcgcaacagcatctgttgcaactcacagtctggggcatcaagcagctccaggcaagaatcctggctgtggaaagatacctaaaggatcaacagctcctggggatttggggttgctctggaaaactcatttgcaccactgctgtgccttggaatgctagttggagtaataaatctctggaacagatttggaatcacacgacctggatggagtgggacagagaaattaacaattacacaagcttaatacactccttaattgaagaatcgcaaaaccagcaagaaaagaatgaacaagaattattggaattagataaatgggcaagtttgtggaattggtttaacataacaaattggctgtggtatataaaattattcataatgatagtaggaggcttggtaggtttaagaatagtttttgctgtactttctatagtgaatagagttaggcagggatattcaccattatcgtttcagacccacctcccaaccccgaggggacccgacaggcccgaaggaatagaagaagaaggtggagagagagacagagacagatccattcgattagtgaacggatctcgacggtatcgccgaattcacaaatggcagtattcatccacaattttaaaagaaaaggggggattggggggtacagtgcaggggaaagaatagtagacataatagcaacagacatacaaactaaagaattacaaaaacaaattacaaaaattcaaaattttcgggtttattacagggacagcagagatccagtttggactagtcgtgaggctccggtgcccgtcagtgggcagagcgcacatcgcccacagtccccgagaagttggggggaggggtcggcaattgaaccggtgcctagagaaggtggcgcggggtaaactgggaaagtgatgtcgtgtactggctccgcctttttcccgagggtgggggagaaccgtatataagtgcagtagtcgccgtgaacgttctttttcgcaacgggtttgccgccagaacacaggtaagtgccgtgtgtggttcccgcgggcctggcctctttacgggttatggcccttgcgtgccttgaattacttccacctggctgcagtacgtgattcttgatcccgagcttcgggttggaagtgggtgggagagttcgaggccttgcgcttaaggagccccttcgcctcgtgcttgagttgaggcctggcctgggcgctggggccgccgcgtgcgaatctggtggcaccttcgcgcctgtctcgctgctttcgataagtctctagccatttaaaatttttgatgacctgctgcgacgctttttttctggcaagatagtcttgtaaatgcgggccaagatctgcacactggtatttcggtttttggggccgcgggcggcgacggggcccgtgcgtcccagcgcacatgttcggcgaggcggggcctgcgagcgcggccaccgagaatcggacgggggtagtctcaagctggccggcctgctctggtgcctggcctcgcgccgccgtgtatcgccccgccctgggcggcaaggctggcccggtcggcaccagttgcgtgagcggaaagatggccgcttcccggccctgctgcagggagctcaaaatggaggacgcggcgctcgggagagcgggcgggtgagtcacccacacaaaggaaaagggcctttccgtcctcagccgtcgcttcatgtgactccacggagtaccgggcgccgtccaggcacctcgattagttctcgagcttttggagtacgtcgtctttaggttggggggaggggttttatgcgatggagtttccccacactgagtgggtggagactgaagttaggccagcttggcacttgatgtaattctccttggaatttgccctttttgagtttggatcttggttcattctcaagcctcagacagtggttcaaagtttttttcttccatttcaggtgtcgtgaagcggccgccaccatggtgagcaagggcgaggagctgttcaccggggtggtgcccatcctggtcgagctggacggcgacgtaaacggccacaagttcagcgtgtccggcgagggcgagggcgatgccacctacggcaagctgacCCtgaagttcatctgcaccaccggcaagctgcccgtgccctggcccacTTtGgtgaccaccctgacctAcggcgtgcagtgcttcagccgctaccccgaccacatgaagcagcacgacttcttcaagtccgccatgcccgaaggctacgtccaggagcgcaccatcttcttcaaggacgacggcaactacaagacccgcgccgaggtgaagttcgagggcgacaccctggtgaaccgcatcgagctgaagggcatcgacttcaaggaggacggcaacatcctggggcacaagctggagtacaactacaacagccacaacgtctatatcatggccgacaagcagaagaacggcatcaaggtgaacttcaagatccgccacaacatcgaggacggcagcgtgcagctcgccgaccactaccagcagaacacccccatcggcgacggccccgtgctgctgcccgacaaccactacctgagcacccagtccgccctgagcaaagaccccaacgagaagcgcgatcacatggtcctgctggagttcgtgaccgccgccgggatcactctcggcatggacgagctgtacaagtaaggatccctcccccccccctaacgttactggccgaagccgcttggaataaggccggtgtgcgtttgtctatatgttattttccaccatattgccgtcttttggcaatgtgagggcccggaaacctggccctgtcttcttgacgagcattcctaggggtctttcccctctcgccaaaggaatgcaaggtctgttgaatgtcgtgaaggaagcagttcctctggaagcttcttgaagacaaacaacgtctgtagcgaccctttgcaggcagcggaaccccccacctggcgacaggtgcctctgcggccaaaagccacgtgtataagatacacctgcaaaggcggcacaaccccagtgccacgttgtgagttggatagttgtggaaagagtcaaatggctctcctcaagcgtattcaacaaggggctgaaggatgcccagaaggtaccccattgtatgggatctgatctggggcctcggtgcacatgctttacatgtgtttagtcgaggttaaaaaaacgtctaggccccccgaaccacggggacgtggttttcctttgaaaaacacgatgataatatggccacacatatgaccgagtacaagcccacggtgcgcctcgccacccgcgacgacgtccccagggccgtacgcaccctcgccgccgcgttcgccgactaccccgccacgcgccacaccgtcgatccggaccgccacatcgagcgggtcaccgagctgcaagaactcttcctcacgcgcgtcgggctcgacatcggcaaggtgtgggtcgcggacgacggcgccgcggtggcggtctggaccacgccggagagcgtcgaagcgggggcggtgttcgccgagatcggcccgcgcatggccgagttgagcggttcccggctggccgcgcagcaacagatggaaggcctcctggcgccgcaccggcccaaggagcccgcgtggttcctggccaccgtcggcgtctcgcccgaccaccagggcaagggtctgggcagcgccgtcgtgctccccggagtggaggcggccgagcgcgccggggtgcccgccttcctggagacctccgcgccccgcaacctccccttctacgagcggctcggcttcaccgtcaccgccgacgtcgaggtgcccgaaggaccgcgcacctggtgcatgacccgcaagcccggtgcctgaatcgatagatcctaatcaacctctggattacaaaatttgtgaaagattgactggtattcttaactatgttgctccttttacgctatgtggatacgctgctttaatgcctttgtatcatgctattgcttcccgtatggctttcattttctcctccttgtataaatcctggttgctgtctctttatgaggagttgtggcccgttgtcaggcaacgtggcgtggtgtgcactgtgtttgctgacgcaacccccactggttggggcattgccaccacctgtcagctcctttccgggactttcgctttccccctccctattgccacggcggaactcatcgccgcctgccttgcccgctgctggacaggggctcggctgttgggcactgacaattccgtggtgttgtcggggaaatcatcgtcctttccttggctgctcgcctgtgttgccacctggattctgcgcgggacgtccttctgctacgtcccttcggccctcaatccagcggaccttccttcccgcggcctgctgccggctctgcggcctcttccgcgtcttcgccttcgccctcagacgagtcggatctccctttgggccgcctccccgcctggtacctttaagaccaatgacttacaaggcagctgtagatcttagccactttttaaaagaaaaggggggactggaagggctaattcactcccaacgaagacaagatcacctgcaggacaggcgcgccctgctttttgcttgtactgggtctctctggttagaccagatctgagcctgggagctctctggctaactagggaacccactgcttaagcctcaataaagcttgccttgagtgcttcaagtagtgtgtgcccgtctgttgtgtgactctggtaactagagatccctcagacccttttagtcagtgtggaaaatctctagcacccgggcgattaaggaaagggctagatcattcttgaagacgaaagggcctcgtgatacgcctatttttataggttaatgtcatgataataatggtttcttagacgtcaggtggcacttttcggggaaatgtgcgcggaacccctatttgtttatttttctaaatacattcaaatatgtatccgctcatgagacaataaccctgataaatgcttcaataatattgaaaaaggaagagtatgagtattcaacatttccgtgtcgcccttattcccttttttgcggcattttgccttcctgtttttgctcacccagaaacgctggtgaaagtaaaagatgctgaagatcagttgggtgcacgagtgggttacatcgaactggatctcaacagcggtaagatccttgagagttttcgccccgaagaacgttttccaatgatgagcacttttaaagttctgctatgtggcgcggtattatcccgtgttgacgccgggcaagagcaactcggtcgccgcatacactattctcagaatgacttggttgagtactcaccagtcacagaaaagcatcttacggatggcatgacagtaagagaattatgcagtgctgccataaccatgagtgataacactgcggccaacttacttctgacaacgatcggaggaccgaaggagctaaccgcttttttgcacaacatgggggatcatgtaactcgccttgatcgttgggaaccggagctgaatgaagccataccaaacgacgagcgtgacaccacgatgcctgtagcaatggcaacaacgttgcgcaaactattaactggcgaactacttactctagcttcccggcaacaattaatagactggatggaggcggataaagttgcaggaccacttctgcgctcggcccttccggctggctggtttattgctgataaatctggagccggtgagcgtgggtctcgcggtatcattgcagcactggggccagatggtaagccctcccgtatcgtagttatctacacgacggggagtcaggcaactatggatgaacgaaatagacagatcgctgagataggtgcctcactgattaagcattggtaactgtcagaccaagtttactcatatatactttagattgatttaaaacttcatttttaatttaaaaggatctaggtgaagatcctttttgataatctcatgaccaaaatcccttaacgtgagttttcgttccactgagcgtcagaccccgtagaaaagatcaaaggatcttcttgagatcctttttttctgcgcgtaatctgctgcttgcaaacaaaaaaaccaccgctaccagcggtggtttgtttgccggatcaagagctaccaactctttttccgaaggtaactggcttcagcagagcgcagataccaaatactgttcttctagtgtagccgtagttaggccaccacttcaagaactctgtagcaccgcctacatacctcgctctgctaatcctgttaccagtggctgctgccagtggcgataagtcgtgtcttaccgggttggactcaagacgatagttaccggataaggcgcagcggtcgggctgaacggggggttcgtgcacacagcccagcttggagcgaacgacctacaccgaactgagatacctacagcgtgagctatgagaaagcgccacgcttcccgaagggagaaaggcggacaggtatccggtaagcggcagggtcggaacaggagagcgcacgagggagcttccagggggaaacgcctggtatctttatagtcctgtcgggtttcgccacctctgacttgagcgtcgatttttgtgatgctcgtcaggggggcggagcctatggaaaaacgccagcaacgcggcctttttacggttcctggccttttgctggccttttgctcacatgttctttcctgcgttatcccctgattctgtggataaccgtattaccgcctttgagtgagctgataccgctcgccgcagccgaacgaccgagcgcagcgagtcagtgagcgaggaagcggaagagcgcccaatacgcaaaccgcctctccccgcgcgttggccgattcattaatgcagcaagctcatggctgactaattttttttatttatgcagaggccgaggccgcctcggcctctgagctattccagaagtagtgaggaggcttttttggaggcctaggcttttgcaaaaagctccccgtggcacgacaggtttcccgactggaaagcgggcagtgagcgcaacgcaattaatgtgagttagctcactcattaggcaccccaggctttacactttatgcttccggctcgtatgttgtgtggaattgtgagcggataacaatttcacacaggaaacagctatgacatgattacgaatttcacaaataaagcatttttttcactgcattctagttgtggtttgtccaaactcatcaatgtatcttatcatgtctggatcaactggataactcaagctaaccaaaatcatcccaaacttcccaccccataccctattaccactgccaattacctgtggtttcatttactctaaacctgtgattcctctgaattattttcattttaaagaaattgtatttgttaaatatgtactacaaacttagtagt

#### **pEF-1a_dEGFP**^△^**^fluor^-IRIS-PuroR-WPRE**

tggaagggctaattcactcccaaagaagacaagatatccttgatctgtggatctaccacacacaaggctacttccctgattagcagaactacacaccagggccaggggtcagatatccactgacctttggatggtgctacaagctagtaccagttgagccagataaggtagaagaggccaataaaggagagaacaccagcttgttacaccctgtgagcctgcatgggatggatgacccggagagagaagtgttagagtggaggtttgacagccgcctagcatttcatcacgtggcccgagagctgcatccggagtacttcaagaactgctgatatcgagcttgctacaagggactttccgctggggactttccagggaggcgtggcctgggcgggactggggagtggcgagccctcagatcctgcatataagcagctgctttttgcctgtactgggtctctctggttagaccagatctgagcctgggagctctctggctaactagggaacccactgcttaagcctcaataaagcttgccttgagtgcttcaagtagtgtgtgcccgtctgttgtgtgactctggtaactagagatccctcagacccttttagtcagtgtggaaaatctctagcagtggcgcccgaacagggacttgaaagcgaaagggaaaccagaggagctctctcgacgcaggactcggcttgctgaagcgcgcacggcaagaggcgaggggcggcgactggtgagtacgccaaaaattttgactagcggaggctagaaggagagagatgggtgcgagagcgtcagtattaagcgggggagaattagatcgcgatgggaaaaaattcggttaaggccagggggaaagaaaaaatataaattaaaacatatagtatgggcaagcagggagctagaacgattcgcagttaatcctggcctgttagaaacatcagaaggctgtagacaaatactgggacagctacaaccatcccttcagacaggatcagaagaacttagatcattatataatacagtagcaaccctctattgtgtgcatcaaaggatagagataaaagacaccaaggaagctttagacaagatagaggaagagcaaaacaaaagtaagaccaccgcacagcaagcggccggccgctgatcttcagacctggaggaggagatatgagggacaattggagaagtgaattatataaatataaagtagtaaaaattgaaccattaggagtagcacccaccaaggcaaagagaagagtggtgcagagagaaaaaagagcagtgggaataggagctttgttccttgggttcttgggagcagcaggaagcactatgggcgcagcgtcaatgacgctgacggtacaggccagacaattattgtctggtatagtgcagcagcagaacaatttgctgagggctattgaggcgcaacagcatctgttgcaactcacagtctggggcatcaagcagctccaggcaagaatcctggctgtggaaagatacctaaaggatcaacagctcctggggatttggggttgctctggaaaactcatttgcaccactgctgtgccttggaatgctagttggagtaataaatctctggaacagatttggaatcacacgacctggatggagtgggacagagaaattaacaattacacaagcttaatacactccttaattgaagaatcgcaaaaccagcaagaaaagaatgaacaagaattattggaattagataaatgggcaagtttgtggaattggtttaacataacaaattggctgtggtatataaaattattcataatgatagtaggaggcttggtaggtttaagaatagtttttgctgtactttctatagtgaatagagttaggcagggatattcaccattatcgtttcagacccacctcccaaccccgaggggacccgacaggcccgaaggaatagaagaagaaggtggagagagagacagagacagatccattcgattagtgaacggatctcgacggtatcgccgaattcacaaatggcagtattcatccacaattttaaaagaaaaggggggattggggggtacagtgcaggggaaagaatagtagacataatagcaacagacatacaaactaaagaattacaaaaacaaattacaaaaattcaaaattttcgggtttattacagggacagcagagatccagtttggactagtcgtgaggctccggtgcccgtcagtgggcagagcgcacatcgcccacagtccccgagaagttggggggaggggtcggcaattgaaccggtgcctagagaaggtggcgcggggtaaactgggaaagtgatgtcgtgtactggctccgcctttttcccgagggtgggggagaaccgtatataagtgcagtagtcgccgtgaacgttctttttcgcaacgggtttgccgccagaacacaggtaagtgccgtgtgtggttcccgcgggcctggcctctttacgggttatggcccttgcgtgccttgaattacttccacctggctgcagtacgtgattcttgatcccgagcttcgggttggaagtgggtgggagagttcgaggccttgcgcttaaggagccccttcgcctcgtgcttgagttgaggcctggcctgggcgctggggccgccgcgtgcgaatctggtggcaccttcgcgcctgtctcgctgctttcgataagtctctagccatttaaaatttttgatgacctgctgcgacgctttttttctggcaagatagtcttgtaaatgcgggccaagatctgcacactggtatttcggtttttggggccgcgggcggcgacggggcccgtgcgtcccagcgcacatgttcggcgaggcggggcctgcgagcgcggccaccgagaatcggacgggggtagtctcaagctggccggcctgctctggtgcctggcctcgcgccgccgtgtatcgccccgccctgggcggcaaggctggcccggtcggcaccagttgcgtgagcggaaagatggccgcttcccggccctgctgcagggagctcaaaatggaggacgcggcgctcgggagagcgggcgggtgagtcacccacacaaaggaaaagggcctttccgtcctcagccgtcgcttcatgtgactccacggagtaccgggcgccgtccaggcacctcgattagttctcgagcttttggagtacgtcgtctttaggttggggggaggggttttatgcgatggagtttccccacactgagtgggtggagactgaagttaggccagcttggcacttgatgtaattctccttggaatttgccctttttgagtttggatcttggttcattctcaagcctcagacagtggttcaaagtttttttcttccatttcaggtgtcgtgaagcggccgccaccatggtgagcaagggcgaggagctgttcaccggggtggtgcccatcctggtcgagctggacggcgacgtaaacggccacaagttcagcgtgtccggcgagggcgagggcgatgccacctacggcaagctgaccctgaaAttcatctgcaccaccggcaagctgcccgtgccctggcccaccctcgtgaccaccctggtgcagtgTttcagccgctaccccgaccacatgaagcagcacgacttcttcaagtccgccatgcccgaaggctacgtccaggagcgcaccatcttcttcaaggacgacggcaactacaagacccgcgccgaggtgaagttcgagggcgacaccctggtgaaccgcatcgagctgaagggcatcgacttcaaggaggacggcaacatcctggggcacaagctggagtacaactacaacagccacaacgtctatatcatggccgacaagcagaagaacggcatcaaggtgaacttcaagatccgccacaacatcgaggacggcagcgtgcagctcgccgaccactaccagcagaacacccccatcggcgacggccccgtgctgctgcccgacaaccactacctgagcacccagtccgccctgagcaaagaccccaacgagaagcgcgatcacatggtcctgctggagttcgtgaccgccgccgggatcactctcggcatggacgagctgtacaagtaaggatccctcccccccccctaacgttactggccgaagccgcttggaataaggccggtgtgcgtttgtctatatgttattttccaccatattgccgtcttttggcaatgtgagggcccggaaacctggccctgtcttcttgacgagcattcctaggggtctttcccctctcgccaaaggaatgcaaggtctgttgaatgtcgtgaaggaagcagttcctctggaagcttcttgaagacaaacaacgtctgtagcgaccctttgcaggcagcggaaccccccacctggcgacaggtgcctctgcggccaaaagccacgtgtataagatacacctgcaaaggcggcacaaccccagtgccacgttgtgagttggatagttgtggaaagagtcaaatggctctcctcaagcgtattcaacaaggggctgaaggatgcccagaaggtaccccattgtatgggatctgatctggggcctcggtgcacatgctttacatgtgtttagtcgaggttaaaaaaacgtctaggccccccgaaccacggggacgtggttttcctttgaaaaacacgatgataatatggccacacatatgaccgagtacaagcccacggtgcgcctcgccacccgcgacgacgtccccagggccgtacgcaccctcgccgccgcgttcgccgactaccccgccacgcgccacaccgtcgatccggaccgccacatcgagcgggtcaccgagctgcaagaactcttcctcacgcgcgtcgggctcgacatcggcaaggtgtgggtcgcggacgacggcgccgcggtggcggtctggaccacgccggagagcgtcgaagcgggggcggtgttcgccgagatcggcccgcgcatggccgagttgagcggttcccggctggccgcgcagcaacagatggaaggcctcctggcgccgcaccggcccaaggagcccgcgtggttcctggccaccgtcggcgtctcgcccgaccaccagggcaagggtctgggcagcgccgtcgtgctccccggagtggaggcggccgagcgcgccggggtgcccgccttcctggagacctccgcgccccgcaacctccccttctacgagcggctcggcttcaccgtcaccgccgacgtcgaggtgcccgaaggaccgcgcacctggtgcatgacccgcaagcccggtgcctgaatcgatagatcctaatcaacctctggattacaaaatttgtgaaagattgactggtattcttaactatgttgctccttttacgctatgtggatacgctgctttaatgcctttgtatcatgctattgcttcccgtatggctttcattttctcctccttgtataaatcctggttgctgtctctttatgaggagttgtggcccgttgtcaggcaacgtggcgtggtgtgcactgtgtttgctgacgcaacccccactggttggggcattgccaccacctgtcagctcctttccgggactttcgctttccccctccctattgccacggcggaactcatcgccgcctgccttgcccgctgctggacaggggctcggctgttgggcactgacaattccgtggtgttgtcggggaaatcatcgtcctttccttggctgctcgcctgtgttgccacctggattctgcgcgggacgtccttctgctacgtcccttcggccctcaatccagcggaccttccttcccgcggcctgctgccggctctgcggcctcttccgcgtcttcgccttcgccctcagacgagtcggatctccctttgggccgcctccccgcctggtacctttaagaccaatgacttacaaggcagctgtagatcttagccactttttaaaagaaaaggggggactggaagggctaattcactcccaacgaagacaagatcacctgcaggacaggcgcgccctgctttttgcttgtactgggtctctctggttagaccagatctgagcctgggagctctctggctaactagggaacccactgcttaagcctcaataaagcttgccttgagtgcttcaagtagtgtgtgcccgtctgttgtgtgactctggtaactagagatccctcagacccttttagtcagtgtggaaaatctctagcacccgggcgattaaggaaagggctagatcattcttgaagacgaaagggcctcgtgatacgcctatttttataggttaatgtcatgataataatggtttcttagacgtcaggtggcacttttcggggaaatgtgcgcggaacccctatttgtttatttttctaaatacattcaaatatgtatccgctcatgagacaataaccctgataaatgcttcaataatattgaaaaaggaagagtatgagtattcaacatttccgtgtcgcccttattcccttttttgcggcattttgccttcctgtttttgctcacccagaaacgctggtgaaagtaaaagatgctgaagatcagttgggtgcacgagtgggttacatcgaactggatctcaacagcggtaagatccttgagagttttcgccccgaagaacgttttccaatgatgagcacttttaaagttctgctatgtggcgcggtattatcccgtgttgacgccgggcaagagcaactcggtcgccgcatacactattctcagaatgacttggttgagtactcaccagtcacagaaaagcatcttacggatggcatgacagtaagagaattatgcagtgctgccataaccatgagtgataacactgcggccaacttacttctgacaacgatcggaggaccgaaggagctaaccgcttttttgcacaacatgggggatcatgtaactcgccttgatcgttgggaaccggagctgaatgaagccataccaaacgacgagcgtgacaccacgatgcctgtagcaatggcaacaacgttgcgcaaactattaactggcgaactacttactctagcttcccggcaacaattaatagactggatggaggcggataaagttgcaggaccacttctgcgctcggcccttccggctggctggtttattgctgataaatctggagccggtgagcgtgggtctcgcggtatcattgcagcactggggccagatggtaagccctcccgtatcgtagttatctacacgacggggagtcaggcaactatggatgaacgaaatagacagatcgctgagataggtgcctcactgattaagcattggtaactgtcagaccaagtttactcatatatactttagattgatttaaaacttcatttttaatttaaaaggatctaggtgaagatcctttttgataatctcatgaccaaaatcccttaacgtgagttttcgttccactgagcgtcagaccccgtagaaaagatcaaaggatcttcttgagatcctttttttctgcgcgtaatctgctgcttgcaaacaaaaaaaccaccgctaccagcggtggtttgtttgccggatcaagagctaccaactctttttccgaaggtaactggcttcagcagagcgcagataccaaatactgttcttctagtgtagccgtagttaggccaccacttcaagaactctgtagcaccgcctacatacctcgctctgctaatcctgttaccagtggctgctgccagtggcgataagtcgtgtcttaccgggttggactcaagacgatagttaccggataaggcgcagcggtcgggctgaacggggggttcgtgcacacagcccagcttggagcgaacgacctacaccgaactgagatacctacagcgtgagctatgagaaagcgccacgcttcccgaagggagaaaggcggacaggtatccggtaagcggcagggtcggaacaggagagcgcacgagggagcttccagggggaaacgcctggtatctttatagtcctgtcgggtttcgccacctctgacttgagcgtcgatttttgtgatgctcgtcaggggggcggagcctatggaaaaacgccagcaacgcggcctttttacggttcctggccttttgctggccttttgctcacatgttctttcctgcgttatcccctgattctgtggataaccgtattaccgcctttgagtgagctgataccgctcgccgcagccgaacgaccgagcgcagcgagtcagtgagcgaggaagcggaagagcgcccaatacgcaaaccgcctctccccgcgcgttggccgattcattaatgcagcaagctcatggctgactaattttttttatttatgcagaggccgaggccgcctcggcctctgagctattccagaagtagtgaggaggcttttttggaggcctaggcttttgcaaaaagctccccgtggcacgacaggtttcccgactggaaagcgggcagtgagcgcaacgcaattaatgtgagttagctcactcattaggcaccccaggctttacactttatgcttccggctcgtatgttgtgtggaattgtgagcggataacaatttcacacaggaaacagctatgacatgattacgaatttcacaaataaagcatttttttcactgcattctagttgtggtttgtccaaactcatcaatgtatcttatcatgtctggatcaactggataactcaagctaaccaaaatcatcccaaacttcccaccccataccctattaccactgccaattacctgtggtttcatttactctaaacctgtgattcctctgaattattttcattttaaagaaattgtatttgttaaatatgtactacaaacttagtagt

#### pET15_LbuCas13a-6xHIS

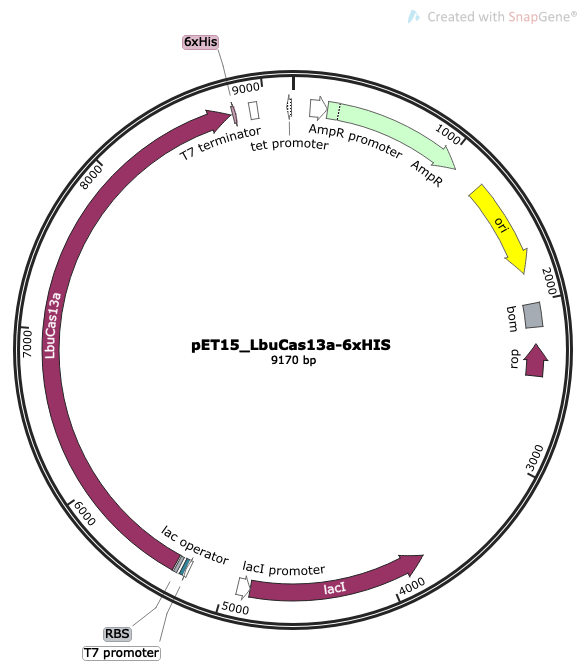

sequence:

TTCTTGAAGACGAAAGGGCCTCGTGATACGCCTATTTTTATAGGTTAATGTCATGATAATAATGGTTTCTTAGACGTCAGGTGGCACTTTTCGGGGAAATGTGCGCGGAACCCCTATTTGTTTATTTTTCTAAATACATTCAAATATGTATCCGCTCATGAGACAATAACCCTGATAAATGCTTCAATAATATTGAAAAAGGAAGAGTATGAGTATTCAACATTTCCGTGTCGCCCTTATTCCCTTTTTTGCGGCATTTTGCCTTCCTGTTTTTGCTCACCCAGAAACGCTGGTGAAAGTAAAAGATGCTGAAGATCAGTTGGGTGCACGAGTGGGTTACATCGAACTGGATCTCAACAGCGGTAAGATCCTTGAGAGTTTTCGCCCCGAAGAACGTTTTCCAATGATGAGCACTTTTAAAGTTCTGCTATGTGGCGCGGTATTATCCCGTGTTGACGCCGGGCAAGAGCAACTCGGTCGCCGCATACACTATTCTCAGAATGACTTGGTTGAGTACTCACCAGTCACAGAAAAGCATCTTACGGATGGCATGACAGTAAGAGAATTATGCAGTGCTGCCATAACCATGAGTGATAACACTGCGGCCAACTTACTTCTGACAACGATCGGAGGACCGAAGGAGCTAACCGCTTTTTTGCACAACATGGGGGATCATGTAACTCGCCTTGATCGTTGGGAACCGGAGCTGAATGAAGCCATACCAAACGACGAGCGTGACACCACGATGCCTGCAGCAATGGCAACAACGTTGCGCAAACTATTAACTGGCGAACTACTTACTCTAGCTTCCCGGCAACAATTAATAGACTGGATGGAGGCGGATAAAGTTGCAGGACCACTTCTGCGCTCGGCCCTTCCGGCTGGCTGGTTTATTGCTGATAAATCTGGAGCCGGTGAGCGTGGGTCTCGCGGTATCATTGCAGCACTGGGGCCAGATGGTAAGCCCTCCCGTATCGTAGTTATCTACACGACGGGGAGTCAGGCAACTATGGATGAACGAAATAGACAGATCGCTGAGATAGGTGCCTCACTGATTAAGCATTGGTAACTGTCAGACCAAGTTTACTCATATATACTTTAGATTGATTTAAAACTTCATTTTTAATTTAAAAGGATCTAGGTGAAGATCCTTTTTGATAATCTCATGACCAAAATCCCTTAACGTGAGTTTTCGTTCCACTGAGCGTCAGACCCCGTAGAAAAGATCAAAGGATCTTCTTGAGATCCTTTTTTTCTGCGCGTAATCTGCTGCTTGCAAACAAAAAAACCACCGCTACCAGCGGTGGTTTGTTTGCCGGATCAAGAGCTACCAACTCTTTTTCCGAAGGTAACTGGCTTCAGCAGAGCGCAGATACCAAATACTGTCCTTCTAGTGTAGCCGTAGTTAGGCCACCACTTCAAGAACTCTGTAGCACCGCCTACATACCTCGCTCTGCTAATCCTGTTACCAGTGGCTGCTGCCAGTGGCGATAAGTCGTGTCTTACCGGGTTGGACTCAAGACGATAGTTACCGGATAAGGCGCAGCGGTCGGGCTGAACGGGGGGTTCGTGCACACAGCCCAGCTTGGAGCGAACGACCTACACCGAACTGAGATACCTACAGCGTGAGCTATGAGAAAGCGCCACGCTTCCCGAAGGGAGAAAGGCGGACAGGTATCCGGTAAGCGGCAGGGTCGGAACAGGAGAGCGCACGAGGGAGCTTCCAGGGGGAAACGCCTGGTATCTTTATAGTCCTGTCGGGTTTCGCCACCTCTGACTTGAGCGTCGATTTTTGTGATGCTCGTCAGGGGGGCGGAGCCTATGGAAAAACGCCAGCAACGCGGCCTTTTTACGGTTCCTGGCCTTTTGCTGGCCTTTTGCTCACATGTTCTTTCCTGCGTTATCCCCTGATTCTGTGGATAACCGTATTACCGCCTTTGAGTGAGCTGATACCGCTCGCCGCAGCCGAACGACCGAGCGCAGCGAGTCAGTGAGCGAGGAAGCGGAAGAGCGCCTGATGCGGTATTTTCTCCTTACGCATCTGTGCGGTATTTCACACCGCATATATGGTGCACTCTCAGTACAATCTGCTCTGATGCCGCATAGTTAAGCCAGTATACACTCCGCTATCGCTACGTGACTGGGTCATGGCTGCGCCCCGACACCCGCCAACACCCGCTGACGCGCCCTGACGGGCTTGTCTGCTCCCGGCATCCGCTTACAGACAAGCTGTGACCGTCTCCGGGAGCTGCATGTGTCAGAGGTTTTCACCGTCATCACCGAAACGCGCGAGGCAGCTGCGGTAAAGCTCATCAGCGTGGTCGTGAAGCGATTCACAGATGTCTGCCTGTTCATCCGCGTCCAGCTCGTTGAGTTTCTCCAGAAGCGTTAATGTCTGGCTTCTGATAAAGCGGGCCATGTTAAGGGCGGTTTTTTCCTGTTTGGTCACTGATGCCTCCGTGTAAGGGGGATTTCTGTTCATGGGGGTAATGATACCGATGAAACGAGAGAGGATGCTCACGATACGGGTTACTGATGATGAACATGCCCGGTTACTGGAACGTTGTGAGGGTAAACAACTGGCGGTATGGATGCGGCGGGACCAGAGAAAAATCACTCAGGGTCAATGCCAGCGCTTCGTTAATACAGATGTAGGTGTTCCACAGGGTAGCCAGCAGCATCCTGCGATGCAGATCCGGAACATAATGGTGCAGGGCGCTGACTTCCGCGTTTCCAGACTTTACGAAACACGGAAACCGAAGACCATTCATGTTGTTGCTCAGGTCGCAGACGTTTTGCAGCAGCAGTCGCTTCACGTTCGCTCGCGTATCGGTGATTCATTCTGCTAACCAGTAAGGCAACCCCGCCAGCCTAGCCGGGTCCTCAACGACAGGAGCACGATCATGCGCACCCGTGGCCAGGACCCAACGCTGCCCGAGATGCGCCGCGTGCGGCTGCTGGAGATGGCGGACGCGATGGATATGTTCTGCCAAGGGTTGGTTTGCGCATTCACAGTTCTCCGCAAGAATTGATTGGCTCCAATTCTTGGAGTGGTGAATCCGTTAGCGAGGTGCCGCCGGCTTCCATTCAGGTCGAGGTGGCCCGGCTCCATGCACCGCGACGCAACGCGGGGAGGCAGACAAGGTATAGGGCGGCGCCTACAATCCATGCCAACCCGTTCCATGTGCTCGCCGAGGCGGCATAAATCGCCGTGACGATCAGCGGTCCAGTGATCGAAGTTAGGCTGGTAAGAGCCGCGAGCGATCCTTGAAGCTGTCCCTGATGGTCGTCATCTACCTGCCTGGACAGCATGGCCTGCAACGCGGGCATCCCGATGCCGCCGGAAGCGAGAAGAATCATAATGGGGAAGGCCATCCAGCCTCGCGTCGCGAACGCCAGCAAGACGTAGCCCAGCGCGTCGGCCGCCATGCCGGCGATAATGGCCTGCTTCTCGCCGAAACGTTTGGTGGCGGGACCAGTGACGAAGGCTTGAGCGAGGGCGTGCAAGATTCCGAATACCGCAAGCGACAGGCCGATCATCGTCGCGCTCCAGCGAAAGCGGTCCTCGCCGAAAATGACCCAGAGCGCTGCCGGCACCTGTCCTACGAGTTGCATGATAAAGAAGACAGTCATAAGTGCGGCGACGATAGTCATGCCCCGCGCCCACCGGAAGGAGCTGACTGGGTTGAAGGCTCTCAAGGGCATCGGTCGAGATCCCGGTGCCTAATGAGTGAGCTAACTTACATTAATTGCGTTGCGCTCACTGCCCGCTTTCCAGTCGGGAAACCTGTCGTGCCAGCTGCATTAATGAATCGGCCAACGCGCGGGGAGAGGCGGTTTGCGTATTGGGCGCCAGGGTGGTTTTTCTTTTCACCAGTGAGACGGGCAACAGCTGATTGCCCTTCACCGCCTGGCCCTGAGAGAGTTGCAGCAAGCGGTCCACGCTGGTTTGCCCCAGCAGGCGAAAATCCTGTTTGATGGTGGTTAACGGCGGGATATAACATGAGCTGTCTTCGGTATCGTCGTATCCCACTACCGAGATATCCGCACCAACGCGCAGCCCGGACTCGGTAATGGCGCGCATTGCGCCCAGCGCCATCTGATCGTTGGCAACCAGCATCGCAGTGGGAACGATGCCCTCATTCAGCATTTGCATGGTTTGTTGAAAACCGGACATGGCACTCCAGTCGCCTTCCCGTTCCGCTATCGGCTGAATTTGATTGCGAGTGAGATATTTATGCCAGCCAGCCAGACGCAGACGCGCCGAGACAGAACTTAATGGGCCCGCTAACAGCGCGATTTGCTGGTGACCCAATGCGACCAGATGCTCCACGCCCAGTCGCGTACCGTCTTCATGGGAGAAAATAATACTGTTGATGGGTGTCTGGTCAGAGACATCAAGAAATAACGCCGGAACATTAGTGCAGGCAGCTTCCACAGCAATGGCATCCTGGTCATCCAGCGGATAGTTAATGATCAGCCCACTGACGCGTTGCGCGAGAAGATTGTGCACCGCCGCTTTACAGGCTTCGACGCCGCTTCGTTCTACCATCGACACCACCACGCTGGCACCCAGTTGATCGGCGCGAGATTTAATCGCCGCGACAATTTGCGACGGCGCGTGCAGGGCCAGACTGGAGGTGGCAACGCCAATCAGCAACGACTGTTTGCCCGCCAGTTGTTGTGCCACGCGGTTGGGAATGTAATTCAGCTCCGCCATCGCCGCTTCCACTTTTTCCCGCGTTTTCGCAGAAACGTGGCTGGCCTGGTTCACCACGCGGGAAACGGTCTGATAAGAGACACCGGCATACTCTGCGACATCGTATAACGTTACTGGTTTCACATTCACCACCCTGAATTGACTCTCTTCCGGGCGCTATCATGCCATACCGCGAAAGGTTTTGCGCCATTCGATGGTGTCCGGGATCTCGACGCTCTCCCTTATGCGACTCCTGCATTAGGAAGCAGCCCAGTAGTAGGTTGAGGCCGTTGAGCACCGCCGCCGCAAGGAATGGTGCATGCAAGGAGATGGCGCCCAACAGTCCCCCGGCCACGGGGCCTGCCACCATACCCACGCCGAAACAAGCGCTCATGAGCCCGAAGTGGCGAGCCCGATCTTCCCCATCGGTGATGTCGGCGATATAGGCGCCAGCAACCGCACCTGTGGCGCCGGTGATGCCGGCCACGATGCGTCCGGCGTAGAGGATCGAGATCTCGATCCCGCGAAATTAATACGACTCACTATAGGGGAATTGTGAGCGGATAACAATTCCCCTCTAGAAATAATTTTGTTTAACTTTAAGAAGGAGATATACCatgaaagtgacgaaggtaggaggcatttcgcataagaagtacacgtccgaaggccgcttagtgaagtcagaatcggaagaaaatcgcacagacgaacgtctgtcggcgttgcttaatatgcgccttgacatgtatatcaagaatcccagcagcacggaaaccaaggaaaatcaaaaacgcattgggaaattaaagaaattcttctcaaacaaaatggtctatcttaaagacaataccttgagtttgaagaatgggaaaaaggagaacattgatcgtgagtattctgagactgacatccttgagagcgatgtccgtgacaagaaaaacttcgccgtgttgaaaaagatctatctgaatgaaaacgtgaactcggaggaattggaagtttttcgtaacgacattaagaagaaactgaacaaaatcaacagcctgaagtactcatttgaaaagaataaggcgaattatcaaaagattaatgagaataacatcgagaaggttgaaggtaagtcaaagcgtaacattatttacgattattatcgtgagtcagcgaaacgtgacgcttatgtaagcaatgtgaaagaagcctttgataagctttacaaggaagaggacattgcaaaacttgttcttgaaattgagaaccttacgaagttagagaaatacaagattcgcgagttctaccacgaaattattggacgtaagaatgacaaggaaaactttgcaaaaatcatctacgaagaaatccagaatgttaataacatgaaagagttgatcgagaaggtaccggacatgagtgagttgaaaaagagccaagtattttacaagtattacttagacaaagaagagttgaacgacaagaacatcaaatacgcgttttgtcatttcgtggaaatcgaaatgagtcagttgctgaagaactacgtatataagcgcttaagtaatatctcgaatgacaaaattaagcgtatctttgaataccagaacttgaaaaaattgatcgaaaataagctgttaaacaaacttgacacgtacgtccgtaattgtggaaagtataattattatttgcaagacggcgaaattgccacttcagatttcatcgcccgcaaccgtcagaatgaagcgtttcttcgcaacatcattggggtgtcatctgtggcctacttttctcttcgcaacattcttgaaacggagaacgagaatgatattactgggcgtatgcgcggcaaaacagttaagaacaataaaggtgaagagaagtacgtgtccggagaagttgataagatctataatgaaaataagaagaacgaggttaaggagaacttaaaaatgttctattcgtacgatttcaatatggacaacaagaatgaaatcgaagatttcttcgccaacatcgacgaggcgatttcttccatccgtcacggtattgtccacttcaacttggaattagaaggtaaggatatctttgcgttcaagaacattgcgccatccgaaatctcaaagaagatgtttcagaatgagattaacgagaaaaaactgaaattgaagatctttcgtcaactgaactctgccaacgtgttccgctatctcgaaaagtataaaattctgaattaccttaaacgtacacgcttcgagtttgtcaataaaaatatcccattcgtcccgtctttcaccaaattatattcgcgcattgatgacctgaagaatagtcttgggatttactggaaaactccgaaaacaaacgacgacaataagactaaggagattattgatgcccaaatctatttgcttaaaaacatctattacggggagttcctgaattatttcatgtcgaacaatggtaatttctttgagatttctaaagaaatcatcgaattgaacaagaacgataaacgcaacttaaagactgggttttacaagctgcaaaagtttgaagacatccaggagaagattccaaaggaatacttggcgaatatccagtccctgtacatgattaatgccggtaatcaggacgaagaagaaaaggacacttatattgatttcattcaaaagatcttcttaaagggatttatgacgtatcttgctaataacggtcgtttaagtctgatttacatcggctcggatgaagaaacaaatacgtcattagcagaaaagaagcaagagtttgacaagttcttgaagaagtacgagcagaacaataatatcaagatcccctatgagatcaatgaattcctgcgtgagatcaaactgggaaacatcctgaagtatactgagcgtttaaacatgttctaccttatcttaaagcttttgaatcacaaggagctgacaaatctgaagggtagtcttgaaaaatatcagtctgccaataaggaagaagcgttctctgaccaattggagttaattaacctgcttaaccttgacaacaaccgcgtgacggaagacttcgaattagaggccgacgagattggaaaatttcttgatttcaatggcaacaaagttaaggataacaaggaactgaaaaagttcgatacaaacaagatctactttgacggcgagaacattatcaaacaccgtgccttctacaatattaagaaatatggcatgttaaacttactggagaaaattgccgacaaggctggatacaagatctcgatcgaagagctgaagaaatactccaataaaaagaatgagatcgagaagaaccataagatgcaggaaaatctgcaccgcaaatacgctcgtccccgtaaagacgagaagtttacagatgaggactatgaaagttacaagcaagctattgagaatattgaggagtacacccaccttaagaacaaggtagaattcaatgagctgaatttactgcagggcctgttgctgcgcattttacatcgtttagtcggatatacctcaatttgggaacgcgatctgcgcttccgccttaaaggtgagttcccagaaaaccaatacatcgaagagatcttcaactttgaaaataagaagaacgtgaagtacaaagggggtcagattgtagagaaatacattaaattctacaaggaattacatcaaaatgatgaagttaagatcaacaagtacagttccgcgaatatcaaggtgttgaagcaagaaaagaaggacctttatattcgaaattacatcgcccacttcaattatattcctcacgccgagatctcactgctggaagtccttgaaaatttgcgtaaattgctgtcctacgatcgcaaactgaaaaatgccgtaatgaaatcagtagttgatatccttaaggagtatggttttgtagccacattcaaaatcggggcggacaagaagatcggtattcagacactggagagcgaaaaaatcgtgcatcttaagaatcttaagaagaagaagttaatgactgaccgcaattccgaggaactttgcaaattggtgaagattatgtttgaatacaaaatggaagagaaaaagtctgaaaacGGCGGTGGCTCGCATCATCATCATCATCACTAAtaacattggaagtggataacgGATCCGGCTGCTAACAAAGCCCGAAAGGAAGCTGAGTTGGCTGCTGCCACCGCTGAGCAATAACTAGCATAACCCCTTGGGGCCTCTAAACGGGTCTTGAGGGGTTTTTTGCTGAAAGGAGGAACTATATCCGGATATCCCGCAAGAGGCCCGGCAGTACCGGCATAACCAAGCCTATGCCTACAGCATCCAGGGTGACGGTGCCGAGGATGACGATGAGCGCATTGTTAGATTTCATACACGGTGCCTGACTGCGTTAGCAATTTAACTGTGATAAACTACCGCATTAAAGCTTATCGATGATAAGCTGTCAAACATGAGAA

#### **pET15_LwaCas13a-6xHIS**

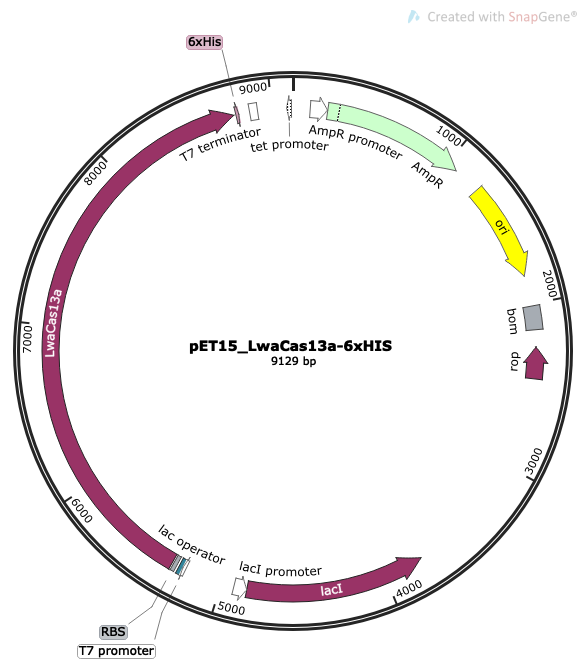

sequence:

TTCTTGAAGACGAAAGGGCCTCGTGATACGCCTATTTTTATAGGTTAATGTCATGATAATAATGGTTTCTTAGACGTCAGGTGGCACTTTTCGGGGAAATGTGCGCGGAACCCCTATTTGTTTATTTTTCTAAATACATTCAAATATGTATCCGCTCATGAGACAATAACCCTGATAAATGCTTCAATAATATTGAAAAAGGAAGAGTATGAGTATTCAACATTTCCGTGTCGCCCTTATTCCCTTTTTTGCGGCATTTTGCCTTCCTGTTTTTGCTCACCCAGAAACGCTGGTGAAAGTAAAAGATGCTGAAGATCAGTTGGGTGCACGAGTGGGTTACATCGAACTGGATCTCAACAGCGGTAAGATCCTTGAGAGTTTTCGCCCCGAAGAACGTTTTCCAATGATGAGCACTTTTAAAGTTCTGCTATGTGGCGCGGTATTATCCCGTGTTGACGCCGGGCAAGAGCAACTCGGTCGCCGCATACACTATTCTCAGAATGACTTGGTTGAGTACTCACCAGTCACAGAAAAGCATCTTACGGATGGCATGACAGTAAGAGAATTATGCAGTGCTGCCATAACCATGAGTGATAACACTGCGGCCAACTTACTTCTGACAACGATCGGAGGACCGAAGGAGCTAACCGCTTTTTTGCACAACATGGGGGATCATGTAACTCGCCTTGATCGTTGGGAACCGGAGCTGAATGAAGCCATACCAAACGACGAGCGTGACACCACGATGCCTGCAGCAATGGCAACAACGTTGCGCAAACTATTAACTGGCGAACTACTTACTCTAGCTTCCCGGCAACAATTAATAGACTGGATGGAGGCGGATAAAGTTGCAGGACCACTTCTGCGCTCGGCCCTTCCGGCTGGCTGGTTTATTGCTGATAAATCTGGAGCCGGTGAGCGTGGGTCTCGCGGTATCATTGCAGCACTGGGGCCAGATGGTAAGCCCTCCCGTATCGTAGTTATCTACACGACGGGGAGTCAGGCAACTATGGATGAACGAAATAGACAGATCGCTGAGATAGGTGCCTCACTGATTAAGCATTGGTAACTGTCAGACCAAGTTTACTCATATATACTTTAGATTGATTTAAAACTTCATTTTTAATTTAAAAGGATCTAGGTGAAGATCCTTTTTGATAATCTCATGACCAAAATCCCTTAACGTGAGTTTTCGTTCCACTGAGCGTCAGACCCCGTAGAAAAGATCAAAGGATCTTCTTGAGATCCTTTTTTTCTGCGCGTAATCTGCTGCTTGCAAACAAAAAAACCACCGCTACCAGCGGTGGTTTGTTTGCCGGATCAAGAGCTACCAACTCTTTTTCCGAAGGTAACTGGCTTCAGCAGAGCGCAGATACCAAATACTGTCCTTCTAGTGTAGCCGTAGTTAGGCCACCACTTCAAGAACTCTGTAGCACCGCCTACATACCTCGCTCTGCTAATCCTGTTACCAGTGGCTGCTGCCAGTGGCGATAAGTCGTGTCTTACCGGGTTGGACTCAAGACGATAGTTACCGGATAAGGCGCAGCGGTCGGGCTGAACGGGGGGTTCGTGCACACAGCCCAGCTTGGAGCGAACGACCTACACCGAACTGAGATACCTACAGCGTGAGCTATGAGAAAGCGCCACGCTTCCCGAAGGGAGAAAGGCGGACAGGTATCCGGTAAGCGGCAGGGTCGGAACAGGAGAGCGCACGAGGGAGCTTCCAGGGGGAAACGCCTGGTATCTTTATAGTCCTGTCGGGTTTCGCCACCTCTGACTTGAGCGTCGATTTTTGTGATGCTCGTCAGGGGGGCGGAGCCTATGGAAAAACGCCAGCAACGCGGCCTTTTTACGGTTCCTGGCCTTTTGCTGGCCTTTTGCTCACATGTTCTTTCCTGCGTTATCCCCTGATTCTGTGGATAACCGTATTACCGCCTTTGAGTGAGCTGATACCGCTCGCCGCAGCCGAACGACCGAGCGCAGCGAGTCAGTGAGCGAGGAAGCGGAAGAGCGCCTGATGCGGTATTTTCTCCTTACGCATCTGTGCGGTATTTCACACCGCATATATGGTGCACTCTCAGTACAATCTGCTCTGATGCCGCATAGTTAAGCCAGTATACACTCCGCTATCGCTACGTGACTGGGTCATGGCTGCGCCCCGACACCCGCCAACACCCGCTGACGCGCCCTGACGGGCTTGTCTGCTCCCGGCATCCGCTTACAGACAAGCTGTGACCGTCTCCGGGAGCTGCATGTGTCAGAGGTTTTCACCGTCATCACCGAAACGCGCGAGGCAGCTGCGGTAAAGCTCATCAGCGTGGTCGTGAAGCGATTCACAGATGTCTGCCTGTTCATCCGCGTCCAGCTCGTTGAGTTTCTCCAGAAGCGTTAATGTCTGGCTTCTGATAAAGCGGGCCATGTTAAGGGCGGTTTTTTCCTGTTTGGTCACTGATGCCTCCGTGTAAGGGGGATTTCTGTTCATGGGGGTAATGATACCGATGAAACGAGAGAGGATGCTCACGATACGGGTTACTGATGATGAACATGCCCGGTTACTGGAACGTTGTGAGGGTAAACAACTGGCGGTATGGATGCGGCGGGACCAGAGAAAAATCACTCAGGGTCAATGCCAGCGCTTCGTTAATACAGATGTAGGTGTTCCACAGGGTAGCCAGCAGCATCCTGCGATGCAGATCCGGAACATAATGGTGCAGGGCGCTGACTTCCGCGTTTCCAGACTTTACGAAACACGGAAACCGAAGACCATTCATGTTGTTGCTCAGGTCGCAGACGTTTTGCAGCAGCAGTCGCTTCACGTTCGCTCGCGTATCGGTGATTCATTCTGCTAACCAGTAAGGCAACCCCGCCAGCCTAGCCGGGTCCTCAACGACAGGAGCACGATCATGCGCACCCGTGGCCAGGACCCAACGCTGCCCGAGATGCGCCGCGTGCGGCTGCTGGAGATGGCGGACGCGATGGATATGTTCTGCCAAGGGTTGGTTTGCGCATTCACAGTTCTCCGCAAGAATTGATTGGCTCCAATTCTTGGAGTGGTGAATCCGTTAGCGAGGTGCCGCCGGCTTCCATTCAGGTCGAGGTGGCCCGGCTCCATGCACCGCGACGCAACGCGGGGAGGCAGACAAGGTATAGGGCGGCGCCTACAATCCATGCCAACCCGTTCCATGTGCTCGCCGAGGCGGCATAAATCGCCGTGACGATCAGCGGTCCAGTGATCGAAGTTAGGCTGGTAAGAGCCGCGAGCGATCCTTGAAGCTGTCCCTGATGGTCGTCATCTACCTGCCTGGACAGCATGGCCTGCAACGCGGGCATCCCGATGCCGCCGGAAGCGAGAAGAATCATAATGGGGAAGGCCATCCAGCCTCGCGTCGCGAACGCCAGCAAGACGTAGCCCAGCGCGTCGGCCGCCATGCCGGCGATAATGGCCTGCTTCTCGCCGAAACGTTTGGTGGCGGGACCAGTGACGAAGGCTTGAGCGAGGGCGTGCAAGATTCCGAATACCGCAAGCGACAGGCCGATCATCGTCGCGCTCCAGCGAAAGCGGTCCTCGCCGAAAATGACCCAGAGCGCTGCCGGCACCTGTCCTACGAGTTGCATGATAAAGAAGACAGTCATAAGTGCGGCGACGATAGTCATGCCCCGCGCCCACCGGAAGGAGCTGACTGGGTTGAAGGCTCTCAAGGGCATCGGTCGAGATCCCGGTGCCTAATGAGTGAGCTAACTTACATTAATTGCGTTGCGCTCACTGCCCGCTTTCCAGTCGGGAAACCTGTCGTGCCAGCTGCATTAATGAATCGGCCAACGCGCGGGGAGAGGCGGTTTGCGTATTGGGCGCCAGGGTGGTTTTTCTTTTCACCAGTGAGACGGGCAACAGCTGATTGCCCTTCACCGCCTGGCCCTGAGAGAGTTGCAGCAAGCGGTCCACGCTGGTTTGCCCCAGCAGGCGAAAATCCTGTTTGATGGTGGTTAACGGCGGGATATAACATGAGCTGTCTTCGGTATCGTCGTATCCCACTACCGAGATATCCGCACCAACGCGCAGCCCGGACTCGGTAATGGCGCGCATTGCGCCCAGCGCCATCTGATCGTTGGCAACCAGCATCGCAGTGGGAACGATGCCCTCATTCAGCATTTGCATGGTTTGTTGAAAACCGGACATGGCACTCCAGTCGCCTTCCCGTTCCGCTATCGGCTGAATTTGATTGCGAGTGAGATATTTATGCCAGCCAGCCAGACGCAGACGCGCCGAGACAGAACTTAATGGGCCCGCTAACAGCGCGATTTGCTGGTGACCCAATGCGACCAGATGCTCCACGCCCAGTCGCGTACCGTCTTCATGGGAGAAAATAATACTGTTGATGGGTGTCTGGTCAGAGACATCAAGAAATAACGCCGGAACATTAGTGCAGGCAGCTTCCACAGCAATGGCATCCTGGTCATCCAGCGGATAGTTAATGATCAGCCCACTGACGCGTTGCGCGAGAAGATTGTGCACCGCCGCTTTACAGGCTTCGACGCCGCTTCGTTCTACCATCGACACCACCACGCTGGCACCCAGTTGATCGGCGCGAGATTTAATCGCCGCGACAATTTGCGACGGCGCGTGCAGGGCCAGACTGGAGGTGGCAACGCCAATCAGCAACGACTGTTTGCCCGCCAGTTGTTGTGCCACGCGGTTGGGAATGTAATTCAGCTCCGCCATCGCCGCTTCCACTTTTTCCCGCGTTTTCGCAGAAACGTGGCTGGCCTGGTTCACCACGCGGGAAACGGTCTGATAAGAGACACCGGCATACTCTGCGACATCGTATAACGTTACTGGTTTCACATTCACCACCCTGAATTGACTCTCTTCCGGGCGCTATCATGCCATACCGCGAAAGGTTTTGCGCCATTCGATGGTGTCCGGGATCTCGACGCTCTCCCTTATGCGACTCCTGCATTAGGAAGCAGCCCAGTAGTAGGTTGAGGCCGTTGAGCACCGCCGCCGCAAGGAATGGTGCATGCAAGGAGATGGCGCCCAACAGTCCCCCGGCCACGGGGCCTGCCACCATACCCACGCCGAAACAAGCGCTCATGAGCCCGAAGTGGCGAGCCCGATCTTCCCCATCGGTGATGTCGGCGATATAGGCGCCAGCAACCGCACCTGTGGCGCCGGTGATGCCGGCCACGATGCGTCCGGCGTAGAGGATCGAGATCTCGATCCCGCGAAATTAATACGACTCACTATAGGGGAATTGTGAGCGGATAACAATTCCCCTCTAGAAATAATTTTGTTTAACTTTAAGAAGGAGATATACCATGaaagtgaccaaggtcgacggcatcagccacaagaagtacatcgaagagggcaagctcgtgaagtccaccagcgaggaaaaccggaccagcgagagactgagcgagctgctgagcatccggctggacatctacatcaagaaccccgacaacgcctccgaggaagagaaccggatcagaagagagaacctgaagaagttctttagcaacaaggtgctgcacctgaaggacagcgtgctgtatctgaagaaccggaaagaaaagaacgccgtgcaggacaagaactatagcgaagaggacatcagcgagtacgacctgaaaaacaagaacagcttctccgtgctgaagaagatcctgctgaacgaggacgtgaactctgaggaactggaaatctttcggaaggacgtggaagccaagctgaacaagatcaacagcctgaagtacagcttcgaagagaacaaggccaactaccagaagatcaacgagaacaacgtggaaaaagtgggcggcaagagcaagcggaacatcatctacgactactacagagagagcgccaagcgcaacgactacatcaacaacgtgcaggaagccttcgacaagctgtataagaaagaggatatcgagaaactgtttttcctgatcgagaacagcaagaagcacgagaagtacaagatccgcgagtactatcacaagatcatcggccggaagaacgacaaagagaacttcgccaagattatctacgaagagatccagaacgtgaacaacatcaaagagctgattgagaagatccccgacatgtctgagctgaagaaaagccaggtgttctacaagtactacctggacaaagaggaactgaacgacaagaatattaagtacgccttctgccacttcgtggaaatcgagatgtcccagctgctgaaaaactacgtgtacaagcggctgagcaacatcagcaacgataagatcaagcggatcttcgagtaccagaatctgaaaaagctgatcgaaaacaaactgctgaacaagctggacacctacgtgcggaactgcggcaagtacaactactatctgcaagtgggcgagatcgccacctccgactttatcgcccggaaccggcagaacgaggccttcctgagaaacatcatcggcgtgtccagcgtggcctacttcagcctgaggaacatcctggaaaccgagaacgagaacggtatcaccggccggatgcggggcaagaccgtgaagaacaacaagggcgaagagaaatacgtgtccggcgaggtggacaagatctacaatgagaacaagcagaacgaagtgaaagaaaatctgaagatgttctacagctacgacttcaacatggacaacaagaacgagatcgaggacttcttcgccaacatcgacgaggccatcagcagcatcagacacggcatcgtgcacttcaacctggaactggaaggcaaggacatcttcgccttcaagaatatcgcccccagcgagatctccaagaagatgtttcagaacgaaatcaacgaaaagaagctgaagctgaaaatcttcaagcagctgaacagcgccaacgtgttcaactactacgagaaggatgtgatcatcaagtacctgaagaataccaagttcaacttcgtgaacaaaaacatccccttcgtgcccagcttcaccaagctgtacaacaagattgaggacctgcggaataccctgaagtttttttggagcgtgcccaaggacaaagaagagaaggacgcccagatctacctgctgaagaatatctactacggcgagttcctgaacaagttcgtgaaaaactccaaggtgttctttaagatcaccaatgaagtgatcaagattaacaagcagcggaaccagaaaaccggccactacaagtatcagaagttcgagaacatcgagaaaaccgtgcccgtggaatacctggccatcatccagagcagagagatgatcaacaaccaggacaaagaggaaaagaatacctacatcgactttattcagcagattttcctgaagggcttcatcgactacctgaacaagaacaatctgaagtatatcgagagcaacaacaacaatgacaacaacgacatcttctccaagatcaagatcaaaaaggataacaaagagaagtacgacaagatcctgaagaactatgagaagcacaatcggaacaaagaaatccctcacgagatcaatgagttcgtgcgcgagatcaagctggggaagattctgaagtacaccgagaatctgaacatgttttacctgatcctgaagctgctgaaccacaaagagctgaccaacctgaagggcagcctggaaaagtaccagtccgccaacaaagaagaaaccttcagcgacgagttggaactgatcaacctgctgaacctggacaacaacagagtgaccgaggacttcgagctggaagccaacgagatcggcaagttcctggacttcaacgaaaacaaaatcaaggaccggaaagagctgaaaaagttcgacaccaacaagatctatttcgacggcgagaacatcatcaagcaccgggccttctacaatatcaagaaatacggcatgctgaatctgctggaaaagatcgccgataaggccaagtataagatcagcctgaaagaactgaaagagtacagcaacaagaagaatgagattgaaaagaactacaccatgcagcagaacctgcaccggaagtacgccagacccaagaaggacgaaaagttcaacgacgaggactacaaagagtatgagaaggccatcggcaacatccagaagtacacccacctgaagaacaaggtggaattcaatgagctgaacctgctgcagggcctgctgctgaagatcctgcaccggctcgtgggctacaccagcatctgggagcgggacctgagattccggctgaagggcgagtttcccgagaaccactacatcgaggaaattttcaatttcgacaactccaagaatgtgaagtacaaaagcggccagatcgtggaaaagtatatcaacttctacaaagaactgtacaaggacaatgtggaaaagcggagcatctactccgacaagaaagtgaagaaactgaagcaggaaaaaaaggacctgtacatccggaactacattgcccacttcaactacatcccccacgccgagattagcctgctggaagtgctggaaaacctgcggaagctgctgtcctacgaccggaagctgaagaacgccatcatgaagtccatcgtggacattctgaaagaatacggcttcgtggccaccttcaagatcggcgctgacaagaagatcgaaatccagaccctggaatcagagaagatcgtgcacctgaagaatctgaagaaaaagaaactgatgaccgaccggaacagcgaggaactgtgcgaactcgtgaaagtcatgttcgagtacaaggccctggaaGGCGGTGGCTCGCATCATCATCATCATCACTAAGGATCCGGCTGCTAACAAAGCCCGAAAGGAAGCTGAGTTGGCTGCTGCCACCGCTGAGCAATAACTAGCATAACCCCTTGGGGCCTCTAAACGGGTCTTGAGGGGTTTTTTGCTGAAAGGAGGAACTATATCCGGATATCCCGCAAGAGGCCCGGCAGTACCGGCATAACCAAGCCTATGCCTACAGCATCCAGGGTGACGGTGCCGAGGATGACGATGAGCGCATTGTTAGATTTCATACACGGTGCCTGACTGCGTTAGCAATTTAACTGTGATAAACTACCGCATTAAAGCTTATCGATGATAAGCTGTCAAACATGAGAA

#### **pET15_LshCas13a-6xHIS**

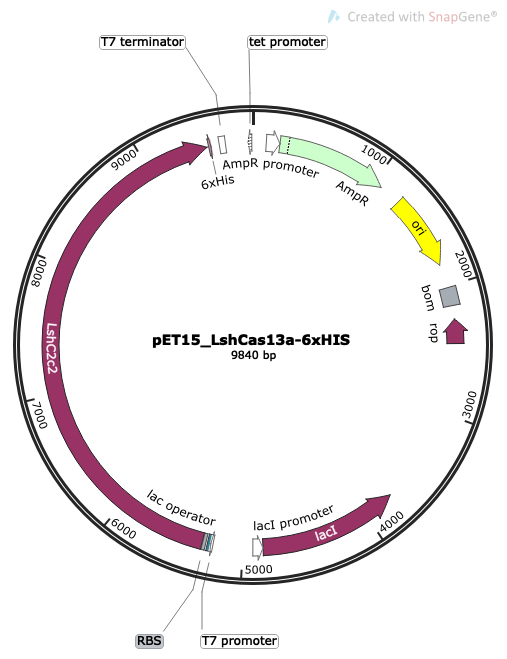

Sequence:

TTCTTGAAGACGAAAGGGCCTCGTGATACGCCTATTTTTATAGGTTAATGTCATGATAATAATGGTTTCTTAGACGTCAGGTGGCACTTTTCGGGGAAATGTGCGCGGAACCCCTATTTGTTTATTTTTCTAAATACATTCAAATATGTATCCGCTCATGAGACAATAACCCTGATAAATGCTTCAATAATATTGAAAAAGGAAGAGTATGAGTATTCAACATTTCCGTGTCGCCCTTATTCCCTTTTTTGCGGCATTTTGCCTTCCTGTTTTTGCTCACCCAGAAACGCTGGTGAAAGTAAAAGATGCTGAAGATCAGTTGGGTGCACGAGTGGGTTACATCGAACTGGATCTCAACAGCGGTAAGATCCTTGAGAGTTTTCGCCCCGAAGAACGTTTTCCAATGATGAGCACTTTTAAAGTTCTGCTATGTGGCGCGGTATTATCCCGTGTTGACGCCGGGCAAGAGCAACTCGGTCGCCGCATACACTATTCTCAGAATGACTTGGTTGAGTACTCACCAGTCACAGAAAAGCATCTTACGGATGGCATGACAGTAAGAGAATTATGCAGTGCTGCCATAACCATGAGTGATAACACTGCGGCCAACTTACTTCTGACAACGATCGGAGGACCGAAGGAGCTAACCGCTTTTTTGCACAACATGGGGGATCATGTAACTCGCCTTGATCGTTGGGAACCGGAGCTGAATGAAGCCATACCAAACGACGAGCGTGACACCACGATGCCTGCAGCAATGGCAACAACGTTGCGCAAACTATTAACTGGCGAACTACTTACTCTAGCTTCCCGGCAACAATTAATAGACTGGATGGAGGCGGATAAAGTTGCAGGACCACTTCTGCGCTCGGCCCTTCCGGCTGGCTGGTTTATTGCTGATAAATCTGGAGCCGGTGAGCGTGGGTCTCGCGGTATCATTGCAGCACTGGGGCCAGATGGTAAGCCCTCCCGTATCGTAGTTATCTACACGACGGGGAGTCAGGCAACTATGGATGAACGAAATAGACAGATCGCTGAGATAGGTGCCTCACTGATTAAGCATTGGTAACTGTCAGACCAAGTTTACTCATATATACTTTAGATTGATTTAAAACTTCATTTTTAATTTAAAAGGATCTAGGTGAAGATCCTTTTTGATAATCTCATGACCAAAATCCCTTAACGTGAGTTTTCGTTCCACTGAGCGTCAGACCCCGTAGAAAAGATCAAAGGATCTTCTTGAGATCCTTTTTTTCTGCGCGTAATCTGCTGCTTGCAAACAAAAAAACCACCGCTACCAGCGGTGGTTTGTTTGCCGGATCAAGAGCTACCAACTCTTTTTCCGAAGGTAACTGGCTTCAGCAGAGCGCAGATACCAAATACTGTCCTTCTAGTGTAGCCGTAGTTAGGCCACCACTTCAAGAACTCTGTAGCACCGCCTACATACCTCGCTCTGCTAATCCTGTTACCAGTGGCTGCTGCCAGTGGCGATAAGTCGTGTCTTACCGGGTTGGACTCAAGACGATAGTTACCGGATAAGGCGCAGCGGTCGGGCTGAACGGGGGGTTCGTGCACACAGCCCAGCTTGGAGCGAACGACCTACACCGAACTGAGATACCTACAGCGTGAGCTATGAGAAAGCGCCACGCTTCCCGAAGGGAGAAAGGCGGACAGGTATCCGGTAAGCGGCAGGGTCGGAACAGGAGAGCGCACGAGGGAGCTTCCAGGGGGAAACGCCTGGTATCTTTATAGTCCTGTCGGGTTTCGCCACCTCTGACTTGAGCGTCGATTTTTGTGATGCTCGTCAGGGGGGCGGAGCCTATGGAAAAACGCCAGCAACGCGGCCTTTTTACGGTTCCTGGCCTTTTGCTGGCCTTTTGCTCACATGTTCTTTCCTGCGTTATCCCCTGATTCTGTGGATAACCGTATTACCGCCTTTGAGTGAGCTGATACCGCTCGCCGCAGCCGAACGACCGAGCGCAGCGAGTCAGTGAGCGAGGAAGCGGAAGAGCGCCTGATGCGGTATTTTCTCCTTACGCATCTGTGCGGTATTTCACACCGCATATATGGTGCACTCTCAGTACAATCTGCTCTGATGCCGCATAGTTAAGCCAGTATACACTCCGCTATCGCTACGTGACTGGGTCATGGCTGCGCCCCGACACCCGCCAACACCCGCTGACGCGCCCTGACGGGCTTGTCTGCTCCCGGCATCCGCTTACAGACAAGCTGTGACCGTCTCCGGGAGCTGCATGTGTCAGAGGTTTTCACCGTCATCACCGAAACGCGCGAGGCAGCTGCGGTAAAGCTCATCAGCGTGGTCGTGAAGCGATTCACAGATGTCTGCCTGTTCATCCGCGTCCAGCTCGTTGAGTTTCTCCAGAAGCGTTAATGTCTGGCTTCTGATAAAGCGGGCCATGTTAAGGGCGGTTTTTTCCTGTTTGGTCACTGATGCCTCCGTGTAAGGGGGATTTCTGTTCATGGGGGTAATGATACCGATGAAACGAGAGAGGATGCTCACGATACGGGTTACTGATGATGAACATGCCCGGTTACTGGAACGTTGTGAGGGTAAACAACTGGCGGTATGGATGCGGCGGGACCAGAGAAAAATCACTCAGGGTCAATGCCAGCGCTTCGTTAATACAGATGTAGGTGTTCCACAGGGTAGCCAGCAGCATCCTGCGATGCAGATCCGGAACATAATGGTGCAGGGCGCTGACTTCCGCGTTTCCAGACTTTACGAAACACGGAAACCGAAGACCATTCATGTTGTTGCTCAGGTCGCAGACGTTTTGCAGCAGCAGTCGCTTCACGTTCGCTCGCGTATCGGTGATTCATTCTGCTAACCAGTAAGGCAACCCCGCCAGCCTAGCCGGGTCCTCAACGACAGGAGCACGATCATGCGCACCCGTGGCCAGGACCCAACGCTGCCCGAGATGCGCCGCGTGCGGCTGCTGGAGATGGCGGACGCGATGGATATGTTCTGCCAAGGGTTGGTTTGCGCATTCACAGTTCTCCGCAAGAATTGATTGGCTCCAATTCTTGGAGTGGTGAATCCGTTAGCGAGGTGCCGCCGGCTTCCATTCAGGTCGAGGTGGCCCGGCTCCATGCACCGCGACGCAACGCGGGGAGGCAGACAAGGTATAGGGCGGCGCCTACAATCCATGCCAACCCGTTCCATGTGCTCGCCGAGGCGGCATAAATCGCCGTGACGATCAGCGGTCCAGTGATCGAAGTTAGGCTGGTAAGAGCCGCGAGCGATCCTTGAAGCTGTCCCTGATGGTCGTCATCTACCTGCCTGGACAGCATGGCCTGCAACGCGGGCATCCCGATGCCGCCGGAAGCGAGAAGAATCATAATGGGGAAGGCCATCCAGCCTCGCGTCGCGAACGCCAGCAAGACGTAGCCCAGCGCGTCGGCCGCCATGCCGGCGATAATGGCCTGCTTCTCGCCGAAACGTTTGGTGGCGGGACCAGTGACGAAGGCTTGAGCGAGGGCGTGCAAGATTCCGAATACCGCAAGCGACAGGCCGATCATCGTCGCGCTCCAGCGAAAGCGGTCCTCGCCGAAAATGACCCAGAGCGCTGCCGGCACCTGTCCTACGAGTTGCATGATAAAGAAGACAGTCATAAGTGCGGCGACGATAGTCATGCCCCGCGCCCACCGGAAGGAGCTGACTGGGTTGAAGGCTCTCAAGGGCATCGGTCGAGATCCCGGTGCCTAATGAGTGAGCTAACTTACATTAATTGCGTTGCGCTCACTGCCCGCTTTCCAGTCGGGAAACCTGTCGTGCCAGCTGCATTAATGAATCGGCCAACGCGCGGGGAGAGGCGGTTTGCGTATTGGGCGCCAGGGTGGTTTTTCTTTTCACCAGTGAGACGGGCAACAGCTGATTGCCCTTCACCGCCTGGCCCTGAGAGAGTTGCAGCAAGCGGTCCACGCTGGTTTGCCCCAGCAGGCGAAAATCCTGTTTGATGGTGGTTAACGGCGGGATATAACATGAGCTGTCTTCGGTATCGTCGTATCCCACTACCGAGATATCCGCACCAACGCGCAGCCCGGACTCGGTAATGGCGCGCATTGCGCCCAGCGCCATCTGATCGTTGGCAACCAGCATCGCAGTGGGAACGATGCCCTCATTCAGCATTTGCATGGTTTGTTGAAAACCGGACATGGCACTCCAGTCGCCTTCCCGTTCCGCTATCGGCTGAATTTGATTGCGAGTGAGATATTTATGCCAGCCAGCCAGACGCAGACGCGCCGAGACAGAACTTAATGGGCCCGCTAACAGCGCGATTTGCTGGTGACCCAATGCGACCAGATGCTCCACGCCCAGTCGCGTACCGTCTTCATGGGAGAAAATAATACTGTTGATGGGTGTCTGGTCAGAGACATCAAGAAATAACGCCGGAACATTAGTGCAGGCAGCTTCCACAGCAATGGCATCCTGGTCATCCAGCGGATAGTTAATGATCAGCCCACTGACGCGTTGCGCGAGAAGATTGTGCACCGCCGCTTTACAGGCTTCGACGCCGCTTCGTTCTACCATCGACACCACCACGCTGGCACCCAGTTGATCGGCGCGAGATTTAATCGCCGCGACAATTTGCGACGGCGCGTGCAGGGCCAGACTGGAGGTGGCAACGCCAATCAGCAACGACTGTTTGCCCGCCAGTTGTTGTGCCACGCGGTTGGGAATGTAATTCAGCTCCGCCATCGCCGCTTCCACTTTTTCCCGCGTTTTCGCAGAAACGTGGCTGGCCTGGTTCACCACGCGGGAAACGGTCTGATAAGAGACACCGGCATACTCTGCGACATCGTATAACGTTACTGGTTTCACATTCACCACCCTGAATTGACTCTCTTCCGGGCGCTATCATGCCATACCGCGAAAGGTTTTGCGCCATTCGATGGTGTCCGGGATCTCGACGCTCTCCCTTATGCGACTCCTGCATTAGGAAGCAGCCCAGTAGTAGGTTGAGGCCGTTGAGCACCGCCGCCGCAAGGAATGGTGCATGCAAGGAGATGGCGCCCAACAGTCCCCCGGCCACGGGGCCTGCCACCATACCCACGCCGAAACAAGCGCTCATGAGCCCGAAGTGGCGAGCCCGATCTTCCCCATCGGTGATGTCGGCGATATAGGCGCCAGCAACCGCACCTGTGGCGCCGGTGATGCCGGCCACGATGCGTCCGGCGTAGAGGATCGAGATCTCGATCCCGCGAAATTAATACGACTCACTATAGGGGAATTGTGAGCGGATAACAATTCCCCTCTAGAAATAATTTTGTTTAACTTTAAGAAGGAGATATACCatgaaagtgacgaaggtaggaggcatttcgcataagaagtacacgtccgaaggccgcttagtgaagtcagaatcggaagaaaatcgcacagacgaacgtctgtcggcgttgcttaatatgcgccttgacatgtatatcaagaatcccagcagcacggaaaccaaggaaaatcaaaaacgcattgggaaattaaagaaattcttctcaaacaaaatggtctatcttaaagacaataccttgagtttgaagaatgggaaaaaggagaacattgatcgtgagtattctgagactgacatccttgagagcgatgtccgtgacaagaaaaacttcgccgtgttgaaaaagatctatctgaatgaaaacgtgaactcggaggaattggaagtttttcgtaacgacattaagaagaaactgaacaaaatcaacagcctgaagtactcatttgaaaagaataaggcgaattatcaaaagattaatgagaataacatcgagaaggttgaaggtaagtcaaagcgtaacattatttacgattattatcgtgagtcagcgaaacgtgacgcttatgtaagcaatgtgaaagaagcctttgataagctttacaaggaagaggacattgcaaaacttgttcttgaaattgagaaccttacgaagttagagaaatacaagattcgcgagttctaccacgaaattattggacgtaagaatgacaaggaaaactttgcaaaaatcatctacgaagaaatccagaatgttaataacatgaaagagttgatcgagaaggtaccggacatgagtgagttgaaaaagagccaagtattttacaagtattacttagacaaagaagagttgaacgacaagaacatcaaatacgcgttttgtcatttcgtggaaatcgaaatgagtcagttgctgaagaactacgtatataagcgcttaagtaatatctcgaatgacaaaattaagcgtatctttgaataccagaacttgaaaaaattgatcgaaaataagctgttaaacaaacttgacacgtacgtccgtaattgtggaaagtataattattatttgcaagacggcgaaattgccacttcagatttcatcgcccgcaaccgtcagaatgaagcgtttcttcgcaacatcattggggtgtcatctgtggcctacttttctcttcgcaacattcttgaaacggagaacgagaatgatattactgggcgtatgcgcggcaaaacagttaagaacaataaaggtgaagagaagtacgtgtccggagaagttgataagatctataatgaaaataagaagaacgaggttaaggagaacttaaaaatgttctattcgtacgatttcaatatggacaacaagaatgaaatcgaagatttcttcgccaacatcgacgaggcgatttcttccatccgtcacggtattgtccacttcaacttggaattagaaggtaaggatatctttgcgttcaagaacattgcgccatccgaaatctcaaagaagatgtttcagaatgagattaacgagaaaaaactgaaattgaagatctttcgtcaactgaactctgccaacgtgttccgctatctcgaaaagtataaaattctgaattaccttaaacgtacacgcttcgagtttgtcaataaaaatatcccattcgtcccgtctttcaccaaattatattcgcgcattgatgacctgaagaatagtcttgggatttactggaaaactccgaaaacaaacgacgacaataagactaaggagattattgatgcccaaatctatttgcttaaaaacatctattacggggagttcctgaattatttcatgtcgaacaatggtaatttctttgagatttctaaagaaatcatcgaattgaacaagaacgataaacgcaacttaaagactgggttttacaagctgcaaaagtttgaagacatccaggagaagattccaaaggaatacttggcgaatatccagtccctgtacatgattaatgccggtaatcaggacgaagaagaaaaggacacttatattgatttcattcaaaagatcttcttaaagggatttatgacgtatcttgctaataacggtcgtttaagtctgatttacatcggctcggatgaagaaacaaatacgtcattagcagaaaagaagcaagagtttgacaagttcttgaagaagtacgagcagaacaataatatcaagatcccctatgagatcaatgaattcctgcgtgagatcaaactgggaaacatcctgaagtatactgagcgtttaaacatgttctaccttatcttaaagcttttgaatcacaaggagctgacaaatctgaagggtagtcttgaaaaatatcagtctgccaataaggaagaagcgttctctgaccaattggagttaattaacctgcttaaccttgacaacaaccgcgtgacggaagacttcgaattagaggccgacgagattggaaaatttcttgatttcaatggcaacaaagttaaggataacaaggaactgaaaaagttcgatacaaacaagatctactttgacggcgagaacattatcaaacaccgtgccttctacaatattaagaaatatggcatgttaaacttactggagaaaattgccgacaaggctggatacaagatctcgatcgaagagctgaagaaatactccaataaaaagaatgagatcgagaagaaccataagatgcaggaaaatctgcaccgcaaatacgctcgtccccgtaaagacgagaagtttacagatgaggactatgaaagttacaagcaagctattgagaatattgaggagtacacccaccttaagaacaaggtagaattcaatgagctgaatttactgcagggcctgttgctgcgcattttacatcgtttagtcggatatacctcaatttgggaacgcgatctgcgcttccgccttaaaggtgagttcccagaaaaccaatacatcgaagagatcttcaactttgaaaataagaagaacgtgaagtacaaagggggtcagattgtagagaaatacattaaattctacaaggaattacatcaaaatgatgaagttaagatcaacaagtacagttccgcgaatatcaaggtgttgaagcaagaaaagaaggacctttatattcgaaattacatcgcccacttcaattatattcctcacgccgagatctcactgctggaagtccttgaaaatttgcgtaaattgctgtcctacgatcgcaaactgaaaaatgccgtaatgaaatcagtagttgatatccttaaggagtatggttttgtagccacattcaaaatcggggcggacaagaagatcggtattcagacactggagagcgaaaaaatcgtgcatcttaagaatcttaagaagaagaagttaatgactgaccgcaattccgaggaactttgcaaattggtgaagattatgtttgaatacaaaatggaagagaaaaagtctgaaaacGGCGGTGGCTCGCATCATCATCATCATCACTAAtaacattggaagtggataacgGATCCGGCTGCTAACAAAGCCCGAAAGGAAGCTGAGTTGGCTGCTGCCACCGCTGAGCAATAACTAGCATAACCCCTTGGGGCCTCTAAACGGGTCTTGAGGGGTTTTTTGCTGAAAGGAGGAACTATATCCGGATATCCCGCAAGAGGCCCGGCAGTACCGGCATAACCAAGCCTATGCCTACAGCATCCAGGGTGACGGTGCCGAGGATGACGATGAGCGCATTGTTAGATTTCATACACGGTGCCTGACTGCGTTAGCAATTTAACTGTGATAAACTACCGCATTAAAGCTTATCGATGATAAGCTGTCAAACATGAGAA

#### **pET15_PspCas13b-6xHIS**

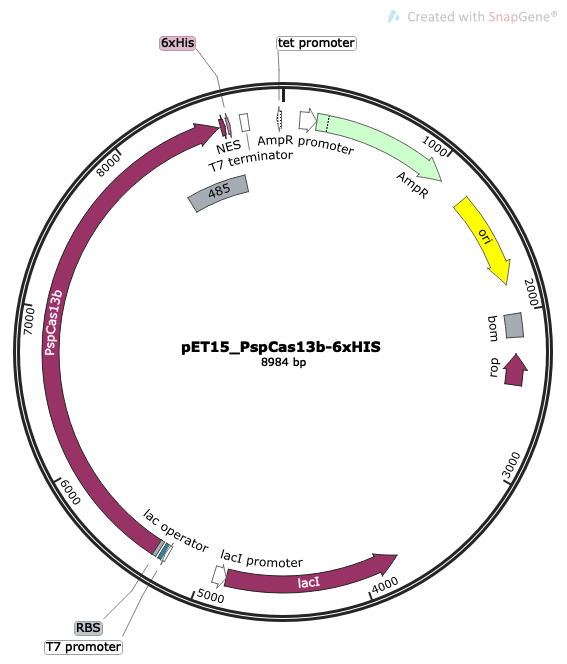

Sequence:

TTCTTGAAGACGAAAGGGCCTCGTGATACGCCTATTTTTATAGGTTAATGTCATGATAATAATGGTTTCTTAGACGTCAGGTGGCACTTTTCGGGGAAATGTGCGCGGAACCCCTATTTGTTTATTTTTCTAAATACATTCAAATATGTATCCGCTCATGAGACAATAACCCTGATAAATGCTTCAATAATATTGAAAAAGGAAGAGTATGAGTATTCAACATTTCCGTGTCGCCCTTATTCCCTTTTTTGCGGCATTTTGCCTTCCTGTTTTTGCTCACCCAGAAACGCTGGTGAAAGTAAAAGATGCTGAAGATCAGTTGGGTGCACGAGTGGGTTACATCGAACTGGATCTCAACAGCGGTAAGATCCTTGAGAGTTTTCGCCCCGAAGAACGTTTTCCAATGATGAGCACTTTTAAAGTTCTGCTATGTGGCGCGGTATTATCCCGTGTTGACGCCGGGCAAGAGCAACTCGGTCGCCGCATACACTATTCTCAGAATGACTTGGTTGAGTACTCACCAGTCACAGAAAAGCATCTTACGGATGGCATGACAGTAAGAGAATTATGCAGTGCTGCCATAACCATGAGTGATAACACTGCGGCCAACTTACTTCTGACAACGATCGGAGGACCGAAGGAGCTAACCGCTTTTTTGCACAACATGGGGGATCATGTAACTCGCCTTGATCGTTGGGAACCGGAGCTGAATGAAGCCATACCAAACGACGAGCGTGACACCACGATGCCTGCAGCAATGGCAACAACGTTGCGCAAACTATTAACTGGCGAACTACTTACTCTAGCTTCCCGGCAACAATTAATAGACTGGATGGAGGCGGATAAAGTTGCAGGACCACTTCTGCGCTCGGCCCTTCCGGCTGGCTGGTTTATTGCTGATAAATCTGGAGCCGGTGAGCGTGGGTCTCGCGGTATCATTGCAGCACTGGGGCCAGATGGTAAGCCCTCCCGTATCGTAGTTATCTACACGACGGGGAGTCAGGCAACTATGGATGAACGAAATAGACAGATCGCTGAGATAGGTGCCTCACTGATTAAGCATTGGTAACTGTCAGACCAAGTTTACTCATATATACTTTAGATTGATTTAAAACTTCATTTTTAATTTAAAAGGATCTAGGTGAAGATCCTTTTTGATAATCTCATGACCAAAATCCCTTAACGTGAGTTTTCGTTCCACTGAGCGTCAGACCCCGTAGAAAAGATCAAAGGATCTTCTTGAGATCCTTTTTTTCTGCGCGTAATCTGCTGCTTGCAAACAAAAAAACCACCGCTACCAGCGGTGGTTTGTTTGCCGGATCAAGAGCTACCAACTCTTTTTCCGAAGGTAACTGGCTTCAGCAGAGCGCAGATACCAAATACTGTCCTTCTAGTGTAGCCGTAGTTAGGCCACCACTTCAAGAACTCTGTAGCACCGCCTACATACCTCGCTCTGCTAATCCTGTTACCAGTGGCTGCTGCCAGTGGCGATAAGTCGTGTCTTACCGGGTTGGACTCAAGACGATAGTTACCGGATAAGGCGCAGCGGTCGGGCTGAACGGGGGGTTCGTGCACACAGCCCAGCTTGGAGCGAACGACCTACACCGAACTGAGATACCTACAGCGTGAGCTATGAGAAAGCGCCACGCTTCCCGAAGGGAGAAAGGCGGACAGGTATCCGGTAAGCGGCAGGGTCGGAACAGGAGAGCGCACGAGGGAGCTTCCAGGGGGAAACGCCTGGTATCTTTATAGTCCTGTCGGGTTTCGCCACCTCTGACTTGAGCGTCGATTTTTGTGATGCTCGTCAGGGGGGCGGAGCCTATGGAAAAACGCCAGCAACGCGGCCTTTTTACGGTTCCTGGCCTTTTGCTGGCCTTTTGCTCACATGTTCTTTCCTGCGTTATCCCCTGATTCTGTGGATAACCGTATTACCGCCTTTGAGTGAGCTGATACCGCTCGCCGCAGCCGAACGACCGAGCGCAGCGAGTCAGTGAGCGAGGAAGCGGAAGAGCGCCTGATGCGGTATTTTCTCCTTACGCATCTGTGCGGTATTTCACACCGCATATATGGTGCACTCTCAGTACAATCTGCTCTGATGCCGCATAGTTAAGCCAGTATACACTCCGCTATCGCTACGTGACTGGGTCATGGCTGCGCCCCGACACCCGCCAACACCCGCTGACGCGCCCTGACGGGCTTGTCTGCTCCCGGCATCCGCTTACAGACAAGCTGTGACCGTCTCCGGGAGCTGCATGTGTCAGAGGTTTTCACCGTCATCACCGAAACGCGCGAGGCAGCTGCGGTAAAGCTCATCAGCGTGGTCGTGAAGCGATTCACAGATGTCTGCCTGTTCATCCGCGTCCAGCTCGTTGAGTTTCTCCAGAAGCGTTAATGTCTGGCTTCTGATAAAGCGGGCCATGTTAAGGGCGGTTTTTTCCTGTTTGGTCACTGATGCCTCCGTGTAAGGGGGATTTCTGTTCATGGGGGTAATGATACCGATGAAACGAGAGAGGATGCTCACGATACGGGTTACTGATGATGAACATGCCCGGTTACTGGAACGTTGTGAGGGTAAACAACTGGCGGTATGGATGCGGCGGGACCAGAGAAAAATCACTCAGGGTCAATGCCAGCGCTTCGTTAATACAGATGTAGGTGTTCCACAGGGTAGCCAGCAGCATCCTGCGATGCAGATCCGGAACATAATGGTGCAGGGCGCTGACTTCCGCGTTTCCAGACTTTACGAAACACGGAAACCGAAGACCATTCATGTTGTTGCTCAGGTCGCAGACGTTTTGCAGCAGCAGTCGCTTCACGTTCGCTCGCGTATCGGTGATTCATTCTGCTAACCAGTAAGGCAACCCCGCCAGCCTAGCCGGGTCCTCAACGACAGGAGCACGATCATGCGCACCCGTGGCCAGGACCCAACGCTGCCCGAGATGCGCCGCGTGCGGCTGCTGGAGATGGCGGACGCGATGGATATGTTCTGCCAAGGGTTGGTTTGCGCATTCACAGTTCTCCGCAAGAATTGATTGGCTCCAATTCTTGGAGTGGTGAATCCGTTAGCGAGGTGCCGCCGGCTTCCATTCAGGTCGAGGTGGCCCGGCTCCATGCACCGCGACGCAACGCGGGGAGGCAGACAAGGTATAGGGCGGCGCCTACAATCCATGCCAACCCGTTCCATGTGCTCGCCGAGGCGGCATAAATCGCCGTGACGATCAGCGGTCCAGTGATCGAAGTTAGGCTGGTAAGAGCCGCGAGCGATCCTTGAAGCTGTCCCTGATGGTCGTCATCTACCTGCCTGGACAGCATGGCCTGCAACGCGGGCATCCCGATGCCGCCGGAAGCGAGAAGAATCATAATGGGGAAGGCCATCCAGCCTCGCGTCGCGAACGCCAGCAAGACGTAGCCCAGCGCGTCGGCCGCCATGCCGGCGATAATGGCCTGCTTCTCGCCGAAACGTTTGGTGGCGGGACCAGTGACGAAGGCTTGAGCGAGGGCGTGCAAGATTCCGAATACCGCAAGCGACAGGCCGATCATCGTCGCGCTCCAGCGAAAGCGGTCCTCGCCGAAAATGACCCAGAGCGCTGCCGGCACCTGTCCTACGAGTTGCATGATAAAGAAGACAGTCATAAGTGCGGCGACGATAGTCATGCCCCGCGCCCACCGGAAGGAGCTGACTGGGTTGAAGGCTCTCAAGGGCATCGGTCGAGATCCCGGTGCCTAATGAGTGAGCTAACTTACATTAATTGCGTTGCGCTCACTGCCCGCTTTCCAGTCGGGAAACCTGTCGTGCCAGCTGCATTAATGAATCGGCCAACGCGCGGGGAGAGGCGGTTTGCGTATTGGGCGCCAGGGTGGTTTTTCTTTTCACCAGTGAGACGGGCAACAGCTGATTGCCCTTCACCGCCTGGCCCTGAGAGAGTTGCAGCAAGCGGTCCACGCTGGTTTGCCCCAGCAGGCGAAAATCCTGTTTGATGGTGGTTAACGGCGGGATATAACATGAGCTGTCTTCGGTATCGTCGTATCCCACTACCGAGATATCCGCACCAACGCGCAGCCCGGACTCGGTAATGGCGCGCATTGCGCCCAGCGCCATCTGATCGTTGGCAACCAGCATCGCAGTGGGAACGATGCCCTCATTCAGCATTTGCATGGTTTGTTGAAAACCGGACATGGCACTCCAGTCGCCTTCCCGTTCCGCTATCGGCTGAATTTGATTGCGAGTGAGATATTTATGCCAGCCAGCCAGACGCAGACGCGCCGAGACAGAACTTAATGGGCCCGCTAACAGCGCGATTTGCTGGTGACCCAATGCGACCAGATGCTCCACGCCCAGTCGCGTACCGTCTTCATGGGAGAAAATAATACTGTTGATGGGTGTCTGGTCAGAGACATCAAGAAATAACGCCGGAACATTAGTGCAGGCAGCTTCCACAGCAATGGCATCCTGGTCATCCAGCGGATAGTTAATGATCAGCCCACTGACGCGTTGCGCGAGAAGATTGTGCACCGCCGCTTTACAGGCTTCGACGCCGCTTCGTTCTACCATCGACACCACCACGCTGGCACCCAGTTGATCGGCGCGAGATTTAATCGCCGCGACAATTTGCGACGGCGCGTGCAGGGCCAGACTGGAGGTGGCAACGCCAATCAGCAACGACTGTTTGCCCGCCAGTTGTTGTGCCACGCGGTTGGGAATGTAATTCAGCTCCGCCATCGCCGCTTCCACTTTTTCCCGCGTTTTCGCAGAAACGTGGCTGGCCTGGTTCACCACGCGGGAAACGGTCTGATAAGAGACACCGGCATACTCTGCGACATCGTATAACGTTACTGGTTTCACATTCACCACCCTGAATTGACTCTCTTCCGGGCGCTATCATGCCATACCGCGAAAGGTTTTGCGCCATTCGATGGTGTCCGGGATCTCGACGCTCTCCCTTATGCGACTCCTGCATTAGGAAGCAGCCCAGTAGTAGGTTGAGGCCGTTGAGCACCGCCGCCGCAAGGAATGGTGCATGCAAGGAGATGGCGCCCAACAGTCCCCCGGCCACGGGGCCTGCCACCATACCCACGCCGAAACAAGCGCTCATGAGCCCGAAGTGGCGAGCCCGATCTTCCCCATCGGTGATGTCGGCGATATAGGCGCCAGCAACCGCACCTGTGGCGCCGGTGATGCCGGCCACGATGCGTCCGGCGTAGAGGATCGAGATCTCGATCCCGCGAAATTAATACGACTCACTATAGGGGAATTGTGAGCGGATAACAATTCCCCTCTAGAAATAATTTTGTTTAACTTTAAGAAGGAGATATACCATGaacatccccgctctggtggaaaaccagaagaagtactttggcacctacagcgtgatggccatgctgaacgctcagaccgtgctggaccacatccagaaggtggccgatattgagggcgagcagaacgagaacaacgagaatctgtggtttcaccccgtgatgagccacctgtacaacgccaagaacggctacgacaagcagcccgagaaaaccatgttcatcatcgagcggctgcagagctacttcccattcctgaagatcatggccgagaaccagagagagtacagcaacggcaagtacaagcagaaccgcgtggaagtgaacagcaacgacatcttcgaggtgctgaagcgcgccttcggcgtgctgaagatgtacagggacctgaccaaccactacaagacctacgaggaaaagctgaacgacggctgcgagttcctgaccagcacagagcaacctctgagcggcatgatcaacaactactacacagtggccctgcggaacatgaacgagagatacggctacaagacagaggacctggccttcatccaggacaagcggttcaagttcgtgaaggacgcctacggcaagaaaaagtcccaagtgaataccggattcttcctgagcctgcaggactacaacggcgacacacagaagaagctgcacctgagcggagtgggaatcgccctgctgatctgcctgttcctggacaagcagtacatcaacatctttctgagcaggctgcccatcttctccagctacaatgcccagagcgaggaacggcggatcatcatcagatccttcggcatcaacagcatcaagctgcccaaggaccggatccacagcgagaagtccaacaagagcgtggccatggatatgctcaacgaagtgaagcggtgccccgacgagctgttcacaacactgtctgccgagaagcagtcccggttcagaatcatcagcgacgaccacaatgaagtgctgatgaagcggagcagcgacagattcgtgcctctgctgctgcagtatatcgattacggcaagctgttcgaccacatcaggttccacgtgaacatgggcaagctgagatacctgctgaaggccgacaagacctgcatcgacggccagaccagagtcagagtgatcgagcagcccctgaacggcttcggcagactggaagaggccgagacaatgcggaagcaagagaacggcaccttcggcaacagcggcatccggatcagagacttcgagaacatgaagcgggacgacgccaatcctgccaactatccctacatcgtggacacctacacacactacatcctggaaaacaacaaggtcgagatgtttatcaacgacaaagaggacagcgccccactgctgcccgtgatcgaggatgatagatacgtggtcaagacaatccccagctgccggatgagcaccctggaaattccagccatggccttccacatgtttctgttcggcagcaagaaaaccgagaagctgatcgtggacgtgcacaaccggtacaagagactgttccaggccatgcagaaagaagaagtgaccgccgagaatatcgccagcttcggaatcgccgagagcgacctgcctcagaagatcctggatctgatcagcggcaatgcccacggcaaggatgtggacgccttcatcagactgaccgtggacgacatgctgaccgacaccgagcggagaatcaagagattcaaggacgaccggaagtccattcggagcgccgacaacaagatgggaaagagaggcttcaagcagatctccacaggcaagctggccgacttcctggccaaggacatcgtgctgtttcagcccagcgtgaacgatggcgagaacaagatcaccggcctgaactaccggatcatgcagagcgccattgccgtgtacgatagcggcgacgattacgaggccaagcagcagttcaagctgatgttcgagaaggcccggctgatcggcaagggcacaacagagcctcatccatttctgtacaaggtgttcgcccgcagcatccccgccaatgccgtcgagttctacgagcgctacctgatcgagcggaagttctacctgaccggcctgtccaacgagatcaagaaaggcaacagagtggatgtgcccttcatccggcgggaccagaacaagtggaaaacacccgccatgaagaccctgggcagaatctacagcgaggatctgcccgtggaactgcccagacagatgttcgacaatgagatcaagtcccacctgaagtccctgccacagatggaaggcatcgacttcaacaatgccaacgtgacctatctgatcgccgagtacatgaagagagtgctggacgacgacttccagaccttctaccagtggaaccgcaactaccggtacatggacatgcttaagggcgagtacgacagaaagggctccctgcagcactgcttcaccagcgtggaagagagagaaggcctctggaaagagcgggcctccagaacagagcggtacagaaagcaggccagcaacaagatccgcagcaaccggcagatgagaaacgccagcagcgaagagatcgagacaatcctggataagcggctgagcaacagccggaacgagtaccagaaaagcgagaaagtgatccggcgctacagagtgcaggatgccctgctgtttctgctggccaaaaagaccctgaccgaactggccgatttcgacggcgagaggttcaaactgaaagaaatcatgcccgacgccgagaagggaatcctgagcgagatcatgcccatgagcttcaccttcgagaaaggcggcaagaagtacaccatcaccagcgagggcatgaagctgaagaactacggcgacttctttgtgctggctagcgacaagaggatcggcaacctgctggaactcgtgggcagcgacatcgtgtccaaagaggatatcatggaagagttcaacaaatacgaccagtgcaggcccgagatcagctccatcgtgttcaacctggaaaagtgggccttcgacacataccccgagctgtctgccagagtggaccgggaagagaaggtggacttcaagagcatcctgaaaatcctgctgaacaacaagaacatcaacaaagagcagagcgacatcctgcggaagatccggaacgccttcgatcacaacaattaccccgacaaaggcgtggtggaaatcaaggccctgcctgagatcgccatgagcatcaagaaggcctttggggagtacgccatcatgaagggatcccttcaactgcctccacttgaaagactgacactgGGCGGTGGCTCGCATCATCATCATCATCACTAATAAGATCCGGCTGCTAACAAAGCCCGAAAGGAAGCTGAGTTGGCTGCTGCCACCGCTGAGCAATAACTAGCATAACCCCTTGGGGCCTCTAAACGGGTCTTGAGGGGTTTTTTGCTGAAAGGAGGAACTATATCCGGATATCCCGCAAGAGGCCCGGCAGTACCGGCATAACCAAGCCTATGCCTACAGCATCCAGGGTGACGGTGCCGAGGATGACGATGAGCGCATTGTTAGATTTCATACACGGTGCCTGACTGCGTTAGCAATTTAACTGTGATAAACTACCGCATTAAAGCTTATCGATGATAAGCTGTCAAACATGAGAA

#### **pET15_BzCas13b-6xHIS**

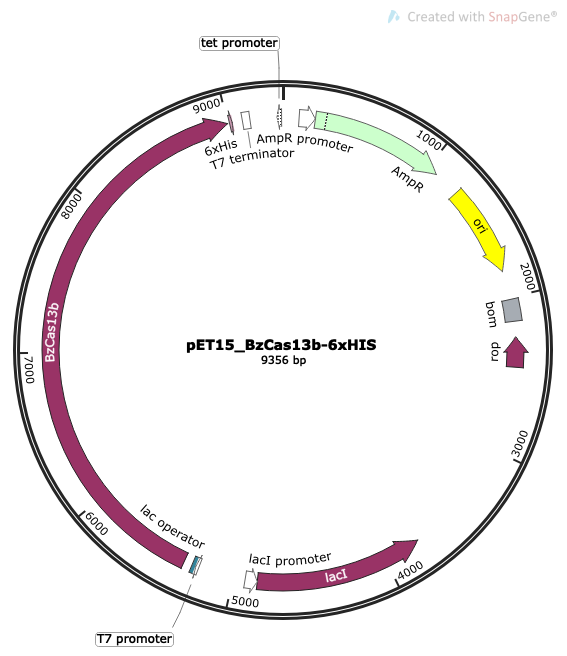

sequence:

TTCTTGAAGACGAAAGGGCCTCGTGATACGCCTATTTTTATAGGTTAATGTCATGATAATAATGGTTTCTTAGACGTCAGGTGGCACTTTTCGGGGAAATGTGCGCGGAACCCCTATTTGTTTATTTTTCTAAATACATTCAAATATGTATCCGCTCATGAGACAATAACCCTGATAAATGCTTCAATAATATTGAAAAAGGAAGAGTATGAGTATTCAACATTTCCGTGTCGCCCTTATTCCCTTTTTTGCGGCATTTTGCCTTCCTGTTTTTGCTCACCCAGAAACGCTGGTGAAAGTAAAAGATGCTGAAGATCAGTTGGGTGCACGAGTGGGTTACATCGAACTGGATCTCAACAGCGGTAAGATCCTTGAGAGTTTTCGCCCCGAAGAACGTTTTCCAATGATGAGCACTTTTAAAGTTCTGCTATGTGGCGCGGTATTATCCCGTGTTGACGCCGGGCAAGAGCAACTCGGTCGCCGCATACACTATTCTCAGAATGACTTGGTTGAGTACTCACCAGTCACAGAAAAGCATCTTACGGATGGCATGACAGTAAGAGAATTATGCAGTGCTGCCATAACCATGAGTGATAACACTGCGGCCAACTTACTTCTGACAACGATCGGAGGACCGAAGGAGCTAACCGCTTTTTTGCACAACATGGGGGATCATGTAACTCGCCTTGATCGTTGGGAACCGGAGCTGAATGAAGCCATACCAAACGACGAGCGTGACACCACGATGCCTGCAGCAATGGCAACAACGTTGCGCAAACTATTAACTGGCGAACTACTTACTCTAGCTTCCCGGCAACAATTAATAGACTGGATGGAGGCGGATAAAGTTGCAGGACCACTTCTGCGCTCGGCCCTTCCGGCTGGCTGGTTTATTGCTGATAAATCTGGAGCCGGTGAGCGTGGGTCTCGCGGTATCATTGCAGCACTGGGGCCAGATGGTAAGCCCTCCCGTATCGTAGTTATCTACACGACGGGGAGTCAGGCAACTATGGATGAACGAAATAGACAGATCGCTGAGATAGGTGCCTCACTGATTAAGCATTGGTAACTGTCAGACCAAGTTTACTCATATATACTTTAGATTGATTTAAAACTTCATTTTTAATTTAAAAGGATCTAGGTGAAGATCCTTTTTGATAATCTCATGACCAAAATCCCTTAACGTGAGTTTTCGTTCCACTGAGCGTCAGACCCCGTAGAAAAGATCAAAGGATCTTCTTGAGATCCTTTTTTTCTGCGCGTAATCTGCTGCTTGCAAACAAAAAAACCACCGCTACCAGCGGTGGTTTGTTTGCCGGATCAAGAGCTACCAACTCTTTTTCCGAAGGTAACTGGCTTCAGCAGAGCGCAGATACCAAATACTGTCCTTCTAGTGTAGCCGTAGTTAGGCCACCACTTCAAGAACTCTGTAGCACCGCCTACATACCTCGCTCTGCTAATCCTGTTACCAGTGGCTGCTGCCAGTGGCGATAAGTCGTGTCTTACCGGGTTGGACTCAAGACGATAGTTACCGGATAAGGCGCAGCGGTCGGGCTGAACGGGGGGTTCGTGCACACAGCCCAGCTTGGAGCGAACGACCTACACCGAACTGAGATACCTACAGCGTGAGCTATGAGAAAGCGCCACGCTTCCCGAAGGGAGAAAGGCGGACAGGTATCCGGTAAGCGGCAGGGTCGGAACAGGAGAGCGCACGAGGGAGCTTCCAGGGGGAAACGCCTGGTATCTTTATAGTCCTGTCGGGTTTCGCCACCTCTGACTTGAGCGTCGATTTTTGTGATGCTCGTCAGGGGGGCGGAGCCTATGGAAAAACGCCAGCAACGCGGCCTTTTTACGGTTCCTGGCCTTTTGCTGGCCTTTTGCTCACATGTTCTTTCCTGCGTTATCCCCTGATTCTGTGGATAACCGTATTACCGCCTTTGAGTGAGCTGATACCGCTCGCCGCAGCCGAACGACCGAGCGCAGCGAGTCAGTGAGCGAGGAAGCGGAAGAGCGCCTGATGCGGTATTTTCTCCTTACGCATCTGTGCGGTATTTCACACCGCATATATGGTGCACTCTCAGTACAATCTGCTCTGATGCCGCATAGTTAAGCCAGTATACACTCCGCTATCGCTACGTGACTGGGTCATGGCTGCGCCCCGACACCCGCCAACACCCGCTGACGCGCCCTGACGGGCTTGTCTGCTCCCGGCATCCGCTTACAGACAAGCTGTGACCGTCTCCGGGAGCTGCATGTGTCAGAGGTTTTCACCGTCATCACCGAAACGCGCGAGGCAGCTGCGGTAAAGCTCATCAGCGTGGTCGTGAAGCGATTCACAGATGTCTGCCTGTTCATCCGCGTCCAGCTCGTTGAGTTTCTCCAGAAGCGTTAATGTCTGGCTTCTGATAAAGCGGGCCATGTTAAGGGCGGTTTTTTCCTGTTTGGTCACTGATGCCTCCGTGTAAGGGGGATTTCTGTTCATGGGGGTAATGATACCGATGAAACGAGAGAGGATGCTCACGATACGGGTTACTGATGATGAACATGCCCGGTTACTGGAACGTTGTGAGGGTAAACAACTGGCGGTATGGATGCGGCGGGACCAGAGAAAAATCACTCAGGGTCAATGCCAGCGCTTCGTTAATACAGATGTAGGTGTTCCACAGGGTAGCCAGCAGCATCCTGCGATGCAGATCCGGAACATAATGGTGCAGGGCGCTGACTTCCGCGTTTCCAGACTTTACGAAACACGGAAACCGAAGACCATTCATGTTGTTGCTCAGGTCGCAGACGTTTTGCAGCAGCAGTCGCTTCACGTTCGCTCGCGTATCGGTGATTCATTCTGCTAACCAGTAAGGCAACCCCGCCAGCCTAGCCGGGTCCTCAACGACAGGAGCACGATCATGCGCACCCGTGGCCAGGACCCAACGCTGCCCGAGATGCGCCGCGTGCGGCTGCTGGAGATGGCGGACGCGATGGATATGTTCTGCCAAGGGTTGGTTTGCGCATTCACAGTTCTCCGCAAGAATTGATTGGCTCCAATTCTTGGAGTGGTGAATCCGTTAGCGAGGTGCCGCCGGCTTCCATTCAGGTCGAGGTGGCCCGGCTCCATGCACCGCGACGCAACGCGGGGAGGCAGACAAGGTATAGGGCGGCGCCTACAATCCATGCCAACCCGTTCCATGTGCTCGCCGAGGCGGCATAAATCGCCGTGACGATCAGCGGTCCAGTGATCGAAGTTAGGCTGGTAAGAGCCGCGAGCGATCCTTGAAGCTGTCCCTGATGGTCGTCATCTACCTGCCTGGACAGCATGGCCTGCAACGCGGGCATCCCGATGCCGCCGGAAGCGAGAAGAATCATAATGGGGAAGGCCATCCAGCCTCGCGTCGCGAACGCCAGCAAGACGTAGCCCAGCGCGTCGGCCGCCATGCCGGCGATAATGGCCTGCTTCTCGCCGAAACGTTTGGTGGCGGGACCAGTGACGAAGGCTTGAGCGAGGGCGTGCAAGATTCCGAATACCGCAAGCGACAGGCCGATCATCGTCGCGCTCCAGCGAAAGCGGTCCTCGCCGAAAATGACCCAGAGCGCTGCCGGCACCTGTCCTACGAGTTGCATGATAAAGAAGACAGTCATAAGTGCGGCGACGATAGTCATGCCCCGCGCCCACCGGAAGGAGCTGACTGGGTTGAAGGCTCTCAAGGGCATCGGTCGAGATCCCGGTGCCTAATGAGTGAGCTAACTTACATTAATTGCGTTGCGCTCACTGCCCGCTTTCCAGTCGGGAAACCTGTCGTGCCAGCTGCATTAATGAATCGGCCAACGCGCGGGGAGAGGCGGTTTGCGTATTGGGCGCCAGGGTGGTTTTTCTTTTCACCAGTGAGACGGGCAACAGCTGATTGCCCTTCACCGCCTGGCCCTGAGAGAGTTGCAGCAAGCGGTCCACGCTGGTTTGCCCCAGCAGGCGAAAATCCTGTTTGATGGTGGTTAACGGCGGGATATAACATGAGCTGTCTTCGGTATCGTCGTATCCCACTACCGAGATATCCGCACCAACGCGCAGCCCGGACTCGGTAATGGCGCGCATTGCGCCCAGCGCCATCTGATCGTTGGCAACCAGCATCGCAGTGGGAACGATGCCCTCATTCAGCATTTGCATGGTTTGTTGAAAACCGGACATGGCACTCCAGTCGCCTTCCCGTTCCGCTATCGGCTGAATTTGATTGCGAGTGAGATATTTATGCCAGCCAGCCAGACGCAGACGCGCCGAGACAGAACTTAATGGGCCCGCTAACAGCGCGATTTGCTGGTGACCCAATGCGACCAGATGCTCCACGCCCAGTCGCGTACCGTCTTCATGGGAGAAAATAATACTGTTGATGGGTGTCTGGTCAGAGACATCAAGAAATAACGCCGGAACATTAGTGCAGGCAGCTTCCACAGCAATGGCATCCTGGTCATCCAGCGGATAGTTAATGATCAGCCCACTGACGCGTTGCGCGAGAAGATTGTGCACCGCCGCTTTACAGGCTTCGACGCCGCTTCGTTCTACCATCGACACCACCACGCTGGCACCCAGTTGATCGGCGCGAGATTTAATCGCCGCGACAATTTGCGACGGCGCGTGCAGGGCCAGACTGGAGGTGGCAACGCCAATCAGCAACGACTGTTTGCCCGCCAGTTGTTGTGCCACGCGGTTGGGAATGTAATTCAGCTCCGCCATCGCCGCTTCCACTTTTTCCCGCGTTTTCGCAGAAACGTGGCTGGCCTGGTTCACCACGCGGGAAACGGTCTGATAAGAGACACCGGCATACTCTGCGACATCGTATAACGTTACTGGTTTCACATTCACCACCCTGAATTGACTCTCTTCCGGGCGCTATCATGCCATACCGCGAAAGGTTTTGCGCCATTCGATGGTGTCCGGGATCTCGACGCTCTCCCTTATGCGACTCCTGCATTAGGAAGCAGCCCAGTAGTAGGTTGAGGCCGTTGAGCACCGCCGCCGCAAGGAATGGTGCATGCAAGGAGATGGCGCCCAACAGTCCCCCGGCCACGGGGCCTGCCACCATACCCACGCCGAAACAAGCGCTCATGAGCCCGAAGTGGCGAGCCCGATCTTCCCCATCGGTGATGTCGGCGATATAGGCGCCAGCAACCGCACCTGTGGCGCCGGTGATGCCGGCCACGATGCGTCCGGCGTAGAGGATCGAGATCTCGATCCCGCGAAATTAATACGACTCACTATAGGGGAATTGTGAGCGGATAACAATTCCCCTCTAGAAATAATTTTGTTTAACTTTAAGAAGGAGAgttaggaggtctttatcatggaaaacaaaacatctttggggaataatatttattacaacccttttaaaccacaagataaatcgtattttgcgggttatttcaatgcagcaatggaaaatacagatagtgtatttagagaattgggaaaaagattaaaaggtaaagagtacacttcggagaatttttttgatgctatctttaaggaaaatatttcattagtggaatacgaaaggtatgtgaaattgctttctgattatttccctatggctcgtcttttggataaaaaagaagttcctataaaagaacgaaaagaaaattttaaaaagaattttaaaggcattataaaagccgtaagagatttaagaaacttttatacgcataaggaacacggagaggtagaaattacagatgaaatatttggcgtattggatgagatgctgaaaagcacggttttaacggtaaaaaagaaaaaagtaaaaaccgataaaacaaaagaaatcctgaaaaaaagcatagaaaaacaattggatattttatgccaaaagaaattagaatatctaagagacacagcaagaaaaatagaggagaagcgtagaaatcaaagagaaagaggagagaaagaattggtagctccattcaaatacagtgataaaagggatgatttaatagcagcgatttataatgatgcttttgatgtttatatcgataagaaaaaagatagcctaaaagaatccagcaaagcaaagtacaataccaaaagtgatcctcagcaagaagaaggcgatttgaaaattccaatttcgaaaaacggagtagtcttcctgctttctttgtttctcaccaaacaagaaatacacgcattcaaatctaaaatagcagggtttaaggctacggttatcgatgaagcaacggtatctgaagcaacggtgagccatggaaaaaacagcatttgttttatggctacgcacgaaattttctcgcatttggcttacaaaaaattgaaacgaaaagtgagaacggctgaaatcaactatggtgaggcagaaaatgctgaacagctttccgtttatgccaaagaaactctaatgatgcagatgcttgatgaattgagcaaagtacccgatgtggtctatcagaatttgagcgaagatgtacaaaaaactttcatcgaagattggaatgaatacctaaaagagaacaatggcgatgtagggacaatggaagaagagcaggtaattcatccagtgattcgcaagcgttacgaagataaattcaattattttgccattcgatttttagatgagtttgctcagttccctacgcttcgttttcaagtgcatttagggaattatcttcacgattctcgccctaaagaaaatctgatttctgaccgaagaataaaagaaaaaatcaccgttttcggaaggctttcggaattggaacataaaaaagcattgtttataaaaaataccgaaaccaatgaagatagagaacactattgggaaatttttcctaaccccaattatgattttcctaaggaaaatatttccgttaatgataaagattttcctatcgcgggtagtattttggacagagagaaacagcctgtcgcagggaaaattggtataaaagtgaaattgcttaaccaacaatatgtttcagaagtagataaagccgtaaaagcccatcagctcaagcaaagaaaagccagcaagccgtctatccagaacattatagaagaaatagtgccaatcaatgaaagcaatccaaaagaagctattgttttcggtgggcaacccacggcttatctgagtatgaatgacattcattccattttgtatgagtttttcgataaatgggagaaaaagaaagaaaagttagaaaagaaaggtgaaaaagaattaagaaaggaaattggaaaagagttagaaaagaaaatcgtaggaaaaattcaagctcaaatccagcaaatcatagataaagataccaatgccaagatattaaaaccttatcaagacggaaattctactgctattgataaagagaaattgataaaagatttgaagcaagagcaaaacattctacaaaaattaaaagatgaacaaacggttcgtgaaaaagaatataatgattttattgcttatcaagataaaaatagggagataaataaagtcagagatagaaatcataaacaatatttaaaagataaccttaaacgaaagtaccctgaagctcctgcacgaaaagaagttctttattatcgagaaaaaggaaaagtagcggtttggttggcgaatgacatcaaacgctttatgcctaccgatttcaaaaacgaatggaaaggggagcaacacagcttgttacaaaaatcgttggcgtattatgagcaatgtaaagaggagttgaaaaatcttttgcctgaaaaagtatttcagcatttaccttttaagttgggaggatattttcaacaaaaatatttgtaccaattttacacttgttatttagataaacggttggaatacatcagtgggttggtgcaacaagccgaaaactttaaatccgaaaataaagtcttcaaaaaggtagaaaatgagtgttttaaatttttgaaaaaacaaaattatactcacaaagaattagatgctcgggttcagtctatattggggtatcctatctttttagagcggggctttatggatgaaaaacctacgataatcaaaggaaaaacttttaaaggaaatgaagctctttttgccgattggtttaggtattacaaggaatatcagaattttcagacattttatgatactgaaaattatccgcttgtagaattggaaaaaaaacaagcagaccgaaaacgaaaaactaaaatctaccaacaaaagaaaaacgatgtgtttacacttttgatggcaaaacatattttcaaatcggtattcaagcaggatagcatcgatcagttttcgttagaagatttgtatcaatctcgggaggaacgcttaggaaatcaggagagagccagacaaacgggcgagcgaaatactaactatatttggaacaaaaccgttgatttaaagttgtgtgacggaaaaattacggtagaaaatgtaaagctaaaaaatgtaggagatttcataaaatatgaatacgaccaaagggttcaagctttcttaaaatatgaggaaaatatagaatggcaggcgtttctaatcaaagagagtaaggaagaggaaaattatccgtatgtggtggaaagagaaatagagcaatatgaaaaagtgagaagggaagaactgttgaaggaagtccaccttatagaggaatatattttggaaaaggtaaaagataaggaaatattaaaaaaaggggataatcaaaatttcaaatactacattctcaacggacttttaaagcaacttaaaaatgaagatgtggagagctataaggtattcaacttaaatactgaacctgaagatgtgaacattaaccaactaaaacaagaagcaaccgatttagaacaaaaagcttttgttttaacctatattcgtaataaatttgcacacaaccaattacctaaaaaagaattttgggactattgtcaagaaaaatacggcaaaatagaaaaagaaaaaacctatgccgagtattttgctgaggtatttaaaaaggaaaaagaagcactgataaagGGCGGTGGCTCGCATCATCATCATCATCACTAAGGATCCGGCTGCTAACAAAGCCCGAAAGGAAGCTGAGTTGGCTGCTGCCACCGCTGAGCAATAACTAGCATAACCCCTTGGGGCCTCTAAACGGGTCTTGAGGGGTTTTTTGCTGAAAGGAGGAACTATATCCGGATATCCCGCAAGAGGCCCGGCAGTACCGGCATAACCAAGCCTATGCCTACAGCATCCAGGGTGACGGTGCCGAGGATGACGATGAGCGCATTGTTAGATTTCATACACGGTGCCTGACTGCGTTAGCAATTTAACTGTGATAAACTACCGCATTAAAGCTTATCGATGATAAGCTGTCAAACATGAGAA

#### **pET15_RspCas13d-6xHIS**

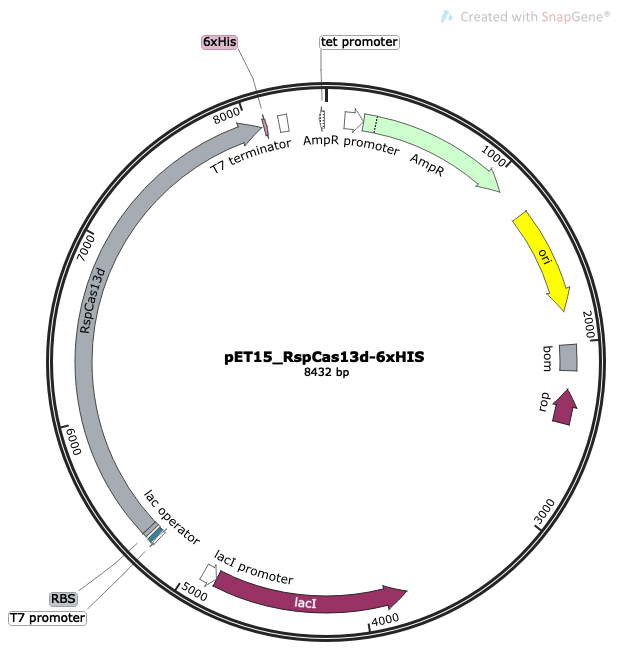

sequence:

TTCTTGAAGACGAAAGGGCCTCGTGATACGCCTATTTTTATAGGTTAATGTCATGATAATAATGGTTTCTTAGACGTCAGGTGGCACTTTTCGGGGAAATGTGCGCGGAACCCCTATTTGTTTATTTTTCTAAATACATTCAAATATGTATCCGCTCATGAGACAATAACCCTGATAAATGCTTCAATAATATTGAAAAAGGAAGAGTATGAGTATTCAACATTTCCGTGTCGCCCTTATTCCCTTTTTTGCGGCATTTTGCCTTCCTGTTTTTGCTCACCCAGAAACGCTGGTGAAAGTAAAAGATGCTGAAGATCAGTTGGGTGCACGAGTGGGTTACATCGAACTGGATCTCAACAGCGGTAAGATCCTTGAGAGTTTTCGCCCCGAAGAACGTTTTCCAATGATGAGCACTTTTAAAGTTCTGCTATGTGGCGCGGTATTATCCCGTGTTGACGCCGGGCAAGAGCAACTCGGTCGCCGCATACACTATTCTCAGAATGACTTGGTTGAGTACTCACCAGTCACAGAAAAGCATCTTACGGATGGCATGACAGTAAGAGAATTATGCAGTGCTGCCATAACCATGAGTGATAACACTGCGGCCAACTTACTTCTGACAACGATCGGAGGACCGAAGGAGCTAACCGCTTTTTTGCACAACATGGGGGATCATGTAACTCGCCTTGATCGTTGGGAACCGGAGCTGAATGAAGCCATACCAAACGACGAGCGTGACACCACGATGCCTGCAGCAATGGCAACAACGTTGCGCAAACTATTAACTGGCGAACTACTTACTCTAGCTTCCCGGCAACAATTAATAGACTGGATGGAGGCGGATAAAGTTGCAGGACCACTTCTGCGCTCGGCCCTTCCGGCTGGCTGGTTTATTGCTGATAAATCTGGAGCCGGTGAGCGTGGGTCTCGCGGTATCATTGCAGCACTGGGGCCAGATGGTAAGCCCTCCCGTATCGTAGTTATCTACACGACGGGGAGTCAGGCAACTATGGATGAACGAAATAGACAGATCGCTGAGATAGGTGCCTCACTGATTAAGCATTGGTAACTGTCAGACCAAGTTTACTCATATATACTTTAGATTGATTTAAAACTTCATTTTTAATTTAAAAGGATCTAGGTGAAGATCCTTTTTGATAATCTCATGACCAAAATCCCTTAACGTGAGTTTTCGTTCCACTGAGCGTCAGACCCCGTAGAAAAGATCAAAGGATCTTCTTGAGATCCTTTTTTTCTGCGCGTAATCTGCTGCTTGCAAACAAAAAAACCACCGCTACCAGCGGTGGTTTGTTTGCCGGATCAAGAGCTACCAACTCTTTTTCCGAAGGTAACTGGCTTCAGCAGAGCGCAGATACCAAATACTGTCCTTCTAGTGTAGCCGTAGTTAGGCCACCACTTCAAGAACTCTGTAGCACCGCCTACATACCTCGCTCTGCTAATCCTGTTACCAGTGGCTGCTGCCAGTGGCGATAAGTCGTGTCTTACCGGGTTGGACTCAAGACGATAGTTACCGGATAAGGCGCAGCGGTCGGGCTGAACGGGGGGTTCGTGCACACAGCCCAGCTTGGAGCGAACGACCTACACCGAACTGAGATACCTACAGCGTGAGCTATGAGAAAGCGCCACGCTTCCCGAAGGGAGAAAGGCGGACAGGTATCCGGTAAGCGGCAGGGTCGGAACAGGAGAGCGCACGAGGGAGCTTCCAGGGGGAAACGCCTGGTATCTTTATAGTCCTGTCGGGTTTCGCCACCTCTGACTTGAGCGTCGATTTTTGTGATGCTCGTCAGGGGGGCGGAGCCTATGGAAAAACGCCAGCAACGCGGCCTTTTTACGGTTCCTGGCCTTTTGCTGGCCTTTTGCTCACATGTTCTTTCCTGCGTTATCCCCTGATTCTGTGGATAACCGTATTACCGCCTTTGAGTGAGCTGATACCGCTCGCCGCAGCCGAACGACCGAGCGCAGCGAGTCAGTGAGCGAGGAAGCGGAAGAGCGCCTGATGCGGTATTTTCTCCTTACGCATCTGTGCGGTATTTCACACCGCATATATGGTGCACTCTCAGTACAATCTGCTCTGATGCCGCATAGTTAAGCCAGTATACACTCCGCTATCGCTACGTGACTGGGTCATGGCTGCGCCCCGACACCCGCCAACACCCGCTGACGCGCCCTGACGGGCTTGTCTGCTCCCGGCATCCGCTTACAGACAAGCTGTGACCGTCTCCGGGAGCTGCATGTGTCAGAGGTTTTCACCGTCATCACCGAAACGCGCGAGGCAGCTGCGGTAAAGCTCATCAGCGTGGTCGTGAAGCGATTCACAGATGTCTGCCTGTTCATCCGCGTCCAGCTCGTTGAGTTTCTCCAGAAGCGTTAATGTCTGGCTTCTGATAAAGCGGGCCATGTTAAGGGCGGTTTTTTCCTGTTTGGTCACTGATGCCTCCGTGTAAGGGGGATTTCTGTTCATGGGGGTAATGATACCGATGAAACGAGAGAGGATGCTCACGATACGGGTTACTGATGATGAACATGCCCGGTTACTGGAACGTTGTGAGGGTAAACAACTGGCGGTATGGATGCGGCGGGACCAGAGAAAAATCACTCAGGGTCAATGCCAGCGCTTCGTTAATACAGATGTAGGTGTTCCACAGGGTAGCCAGCAGCATCCTGCGATGCAGATCCGGAACATAATGGTGCAGGGCGCTGACTTCCGCGTTTCCAGACTTTACGAAACACGGAAACCGAAGACCATTCATGTTGTTGCTCAGGTCGCAGACGTTTTGCAGCAGCAGTCGCTTCACGTTCGCTCGCGTATCGGTGATTCATTCTGCTAACCAGTAAGGCAACCCCGCCAGCCTAGCCGGGTCCTCAACGACAGGAGCACGATCATGCGCACCCGTGGCCAGGACCCAACGCTGCCCGAGATGCGCCGCGTGCGGCTGCTGGAGATGGCGGACGCGATGGATATGTTCTGCCAAGGGTTGGTTTGCGCATTCACAGTTCTCCGCAAGAATTGATTGGCTCCAATTCTTGGAGTGGTGAATCCGTTAGCGAGGTGCCGCCGGCTTCCATTCAGGTCGAGGTGGCCCGGCTCCATGCACCGCGACGCAACGCGGGGAGGCAGACAAGGTATAGGGCGGCGCCTACAATCCATGCCAACCCGTTCCATGTGCTCGCCGAGGCGGCATAAATCGCCGTGACGATCAGCGGTCCAGTGATCGAAGTTAGGCTGGTAAGAGCCGCGAGCGATCCTTGAAGCTGTCCCTGATGGTCGTCATCTACCTGCCTGGACAGCATGGCCTGCAACGCGGGCATCCCGATGCCGCCGGAAGCGAGAAGAATCATAATGGGGAAGGCCATCCAGCCTCGCGTCGCGAACGCCAGCAAGACGTAGCCCAGCGCGTCGGCCGCCATGCCGGCGATAATGGCCTGCTTCTCGCCGAAACGTTTGGTGGCGGGACCAGTGACGAAGGCTTGAGCGAGGGCGTGCAAGATTCCGAATACCGCAAGCGACAGGCCGATCATCGTCGCGCTCCAGCGAAAGCGGTCCTCGCCGAAAATGACCCAGAGCGCTGCCGGCACCTGTCCTACGAGTTGCATGATAAAGAAGACAGTCATAAGTGCGGCGACGATAGTCATGCCCCGCGCCCACCGGAAGGAGCTGACTGGGTTGAAGGCTCTCAAGGGCATCGGTCGAGATCCCGGTGCCTAATGAGTGAGCTAACTTACATTAATTGCGTTGCGCTCACTGCCCGCTTTCCAGTCGGGAAACCTGTCGTGCCAGCTGCATTAATGAATCGGCCAACGCGCGGGGAGAGGCGGTTTGCGTATTGGGCGCCAGGGTGGTTTTTCTTTTCACCAGTGAGACGGGCAACAGCTGATTGCCCTTCACCGCCTGGCCCTGAGAGAGTTGCAGCAAGCGGTCCACGCTGGTTTGCCCCAGCAGGCGAAAATCCTGTTTGATGGTGGTTAACGGCGGGATATAACATGAGCTGTCTTCGGTATCGTCGTATCCCACTACCGAGATATCCGCACCAACGCGCAGCCCGGACTCGGTAATGGCGCGCATTGCGCCCAGCGCCATCTGATCGTTGGCAACCAGCATCGCAGTGGGAACGATGCCCTCATTCAGCATTTGCATGGTTTGTTGAAAACCGGACATGGCACTCCAGTCGCCTTCCCGTTCCGCTATCGGCTGAATTTGATTGCGAGTGAGATATTTATGCCAGCCAGCCAGACGCAGACGCGCCGAGACAGAACTTAATGGGCCCGCTAACAGCGCGATTTGCTGGTGACCCAATGCGACCAGATGCTCCACGCCCAGTCGCGTACCGTCTTCATGGGAGAAAATAATACTGTTGATGGGTGTCTGGTCAGAGACATCAAGAAATAACGCCGGAACATTAGTGCAGGCAGCTTCCACAGCAATGGCATCCTGGTCATCCAGCGGATAGTTAATGATCAGCCCACTGACGCGTTGCGCGAGAAGATTGTGCACCGCCGCTTTACAGGCTTCGACGCCGCTTCGTTCTACCATCGACACCACCACGCTGGCACCCAGTTGATCGGCGCGAGATTTAATCGCCGCGACAATTTGCGACGGCGCGTGCAGGGCCAGACTGGAGGTGGCAACGCCAATCAGCAACGACTGTTTGCCCGCCAGTTGTTGTGCCACGCGGTTGGGAATGTAATTCAGCTCCGCCATCGCCGCTTCCACTTTTTCCCGCGTTTTCGCAGAAACGTGGCTGGCCTGGTTCACCACGCGGGAAACGGTCTGATAAGAGACACCGGCATACTCTGCGACATCGTATAACGTTACTGGTTTCACATTCACCACCCTGAATTGACTCTCTTCCGGGCGCTATCATGCCATACCGCGAAAGGTTTTGCGCCATTCGATGGTGTCCGGGATCTCGACGCTCTCCCTTATGCGACTCCTGCATTAGGAAGCAGCCCAGTAGTAGGTTGAGGCCGTTGAGCACCGCCGCCGCAAGGAATGGTGCATGCAAGGAGATGGCGCCCAACAGTCCCCCGGCCACGGGGCCTGCCACCATACCCACGCCGAAACAAGCGCTCATGAGCCCGAAGTGGCGAGCCCGATCTTCCCCATCGGTGATGTCGGCGATATAGGCGCCAGCAACCGCACCTGTGGCGCCGGTGATGCCGGCCACGATGCGTCCGGCGTAGAGGATCGAGATCTCGATCCCGCGAAATTAATACGACTCACTATAGGGGAATTGTGAGCGGATAACAATTCCCCTCTAGAAATAATTTTGTTTAACTTTAAGAAGGAGATATACCATGgccaaaaagaacaaaatgaaaccgcgtgaactgcgtgaagcacagaaaaaagcacgtcagctgaaagcagcagaaattaacaataatgcagcaccggcaattgcagcaatgcctgcagcagaagttattgcaccggttgcagaaaaaaagaaaagcagcgttaaagcagccggtatgaaaagcattctggtgagcaaaaacaaaatgtatatcaccagctttggcaaaggtaatagcgcagttctggaatatgaagtggataacaacgattacaatcagacccagctgagcagcaaaggtagcagcaatattgagctgcgtggtgttaatgaagtgaacattacctttagcagcaaacacggttttgaaagcggtgttgaaatcaataccagcaatccgacacatcgtagcggtgaaagcagtccggttcgtggtgatatgctgggcctgaaaagcgaactggaaaaacgcttttttggcaaaaccttcgatgacaacattcatatccagctgatctataacatcctggacatcgaaaaaattctggccgtttatgtgaccaacattgtgtatgcactgaataacatgctgagcattaaagatagcgagagctatgatgatttcatgggttatctgagcgcacgcaatacctatgaagtttttacccatccggataaaagcaacctgagcgataaagcaaaaggcaacatcaaaaaatcctttagcaccttcaacgacctgctgaaaaccaaacgtctgggttattttggtctggaagaaccgaaaacaaaagatacccgtgttagccaggcatacaaaaaacgtgtttatcacatgctggcaattgtgggtcagattcgtcagagcgtgtttcatgataaaagtagcaaactggatgaggacctgtatagctttatcgatatcatcgattccgaatatcgtgaaaccctggattatctggttgatgaacgttttgacagcatcaacaaaggttttatccagggcaacaaagtgaatattagcctgctgatcgatatgatgaaaggctatgaagccgatgatattatccgcctgtattatgattttatcgtgctgaaaagccagaagaatctgggctttagtatcaaaaaactgcgcgagaaaatgctggatgaatatggctttcgcttcaaagataaacagtatgatagcgtgcgtagcaagatgtataaactgatggattttctgctgttctgcaactattatcgcaatgatgttgttgccggtgaagcactggttcgtaaactgcgttttagcatgaccgatgatgaaaaagaaggtatttacgcagacgaagcaagcaaactgtggggtaaatttcgtaacgattttgagaacattgccgaccatatgaacggtgatgttattaaagaactgggcaaagccgatatggatttcgatgaaaaaatactggacagcgagaaaaaaaacgcaagcgatctgctgtatttcagcaagatgatttatatgctgacctatttcctggatggcaaagaaattaatgatctgctgaccacgctgattagcaaatttgataacatcaaagaatttctgaaaatcatgaaaagctcagccgttgatgttgaatgtgaactgaccgcaggttataaactgtttaatgatagccagcgcattaccaacgaactgttcattgttaaaaacattgcgagcatgcgtaaaccggcaagcagcgcaaaactgaccatgtttcgtgatgcactgaccattctgggtattgatgataacattaccgatgatcgcattagcgaaatcctgaagctgaaagaaaaaggtaaaggtattcatggcctgcgcaactttattaccaataacgtgattgaaagctcccgctttgtgtacctgatcaaatatgcaaatgcccagaaaattcgcaaagtggcgaaaaatgaaaaggtggtgatgtttgtgttaggtggtattccggatacacagattgaacgctattacaaaagctgcgttgaatttccggatatgaacagcagcctggaagttaaacgtagtgaactggcacgtatgatcaaaaacatcagcttcgacgatttcaagaacgttaaacagcaggcaaaaggtcgtgaaaatgttgcaaaagaacgtgccaaagcagtgattggtctgtatctgaccgttatgtatctgctggttaaaaatctggttaacgtgaatgcccgttatgtgattgcaattcattgtctggaacgtgatttcggcctgtataaagaaatcattccggaactggcaagcaagaacctgaaaaatgattatcgtattctgagtcagaccctgtgtgaactgtgtgataaaagcccgaacctgtttctgaagaaaaacgaacgtctgcgcaaatgcgttgaggtggatattaacaatgcagatagcagcatgacccgcaaatatcgtaattgtattgcacatctgacagttgtgcgcgaactgaaagaatatattggtgatattcgtaccgtggacagctactttagcatctatcattatgttatgcagcggtgtattaccaaacgcgaaaatgataccaaacaagaagagaaaatcaaatacgaggacgatctgcttaaaaaccacggctataccaaagattttgtgaaagccctgaatagcccgttcggttataacattccgcgtttcaaaaatctgagcatcgagcagctgtttgatcgtaatgaatatctgacggaaaaaGGCGGTGGCTCGCATCATCATCATCATCACTAATAAGATCCGGCTGCTAACAAAGCCCGAAAGGAAGCTGAGTTGGCTGCTGCCACCGCTGAGCAATAACTAGCATAACCCCTTGGGGCCTCTAAACGGGTCTTGAGGGGTTTTTTGCTGAAAGGAGGAACTATATCCGGATATCCCGCAAGAGGCCCGGCAGTACCGGCATAACCAAGCCTATGCCTACAGCATCCAGGGTGACGGTGCCGAGGATGACGATGAGCGCATTGTTAGATTTCATACACGGTGCCTGACTGCGTTAGCAATTTAACTGTGATAAACTACCGCATTAAAGCTTATCGATGATAAGCTGTCAAACATGAGAA

#### pET-28b-RfxCas13d-His

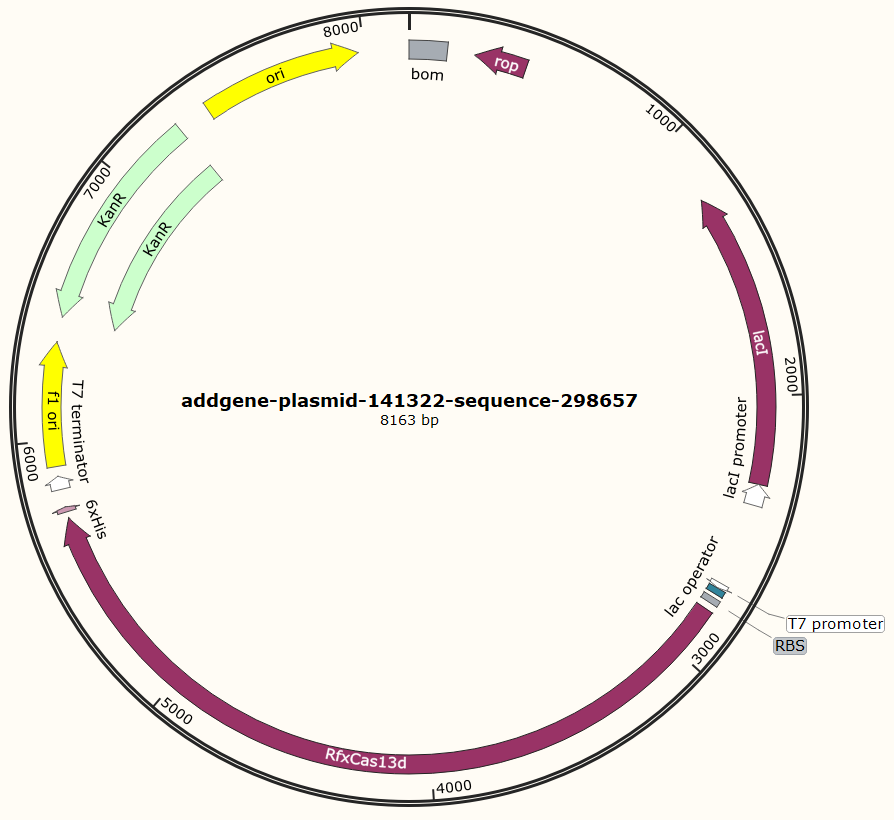

cctgatgcggtattttctccttacgcatctgtgcggtatttcacaccgcaatggtgcactctcagtacaatctgctctgatgccgcatagttaagccagtatacactccgctatcgctacgtgactgggtcatggctgcgccccgacacccgccaacacccgctgacgcgccctgacgggcttgtctgctcccggcatccgcttacagacaagctgtgaccgtctccgggagctgcatgtgtcagaggttttcaccgtcatcaccgaaacgcgcgaggcagctgcggtaaagctcatcagcgtggtcgtgaagcgattcacagatgtctgcctgttcatccgcgtccagctcgttgagtttctccagaagcgttaatgtctggcttctgataaagcgggccatgttaagggcggttttttcctgtttggtcactgatgcctccgtgtaagggggatttctgttcatgggggtaatgataccgatgaaacgagagaggatgctcacgatacgggttactgatgatgaacatgcccggttactggaacgttgtgagggtaaacaactggcggtatggatgcggcgggaccagagaaaaatcactcagggtcaatgccagcgcttcgttaatacagatgtaggtgttccacagggtagccagcagcatcctgcgatgcagatccggaacataatggtgcagggcgctgacttccgcgtttccagactttacgaaacacggaaaccgaagaccattcatgttgttgctcaggtcgcagacgttttgcagcagcagtcgcttcacgttcgctcgcgtatcggtgattcattctgctaaccagtaaggcaaccccgccagcctagccgggtcctcaacgacaggagcacgatcatgcgcacccgtggggccgccatgccggcgataatggcctgcttctcgccgaaacgtttggtggcgggaccagtgacgaaggcttgagcgagggcgtgcaagattccgaataccgcaagcgacaggccgatcatcgtcgcgctccagcgaaagcggtcctcgccgaaaatgacccagagcgctgccggcacctgtcctacgagttgcatgataaagaagacagtcataagtgcggcgacgatagtcatgccccgcgcccaccggaaggagctgactgggttgaaggctctcaagggcatcggtcgagatcccggtgcctaatgagtgagctaacttacattaattgcgttgcgctcactgcccgctttccagtcgggaaacctgtcgtgccagctgcattaatgaatcggccaacgcgcggggagaggcggtttgcgtattgggcgccagggtggtttttcttttcaccagtgagacgggcaacagctgattgcccttcaccgcctggccctgagagagttgcagcaagcggtccacgctggtttgccccagcaggcgaaaatcctgtttgatggtggttaacggcgggatataacatgagctgtcttcggtatcgtcgtatcccactaccgagatatccgcaccaacgcgcagcccggactcggtaatggcgcgcattgcgcccagcgccatctgatcgttggcaaccagcatcgcagtgggaacgatgccctcattcagcatttgcatggtttgttgaaaaccggacatggcactccagtcgccttcccgttccgctatcggctgaatttgattgcgagtgagatatttatgccagccagccagacgcagacgcgccgagacagaacttaatgggcccgctaacagcgcgatttgctggtgacccaatgcgaccagatgctccacgcccagtcgcgtaccgtcttcatgggagaaaataatactgttgatgggtgtctggtcagagacatcaagaaataacgccggaacattagtgcaggcagcttccacagcaatggcatcctggtcatccagcggatagttaatgatcagcccactgacgcgttgcgcgagaagattgtgcaccgccgctttacaggcttcgacgccgcttcgttctaccatcgacaccaccacgctggcacccagttgatcggcgcgagatttaatcgccgcgacaatttgcgacggcgcgtgcagggccagactggaggtggcaacgccaatcagcaacgactgtttgcccgccagttgttgtgccacgcggttgggaatgtaattcagctccgccatcgccgcttccactttttcccgcgttttcgcagaaacgtggctggcctggttcaccacgcgggaaacggtctgataagagacaccggcatactctgcgacatcgtataacgttactggtttcacattcaccaccctgaattgactctcttccgggcgctatcatgccataccgcgaaaggttttgcgccattcgatggtgtccgggatctcgacgctctcccttatgcgactcctgcattaggaagcagcccagtagtaggttgaggccgttgagcaccgccgccgcaaggaatggtgcatgcaaggagatggcgcccaacagtcccccggccacggggcctgccaccatacccacgccgaaacaagcgctcatgagcccgaagtggcgagcccgatcttccccatcggtgatgtcggcgatataggcgccagcaaccgcacctgtggcgccggtgatgccggccacgatgcgtccggcgtagaggatcgagatctcgatcccgcgaaattaatacgactcactataggggaattgtgagcggataacaattcccctctagaaataattttgtttaactttaagaaggagatataccatggcgagcgaggccagcatcgaaaaaaaaaagtccttcgccaagggcatgggcgtgaagtccacactcgtgtccggctccaaagtgtacatgacaaccttcgccgaaggcagcgacgccaggctggaaaagatcgtggagggcgacagcatcaggagcgtgaatgagggcgaggccttcagcgctgaaatggccgataaaaacgccggctataagatcggcaacgccaaattcagccatcctaagggctacgccgtggtggctaacaaccctctgtatacaggacccgtccagcaggatatgctcggcctgaaggaaactctggaaaagaggtacttcggcgagagcgctgatggcaatgacaatatttgtatccaggtgatccataacatcctggacattgaaaaaatcctcgccgaatacattaccaacgccgcctacgccgtcaacaatatctccggcctggataaggacattattggattcggcaagttctccacagtgtatacctacgacgaattcaaagaccccgagcaccatagggccgctttcaacaataacgataagctcatcaacgccatcaaggcccagtatgacgagttcgacaacttcctcgataaccccagactcggctatttcggccaggcctttttcagcaaggagggcagaaattacatcatcaattacggcaacgaatgctatgacattctggccctcctgagcggactgaggcactgggtggtccataacaacgaagaagagtccaggatctccaggacctggctctacaacctcgataagaacctcgacaacgaatacatctccaccctcaactacctctacgacaggatcaccaatgagctgaccaactccttctccaagaactccgccgccaacgtgaactatattgccgaaactctgggaatcaaccctgccgaattcgccgaacaatatttcagattcagcattatgaaagagcagaaaaacctcggattcaatatcaccaagctcagggaagtgatgctggacaggaaggatatgtccgagatcaggaaaaatcataaggtgttcgactccatcaggaccaaggtctacaccatgatggactttgtgatttataggtattacatcgaagaggatgccaaggtggctgccgccaataagtccctccccgataatgagaagtccctgagcgagaaggatatctttgtgattaacctgaggggctccttcaacgacgaccagaaggatgccctctactacgatgaagctaatagaatttggagaaagctcgaaaatatcatgcacaacatcaaggaatttaggggaaacaagacaagagagtataagaagaaggacgcccctagactgcccagaatcctgcccgctggccgtgatgtttccgccttcagcaaactcatgtatgccctgaccatgttcctggatggcaaggagatcaacgacctcctgaccaccctgattaataaattcgataacatccagagcttcctgaaggtgatgcctctcatcggagtcaacgctaagttcgtggaggaatacgcctttttcaaagactccgccaagatcgccgatgagctgaggctgatcaagtccttcgctagaatgggagaacctattgccgatgccaggagggccatgtatatcgacgccatccgtattttaggaaccaacctgtcctatgatgagctcaaggccctcgccgacaccttttccctggacgagaacggaaacaagctcaagaaaggcaagcacggcatgagaaatttcattattaataacgtgatcagcaataaaaggttccactacctgatcagatacggtgatcctgcccacctccatgagatcgccaaaaacgaggccgtggtgaagttcgtgctcggcaggatcgctgacatccagaaaaaacagggccagaacggcaagaaccagatcgacaggtactacgaaacttgtatcggaaaggataagggcaagagcgtgagcgaaaaggtggacgctctcacaaagatcatcaccggaatgaactacgaccaattcgacaagaaaaggagcgtcattgaggacaccggcagggaaaacgccgagagggagaagtttaaaaagatcatcagcctgtacctcaccgtgatctaccacatcctcaagaatattgtcaatatcaacgccaggtacgtcatcggattccattgcgtcgagcgtgatgctcaactgtacaaggagaaaggctacgacatcaatctcaagaaactggaagagaagggattcagctccgtcaccaagctctgcgctggcattgatgaaactgcccccgataagagaaaggacgtggaaaaggagatggctgaaagagccaaggagagcattgacagcctcgagagcgccaaccccaagctgtatgccaattacatcaaatacagcgacgagaagaaagccgaggagttcaccaggcagattaacagggagaaggccaaaaccgccctgaacgcctacctgaggaacaccaagtggaatgtgatcatcagggaggacctcctgagaattgacaacaagacatgtaccctgttcagaaacaaggccgtccacctggaagtggccaggtatgtccacgcctatatcaacgacattgccgaggtcaattcctacttccaactgtaccattacatcatgcagagaattatcatgaatgagaggtacgagaaaagcagcggaaaggtgtccgagtacttcgacgctgtgaatgacgagaagaagtacaacgataggctcctgaaactgctgtgtgtgcctttcggctactgtatccccaggtttaagaacctgagcatcgaggccctgttcgataggaacgaggccgccaagttcgacaaggagaaaaagaaggtgtccggcaattccggatccggagcggccgcactcgagcaccaccaccaccaccactgagatccggctgctaacaaagcccgaaaggaagctgagttggctgctgccaccgctgagcaataactagcataaccccttggggcctctaaacgggtcttgaggggttttttgctgaaaggaggaactatatccggattggcgaatgggacgcgccctgtagcggcgcattaagcgcggcgggtgtggtggttacgcgcagcgtgaccgctacacttgccagcgccctagcgcccgctcctttcgctttcttcccttcctttctcgccacgttcgccggctttccccgtcaagctctaaatcgggggctccctttagggttccgatttagtgctttacggcacctcgaccccaaaaaacttgattagggtgatggttcacgtagtgggccatcgccctgatagacggtttttcgccctttgacgttggagtccacgttctttaatagtggactcttgttccaaactggaacaacactcaaccctatctcggtctattcttttgatttataagggattttgccgatttcggcctattggttaaaaaatgagctgatttaacaaaaatttaacgcgaattttaacaaaatattaacgcttacaatttaggtggcacttttcggggaaatgtgcgcggaacccctatttgtttatttttctaaatacattcaaatatgtatccgctcatgaattaattcttagaaaaactcatcgagcatcaaatgaaactgcaatttattcatatcaggattatcaataccatatttttgaaaaagccgtttctgtaatgaaggagaaaactcaccgaggcagttccataggatggcaagatcctggtatcggtctgcgattccgactcgtccaacatcaatacaacctattaatttcccctcgtcaaaaataaggttatcaagtgagaaatcaccatgagtgacgactgaatccggtgagaatggcaaaagtttatgcatttctttccagacttgttcaacaggccagccattacgctcgtcatcaaaatcactcgcatcaaccaaaccgttattcattcgtgattgcgcctgagcgagacgaaatacgcgatcgctgttaaaaggacaattacaaacaggaatcgaatgcaaccggcgcaggaacactgccagcgcatcaacaatattttcacctgaatcaggatattcttctaatacctggaatgctgttttcccggggatcgcagtggtgagtaaccatgcatcatcaggagtacggataaaatgcttgatggtcggaagaggcataaattccgtcagccagtttagtctgaccatctcatctgtaacatcattggcaacgctacctttgccatgtttcagaaacaactctggcgcatcgggcttcccatacaatcgatagattgtcgcacctgattgcccgacattatcgcgagcccatttatacccatataaatcagcatccatgttggaatttaatcgcggcctagagcaagacgtttcccgttgaatatggctcataacaccccttgtattactgtttatgtaagcagacagttttattgttcatgaccaaaatcccttaacgtgagttttcgttccactgagcgtcagaccccgtagaaaagatcaaaggatcttcttgagatcctttttttctgcgcgtaatctgctgcttgcaaacaaaaaaaccaccgctaccagcggtggtttgtttgccggatcaagagctaccaactctttttccgaaggtaactggcttcagcagagcgcagataccaaatactgtccttctagtgtagccgtagttaggccaccacttcaagaactctgtagcaccgcctacatacctcgctctgctaatcctgttaccagtggctgctgccagtggcgataagtcgtgtcttaccgggttggactcaagacgatagttaccggataaggcgcagcggtcgggctgaacggggggttcgtgcacacagcccagcttggagcgaacgacctacaccgaactgagatacctacagcgtgagctatgagaaagcgccacgcttcccgaagggagaaaggcggacaggtatccggtaagcggcagggtcggaacaggagagcgcacgagggagcttccagggggaaacgcctggtatctttatagtcctgtcgggtttcgccacctctgacttgagcgtcgatttttgtgatgctcgtcaggggggcggagcctatggaaaaacgccagcaacgcggcctttttacggttcctggccttttgctggccttttgctcacatgttctttcctgcgttatcccctgattctgtggataaccgtattaccgcctttgagtgagctgataccgctcgccgcagccgaacgaccgagcgcagcgagtcagtgagcgaggaagcggaagagcg

#### pLentiviral_EF-1a-LbuCas13a-IRES-TagBFP-P2A-BSD

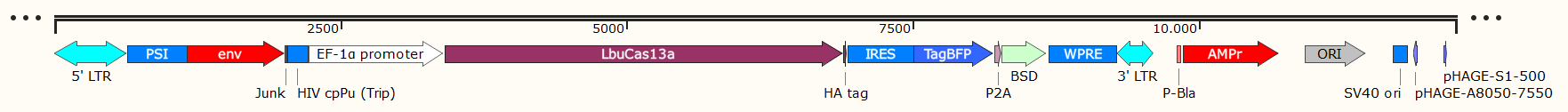

tggaagggctaattcactcccaaagaagacaagatatccttgatctgtggatctaccacacacaaggctacttccctgattagcagaactacacaccagggccaggggtcagatatccactgacctttggatggtgctacaagctagtaccagttgagccagataaggtagaagaggccaataaaggagagaacaccagcttgttacaccctgtgagcctgcatgggatggatgacccggagagagaagtgttagagtggaggtttgacagccgcctagcatttcatcacgtggcccgagagctgcatccggagtacttcaagaactgctgatatcgagcttgctacaagggactttccgctggggactttccagggaggcgtggcctgggcgggactggggagtggcgagccctcagatcctgcatataagcagctgctttttgcctgtactgggtctctctggttagaccagatctgagcctgggagctctctggctaactagggaacccactgcttaagcctcaataaagcttgccttgagtgcttcaagtagtgtgtgcccgtctgttgtgtgactctggtaactagagatccctcagacccttttagtcagtgtggaaaatctctagcagtggcgcccgaacagggacttgaaagcgaaagggaaaccagaggagctctctcgacgcaggactcggcttgctgaagcgcgcacggcaagaggcgaggggcggcgactggtgagtacgccaaaaattttgactagcggaggctagaaggagagagatgggtgcgagagcgtcagtattaagcgggggagaattagatcgcgatgggaaaaaattcggttaaggccagggggaaagaaaaaatataaattaaaacatatagtatgggcaagcagggagctagaacgattcgcagttaatcctggcctgttagaaacatcagaaggctgtagacaaatactgggacagctacaaccatcccttcagacaggatcagaagaacttagatcattatataatacagtagcaaccctctattgtgtgcatcaaaggatagagataaaagacaccaaggaagctttagacaagatagaggaagagcaaaacaaaagtaagaccaccgcacagcaagcggccggccgctgatcttcagacctggaggaggagatatgagggacaattggagaagtgaattatataaatataaagtagtaaaaattgaaccattaggagtagcacccaccaaggcaaagagaagagtggtgcagagagaaaaaagagcagtgggaataggagctttgttccttgggttcttgggagcagcaggaagcactatgggcgcagcgtcaatgacgctgacggtacaggccagacaattattgtctggtatagtgcagcagcagaacaatttgctgagggctattgaggcgcaacagcatctgttgcaactcacagtctggggcatcaagcagctccaggcaagaatcctggctgtggaaagatacctaaaggatcaacagctcctggggatttggggttgctctggaaaactcatttgcaccactgctgtgccttggaatgctagttggagtaataaatctctggaacagatttggaatcacacgacctggatggagtgggacagagaaattaacaattacacaagcttaatacactccttaattgaagaatcgcaaaaccagcaagaaaagaatgaacaagaattattggaattagataaatgggcaagtttgtggaattggtttaacataacaaattggctgtggtatataaaattattcataatgatagtaggaggcttggtaggtttaagaatagtttttgctgtactttctatagtgaatagagttaggcagggatattcaccattatcgtttcagacccacctcccaaccccgaggggacccgacaggcccgaaggaatagaagaagaaggtggagagagagacagagacagatccattcgattagtgaacggatctcgacggtatcgccgaattcacaaatggcagtattcatccacaattttaaaagaaaaggggggattggggggtacagtgcaggggaaagaatagtagacataatagcaacagacatacaaactaaagaattacaaaaacaaattacaaaaattcaaaattttcgggtttattacagggacagcagagatccagtttggactagtcgtgaggctccggtgcccgtcagtgggcagagcgcacatcgcccacagtccccgagaagttggggggaggggtcggcaattgaaccggtgcctagagaaggtggcgcggggtaaactgggaaagtgatgtcgtgtactggctccgcctttttcccgagggtgggggagaaccgtatataagtgcagtagtcgccgtgaacgttctttttcgcaacgggtttgccgccagaacacaggtaagtgccgtgtgtggttcccgcgggcctggcctctttacgggttatggcccttgcgtgccttgaattacttccacctggctgcagtacgtgattcttgatcccgagcttcgggttggaagtgggtgggagagttcgaggccttgcgcttaaggagccccttcgcctcgtgcttgagttgaggcctggcctgggcgctggggccgccgcgtgcgaatctggtggcaccttcgcgcctgtctcgctgctttcgataagtctctagccatttaaaatttttgatgacctgctgcgacgctttttttctggcaagatagtcttgtaaatgcgggccaagatctgcacactggtatttcggtttttggggccgcgggcggcgacggggcccgtgcgtcccagcgcacatgttcggcgaggcggggcctgcgagcgcggccaccgagaatcggacgggggtagtctcaagctggccggcctgctctggtgcctggcctcgcgccgccgtgtatcgccccgccctgggcggcaaggctggcccggtcggcaccagttgcgtgagcggaaagatggccgcttcccggccctgctgcagggagctcaaaatggaggacgcggcgctcgggagagcgggcgggtgagtcacccacacaaaggaaaagggcctttccgtcctcagccgtcgcttcatgtgactccacggagtaccgggcgccgtccaggcacctcgattagttctcgagcttttggagtacgtcgtctttaggttggggggaggggttttatgcgatggagtttccccacactgagtgggtggagactgaagttaggccagcttggcacttgatgtaattctccttggaatttgccctttttgagtttggatcttggttcattctcaagcctcagacagtggttcaaagtttttttcttccatttcaggtgtcgtgaagcggccgccaccATGAAGGTAACCAAAGTAGGAGGTATCTCACATAAGAAGTATACCAGTGAAGGGAGACTTGTAAAGAGTGAATCAGAGGAGAATCGCACAGACGAACGCTTGTCTGCTCTGCTCAACATGCGCCTGGATATGTATATAAAGAACCCTTCCTCCACGGAGACAAAGGAGAACCAGAAGAGGATTGGGAAGTTGAAGAAGTTCTTCTCCAATAAAATGGTGTATCTGAAAGACAATACCCTTTCCCTTAAGAACGGCAAAAAAGAGAATATTGACAGAGAGTACTCAGAGACTGACATACTTGAGTCAGACGTACGCGACAAAAAGAACTTCGCAGTTCTGAAGAAAATTTACCTGAATGAAAACGTGAATAGCGAAGAACTGGAGGTGTTTAGAAACGATATCAAGAAAAAACTCAATAAAATCAACTCACTGAAATACAGTTTTGAGAAGAATAAAGCTAACTACCAGAAAATCAACGAGAACAATATCGAGAAGGTGGAAGGTAAGTCCAAAAGAAATATAATTTACGACTATTACAGAGAATCTGCGAAAAGGGACGCGTACGTATCCAATGTTAAGGAGGCATTTGATAAGCTTTACAAAGAAGAAGATATTGCTAAGTTGGTACTTGAAATAGAAAACCTCACTAAACTCGAGAAATACAAAATCCGAGAGTTTTACCACGAGATAATTGGCCGAAAAAATGATAAAGAGAATTTTGCTAAGATTATATACGAGGAAATACAAAATGTAAACAACATGAAGGAGCTTATAGAAAAGGTCCCGGATATGTCTGAACTGAAAAAATCACAAGTTTTTTACAAATACTACCTCGACAAGGAGGAGCTGAACGACAAGAATATCAAATATGCCTTCTGCCACTTTGTCGAAATTGAGATGAGTCAACTGCTCAAGAACTATGTGTACAAGCGACTCTCCAACATTAGCAATGACAAAATCAAACGAATTTTCGAATACCAGAATTTGAAGAAGTTGATAGAAAACAAGCTGCTTAATAAACTCGATACTTACGTTCGGAACTGTGGGAAATATAACTATTACCTTCAGGATGGCGAAATAGCCACGTCAGACTTTATTGCACGGAACAGGCAAAATGAGGCCTTTCTGAGGAATATTATCGGAGTCAGCAGTGTCGCTTATTTTAGTCTGCGGAACATCCTGGAGACAGAGAACGAAAACGATATAACTGGACGCATGCGCGGAAAGACGGTAAAAAATAATAAAGGTGAAGAAAAGTACGTTAGTGGGGAGGTAGATAAGATTTATAATGAAAATAAGAAGAACGAAGTTAAAGAGAACTTGAAGATGTTCTACAGTTATGATTTCAATATGGACAACAAGAACGAAATCGAAGATTTTTTTGCTAATATTGACGAAGCTATCTCCAGCATCAGGCATGGTATAGTGCACTTCAATCTTGAGTTGGAAGGCAAGGACATTTTTGCGTTCAAGAACATCGCCCCAAGCGAAATAAGCAAAAAGATGTTCCAGAACGAGATAAATGAGAAGAAATTGAAATTGAAAATCTTCCGGCAATTGAACTCAGCCAACGTTTTCAGATACCTCGAAAAGTATAAGATCCTCAATTACTTGAAACGCACACGATTCGAATTTGTTAACAAGAATATTCCGTTCGTCCCTAGCTTTACCAAGCTTTATAGCCGCATCGATGATCTGAAGAATTCCCTTGGGATATACTGGAAGACACCTAAAACTAATGACGATAATAAAACGAAGGAGATAATCGATGCACAGATATACTTGTTGAAAAATATATACTACGGCGAGTTTCTTAATTATTTCATGTCAAATAATGGGAATTTTTTTGAAATTTCAAAAGAAATTATAGAACTGAATAAGAACGACAAGCGCAACCTTAAGACTGGCTTTTACAAACTTCAGAAATTTGAGGACATCCAAGAAAAGATCCCGAAGGAGTATTTGGCGAACATTCAAAGCCTTTACATGATAAATGCCGGGAACCAGGATGAAGAAGAAAAAGACACCTATATTGATTTTATCCAGAAAATCTTTTTGAAGGGTTTTATGACTTACCTCGCGAACAATGGAAGACTCAGCTTGATCTATATAGGCAGTGACGAGGAGACTAACACGAGTTTGGCCGAAAAGAAGCAGGAGTTTGATAAATTCCTTAAAAAGTATGAACAAAATAATAACATTAAGATACCATACGAAATAAACGAGTTTCTTCGCGAAATCAAATTGGGAAACATCCTTAAGTATACGGAGAGGCTTAACATGTTTTACCTTATCCTCAAACTCCTCAACCACAAGGAGCTCACCAACCTGAAAGGGAGCCTCGAAAAGTACCAGTCCGCGAATAAAGAGGAGGCCTTCTCCGACCAACTTGAACTTATTAATCTTCTGAACCTGGACAATAATAGAGTCACGGAAGATTTTGAGTTGGAGGCGGATGAAATTGGCAAATTCCTTGATTTTAATGGCAATAAAGTCAAAGACAACAAAGAATTGAAGAAATTCGACACTAACAAGATATATTTTGACGGGGAGAATATCATCAAACATAGAGCCTTTTACAACATCAAAAAATATGGGATGCTCAATTTGCTTGAAAAAATAGCTGATAAGGCCGGGTACAAGATCTCAATCGAAGAACTGAAAAAATACTCTAACAAGAAGAATGAGATTGAGAAAAATCACAAGATGCAAGAAAACCTCCACCGCAAGTATGCGCGGCCTCGCAAGGACGAGAAGTTCACAGATGAAGACTACGAGTCTTACAAGCAGGCCATCGAAAATATCGAGGAATATACTCACCTCAAAAACAAGGTAGAATTTAATGAACTCAATCTGCTGCAAGGGCTGCTCCTTCGCATCCTTCACAGGCTCGTGGGATACACGTCCATTTGGGAACGCGATTTGCGGTTTCGGCTCAAGGGGGAATTTCCGGAGAATCAGTATATCGAAGAAATATTCAACTTCGAGAACAAAAAGAATGTGAAGTATAAGGGTGGACAGATCGTAGAAAAGTATATTAAGTTTTACAAGGAACTGCACCAGAACGATGAGGTCAAGATCAATAAATACAGCTCCGCTAATATCAAGGTGCTCAAACAGGAAAAAAAGGACTTGTACATTCGAAACTATATAGCGCACTTTAACTATATCCCCCATGCTGAAATATCACTCCTTGAAGTCTTGGAAAACCTCAGAAAACTTTTGAGCTACGATCGAAAGCTGAAGAACGCAGTCATGAAATCTGTTGTTGACATACTCAAGGAGTATGGATTTGTCGCAACGTTTAAGATCGGGGCGGACAAAAAGATCGGAATTCAGACGCTCGAATCAGAAAAGATTGTGCATCTGAAGAACCTCAAAAAGAAAAAGTTGATGACAGACAGGAACTCCGAAGAATTGTGCAAGCTCGTGAAGATTATGTTTGAGTATAAAATGGAAGAAAAGAAAAGTGAAAACggatacccttatgacgtacctgactatgcttaaggatccctcccccccccctaacgttactggccgaagccgcttggaataaggccggtgtgcgtttgtctatatgttattttccaccatattgccgtcttttggcaatgtgagggcccggaaacctggccctgtcttcttgacgagcattcctaggggtctttcccctctcgccaaaggaatgcaaggtctgttgaatgtcgtgaaggaagcagttcctctggaagcttcttgaagacaaacaacgtctgtagcgaccctttgcaggcagcggaaccccccacctggcgacaggtgcctctgcggccaaaagccacgtgtataagatacacctgcaaaggcggcacaaccccagtgccacgttgtgagttggatagttgtggaaagagtcaaatggctctcctcaagcgtattcaacaaggggctgaaggatgcccagaaggtaccccattgtatgggatctgatctggggcctcggtgcacatgctttacatgtgtttagtcgaggttaaaaaaacgtctaggccccccgaaccacggggacgtggttttcctttgaaaaacacgaTGATAATATGGCCACACATatgagcgagctgattaaggagaacatgcacatgaagctgtacatggagggcaccgtggacaaccatcacttcaagtgcacatccgagggcgaaggcaagccctacgagggcacccagaccatgagaatcaaggtggtcgagggcggccctctccccttcgccttcgacatcctggctactagcttcctctacggcagcaagaccttcatcaaccacacccagggcatccccgacttcttcaagcagtccttccctgagggcttcacatgggagagagtcaccacatacgaagacgggggcgtgctgaccgctacccaggacaccagcctccaggacggctgcctcatctacaacgtcaagatcagaggggtgaacttcacatccaacggccctgtgatgcagaagaaaacactcggctgggaggccttcaccgagacgctgtaccccgctgacggcggcctggaaggcagaaacgacatggccctgaagctcgtgggcgggagccatctgatcgcaaacatcaagaccacatatagatccaagaaacccgctaagaacctcaagatgcctggcgtctactatgtggactacagactggaaagaatcaaggaggccaacaacgagacctacgtcgagcagcacgaggtggcagtggccagatactgcgacctccctagcaaactggggcacaagcttaatggatccggcgcaacaaacttctctctgctgaaacaagccggagatgtcgaagagaatcctggaccgatggccaagcctttgtctcaagaagaatccaccctcattgaaagagcaacggctacaatcaacagcatccccatctctgaagactacagcgtcgccagcgcagctctctctagcgacggccgcatcttcactggtgtcaatgtatatcattttactgggggaccttgtgcagaactcgtggtgctgggcactgctgctgctgcggcagctggcaacctgacttgtatcgtcgcgatcggaaatgagaacaggggcatcttgagcccctgcggacggtgccgacaggtgcttctcgatctgcatcctgggatcaaagccatagtgaaggacagtgatggacagccgacggcagttgggattcgtgaattgctgccctctggttatgtgtgggagggctaagATCGATAGATCCTAATCAACCTctggattacaaaatttgtgaaagattgactggtattcttaactatgttgctccttttacgctatgtggatacgctgctttaatgcctttgtatcatgctattgcttcccgtatggctttcattttctcctccttgtataaatcctggttgctgtctctttatgaggagttgtggcccgttgtcaggcaacgtggcgtggtgtgcactgtgtttgctgacgcaacccccactggttggggcattgccaccacctgtcagctcctttccgggactttcgctttccccctccctattgccacggcggaactcatcgccgcctgccttgcccgctgctggacaggggctcggctgttgggcactgacaattccgtggtgttgtcggggaaatcatcgtcctttccttggctgctcgcctgtgttgccacctggattctgcgcgggacgtccttctgctacgtcccttcggccctcaatccagcggaccttccttcccgcggcctgctgccggctctgcggcctcttccgcgtcttcgccttcgccctcagacgagtcggatctccctttgggccgcctccccgcctggtacctttaagaccaatgacttacaaggcagctgtagatcttagccactttttaaaagaaaaggggggactggaagggctaattcactcccaacgaagacaagatcacctgcaggacaggcgcgccctgctttttgcttgtactgggtctctctggttagaccagatctgagcctgggagctctctggctaactagggaacccactgcttaagcctcaataaagcttgccttgagtgcttcaagtagtgtgtgcccgtctgttgtgtgactctggtaactagagatccctcagacccttttagtcagtgtggaaaatctctagcacccgggcgattaaggaaagggctagatcattcttgaagacgaaagggcctcgtgatacgcctatttttataggttaatgtcatgataataatggtttcttagacgtcaggtggcacttttcggggaaatgtgcgcggaacccctatttgtttatttttctaaatacattcaaatatgtatccgctcatgagacaataaccctgataaatgcttcaataatattgaaaaaggaagagtatgagtattcaacatttccgtgtcgcccttattcccttttttgcggcattttgccttcctgtttttgctcacccagaaacgctggtgaaagtaaaagatgctgaagatcagttgggtgcacgagtgggttacatcgaactggatctcaacagcggtaagatccttgagagttttcgccccgaagaacgttttccaatgatgagcacttttaaagttctgctatgtggcgcggtattatcccgtgttgacgccgggcaagagcaactcggtcgccgcatacactattctcagaatgacttggttgagtactcaccagtcacagaaaagcatcttacggatggcatgacagtaagagaattatgcagtgctgccataaccatgagtgataacactgcggccaacttacttctgacaacgatcggaggaccgaaggagctaaccgcttttttgcacaacatgggggatcatgtaactcgccttgatcgttgggaaccggagctgaatgaagccataccaaacgacgagcgtgacaccacgatgcctgtagcaatggcaacaacgttgcgcaaactattaactggcgaactacttactctagcttcccggcaacaattaatagactggatggaggcggataaagttgcaggaccacttctgcgctcggcccttccggctggctggtttattgctgataaatctggagccggtgagcgtgggtctcgcggtatcattgcagcactggggccagatggtaagccctcccgtatcgtagttatctacacgacggggagtcaggcaactatggatgaacgaaatagacagatcgctgagataggtgcctcactgattaagcattggtaactgtcagaccaagtttactcatatatactttagattgatttaaaacttcatttttaatttaaaaggatctaggtgaagatcctttttgataatctcatgaccaaaatcccttaacgtgagttttcgttccactgagcgtcagaccccgtagaaaagatcaaaggatcttcttgagatcctttttttctgcgcgtaatctgctgcttgcaaacaaaaaaaccaccgctaccagcggtggtttgtttgccggatcaagagctaccaactctttttccgaaggtaactggcttcagcagagcgcagataccaaatactgttcttctagtgtagccgtagttaggccaccacttcaagaactctgtagcaccgcctacatacctcgctctgctaatcctgttaccagtggctgctgccagtggcgataagtcgtgtcttaccgggttggactcaagacgatagttaccggataaggcgcagcggtcgggctgaacggggggttcgtgcacacagcccagcttggagcgaacgacctacaccgaactgagatacctacagcgtgagctatgagaaagcgccacgcttcccgaagggagaaaggcggacaggtatccggtaagcggcagggtcggaacaggagagcgcacgagggagcttccagggggaaacgcctggtatctttatagtcctgtcgggtttcgccacctctgacttgagcgtcgatttttgtgatgctcgtcaggggggcggagcctatggaaaaacgccagcaacgcggcctttttacggttcctggccttttgctggccttttgctcacatgttctttcctgcgttatcccctgattctgtggataaccgtattaccgcctttgagtgagctgataccgctcgccgcagccgaacgaccgagcgcagcgagtcagtgagcgaggaagcggaagagcgcccaatacgcaaaccgcctctccccgcgcgttggccgattcattaatgcagcaagctcatggctgactaattttttttatttatgcagaggccgaggccgcctcggcctctgagctattccagaagtagtgaggaggcttttttggaggcctaggcttttgcaaaaagctccccgtggcacgacaggtttcccgactggaaagcgggcagtgagcgcaacgcaattaatgtgagttagctcactcattaggcaccccaggctttacactttatgcttccggctcgtatgttgtgtggaattgtgagcggataacaatttcacacaggaaacagctatgacatgattacgaatttcacaaataaagcatttttttcactgcattctagttgtggtttgtccaaactcatcaatgtatcttatcatgtctggatcaactggataactcaagctaaccaaaatcatcccaaacttcccaccccataccctattaccactgccaattacctgtggtttcatttactctaaacctgtgattcctctgaattattttcattttaaagaaattgtatttgttaaatatgtactacaaacttagtagt

#### pLentiviral_EF-1a-RfxCas13d-IRES-TagBFP-P2A-BSD

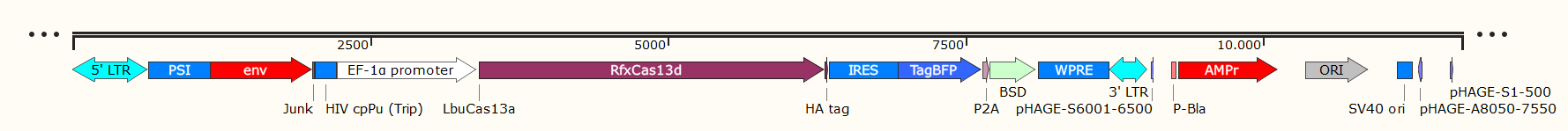

tggaagggctaattcactcccaaagaagacaagatatccttgatctgtggatctaccacacacaaggctacttccctgattagcagaactacacaccagggccaggggtcagatatccactgacctttggatggtgctacaagctagtaccagttgagccagataaggtagaagaggccaataaaggagagaacaccagcttgttacaccctgtgagcctgcatgggatggatgacccggagagagaagtgttagagtggaggtttgacagccgcctagcatttcatcacgtggcccgagagctgcatccggagtacttcaagaactgctgatatcgagcttgctacaagggactttccgctggggactttccagggaggcgtggcctgggcgggactggggagtggcgagccctcagatcctgcatataagcagctgctttttgcctgtactgggtctctctggttagaccagatctgagcctgggagctctctggctaactagggaacccactgcttaagcctcaataaagcttgccttgagtgcttcaagtagtgtgtgcccgtctgttgtgtgactctggtaactagagatccctcagacccttttagtcagtgtggaaaatctctagcagtggcgcccgaacagggacttgaaagcgaaagggaaaccagaggagctctctcgacgcaggactcggcttgctgaagcgcgcacggcaagaggcgaggggcggcgactggtgagtacgccaaaaattttgactagcggaggctagaaggagagagatgggtgcgagagcgtcagtattaagcgggggagaattagatcgcgatgggaaaaaattcggttaaggccagggggaaagaaaaaatataaattaaaacatatagtatgggcaagcagggagctagaacgattcgcagttaatcctggcctgttagaaacatcagaaggctgtagacaaatactgggacagctacaaccatcccttcagacaggatcagaagaacttagatcattatataatacagtagcaaccctctattgtgtgcatcaaaggatagagataaaagacaccaaggaagctttagacaagatagaggaagagcaaaacaaaagtaagaccaccgcacagcaagcggccggccgctgatcttcagacctggaggaggagatatgagggacaattggagaagtgaattatataaatataaagtagtaaaaattgaaccattaggagtagcacccaccaaggcaaagagaagagtggtgcagagagaaaaaagagcagtgggaataggagctttgttccttgggttcttgggagcagcaggaagcactatgggcgcagcgtcaatgacgctgacggtacaggccagacaattattgtctggtatagtgcagcagcagaacaatttgctgagggctattgaggcgcaacagcatctgttgcaactcacagtctggggcatcaagcagctccaggcaagaatcctggctgtggaaagatacctaaaggatcaacagctcctggggatttggggttgctctggaaaactcatttgcaccactgctgtgccttggaatgctagttggagtaataaatctctggaacagatttggaatcacacgacctggatggagtgggacagagaaattaacaattacacaagcttaatacactccttaattgaagaatcgcaaaaccagcaagaaaagaatgaacaagaattattggaattagataaatgggcaagtttgtggaattggtttaacataacaaattggctgtggtatataaaattattcataatgatagtaggaggcttggtaggtttaagaatagtttttgctgtactttctatagtgaatagagttaggcagggatattcaccattatcgtttcagacccacctcccaaccccgaggggacccgacaggcccgaaggaatagaagaagaaggtggagagagagacagagacagatccattcgattagtgaacggatctcgacggtatcgccgaattcacaaatggcagtattcatccacaattttaaaagaaaaggggggattggggggtacagtgcaggggaaagaatagtagacataatagcaacagacatacaaactaaagaattacaaaaacaaattacaaaaattcaaaattttcgggtttattacagggacagcagagatccagtttggactagtcgtgaggctccggtgcccgtcagtgggcagagcgcacatcgcccacagtccccgagaagttggggggaggggtcggcaattgaaccggtgcctagagaaggtggcgcggggtaaactgggaaagtgatgtcgtgtactggctccgcctttttcccgagggtgggggagaaccgtatataagtgcagtagtcgccgtgaacgttctttttcgcaacgggtttgccgccagaacacaggtaagtgccgtgtgtggttcccgcgggcctggcctctttacgggttatggcccttgcgtgccttgaattacttccacctggctgcagtacgtgattcttgatcccgagcttcgggttggaagtgggtgggagagttcgaggccttgcgcttaaggagccccttcgcctcgtgcttgagttgaggcctggcctgggcgctggggccgccgcgtgcgaatctggtggcaccttcgcgcctgtctcgctgctttcgataagtctctagccatttaaaatttttgatgacctgctgcgacgctttttttctggcaagatagtcttgtaaatgcgggccaagatctgcacactggtatttcggtttttggggccgcgggcggcgacggggcccgtgcgtcccagcgcacatgttcggcgaggcggggcctgcgagcgcggccaccgagaatcggacgggggtagtctcaagctggccggcctgctctggtgcctggcctcgcgccgccgtgtatcgccccgccctgggcggcaaggctggcccggtcggcaccagttgcgtgagcggaaagatggccgcttcccggccctgctgcagggagctcaaaatggaggacgcggcgctcgggagagcgggcgggtgagtcacccacacaaaggaaaagggcctttccgtcctcagccgtcgcttcatgtgactccacggagtaccgggcgccgtccaggcacctcgattagttctcgagcttttggagtacgtcgtctttaggttggggggaggggttttatgcgatggagtttccccacactgagtgggtggagactgaagttaggccagcttggcacttgatgtaattctccttggaatttgccctttttgagtttggatcttggttcattctcaagcctcagacagtggttcaaagtttttttcttccatttcaggtgtcgtgaagcggccgccaccATGatcgaaaaaaaaaagtccttcgccaagggcatgggcgtgaagtccacactcgtgtccggctccaaagtgtacatgacaaccttcgccgaaggcagcgacgccaggctggaaaagatcgtggagggcgacagcatcaggagcgtgaatgagggcgaggccttcagcgctgaaatggccgataaaaacgccggctataagatcggcaacgccaaattcagccatcctaagggctacgccgtggtggctaacaaccctctgtatacaggacccgtccagcaggatatgctcggcctgaaggaaactctggaaaagaggtacttcggcgagagcgctgatggcaatgacaatatttgtatccaggtgatccataacatcctggacattgaaaaaatcctcgccgaatacattaccaacgccgcctacgccgtcaacaatatctccggcctggataaggacattattggattcggcaagttctccacagtgtatacctacgacgaattcaaagaccccgagcaccatagggccgctttcaacaataacgataagctcatcaacgccatcaaggcccagtatgacgagttcgacaacttcctcgataaccccagactcggctatttcggccaggcctttttcagcaaggagggcagaaattacatcatcaattacggcaacgaatgctatgacattctggccctcctgagcggactgaggcactgggtggtccataacaacgaagaagagtccaggatctccaggacctggctctacaacctcgataagaacctcgacaacgaatacatctccaccctcaactacctctacgacaggatcaccaatgagctgaccaactccttctccaagaactccgccgccaacgtgaactatattgccgaaactctgggaatcaaccctgccgaattcgccgaacaatatttcagattcagcattatgaaagagcagaaaaacctcggattcaatatcaccaagctcagggaagtgatgctggacaggaaggatatgtccgagatcaggaaaaatcataaggtgttcgactccatcaggaccaaggtctacaccatgatggactttgtgatttataggtattacatcgaagaggatgccaaggtggctgccgccaataagtccctccccgataatgagaagtccctgagcgagaaggatatctttgtgattaacctgaggggctccttcaacgacgaccagaaggatgccctctactacgatgaagctaatagaatttggagaaagctcgaaaatatcatgcacaacatcaaggaatttaggggaaacaagacaagagagtataagaagaaggacgcccctagactgcccagaatcctgcccgctggccgtgatgtttccgccttcagcaaactcatgtatgccctgaccatgttcctggatggcaaggagatcaacgacctcctgaccaccctgattaataaattcgataacatccagagcttcctgaaggtgatgcctctcatcggagtcaacgctaagttcgtggaggaatacgcctttttcaaagactccgccaagatcgccgatgagctgaggctgatcaagtccttcgctagaatgggagaacctattgccgatgccaggagggccatgtatatcgacgccatccgtattttaggaaccaacctgtcctatgatgagctcaaggccctcgccgacaccttttccctggacgagaacggaaacaagctcaagaaaggcaagcacggcatgagaaatttcattattaataacgtgatcagcaataaaaggttccactacctgatcagatacggtgatcctgcccacctccatgagatcgccaaaaacgaggccgtggtgaagttcgtgctcggcaggatcgctgacatccagaaaaaacagggccagaacggcaagaaccagatcgacaggtactacgaaacttgtatcggaaaggataagggcaagagcgtgagcgaaaaggtggacgctctcacaaagatcatcaccggaatgaactacgaccaattcgacaagaaaaggagcgtcattgaggacaccggcagggaaaacgccgagagggagaagtttaaaaagatcatcagcctgtacctcaccgtgatctaccacatcctcaagaatattgtcaatatcaacgccaggtacgtcatcggattccattgcgtcgagcgtgatgctcaactgtacaaggagaaaggctacgacatcaatctcaagaaactggaagagaagggattcagctccgtcaccaagctctgcgctggcattgatgaaactgcccccgataagagaaaggacgtggaaaaggagatggctgaaagagccaaggagagcattgacagcctcgagagcgccaaccccaagctgtatgccaattacatcaaatacagcgacgagaagaaagccgaggagttcaccaggcagattaacagggagaaggccaaaaccgccctgaacgcctacctgaggaacaccaagtggaatgtgatcatcagggaggacctcctgagaattgacaacaagacatgtaccctgttcagaaacaaggccgtccacctggaagtggccaggtatgtccacgcctatatcaacgacattgccgaggtcaattcctacttccaactgtaccattacatcatgcagagaattatcatgaatgagaggtacgagaaaagcagcggaaaggtgtccgagtacttcgacgctgtgaatgacgagaagaagtacaacgataggctcctgaaactgctgtgtgtgcctttcggctactgtatccccaggtttaagaacctgagcatcgaggccctgttcgataggaacgaggccgccaagttcgacaaggagaaaaagaaggtgtccggcaattccggatacccttatgacgtacctgactatgcttaaggatccctcccccccccctaacgttactggccgaagccgcttggaataaggccggtgtgcgtttgtctatatgttattttccaccatattgccgtcttttggcaatgtgagggcccggaaacctggccctgtcttcttgacgagcattcctaggggtctttcccctctcgccaaaggaatgcaaggtctgttgaatgtcgtgaaggaagcagttcctctggaagcttcttgaagacaaacaacgtctgtagcgaccctttgcaggcagcggaaccccccacctggcgacaggtgcctctgcggccaaaagccacgtgtataagatacacctgcaaaggcggcacaaccccagtgccacgttgtgagttggatagttgtggaaagagtcaaatggctctcctcaagcgtattcaacaaggggctgaaggatgcccagaaggtaccccattgtatgggatctgatctggggcctcggtgcacatgctttacatgtgtttagtcgaggttaaaaaaacgtctaggccccccgaaccacggggacgtggttttcctttgaaaaacacgaTGATAATATGGCCACACATatgagcgagctgattaaggagaacatgcacatgaagctgtacatggagggcaccgtggacaaccatcacttcaagtgcacatccgagggcgaaggcaagccctacgagggcacccagaccatgagaatcaaggtggtcgagggcggccctctccccttcgccttcgacatcctggctactagcttcctctacggcagcaagaccttcatcaaccacacccagggcatccccgacttcttcaagcagtccttccctgagggcttcacatgggagagagtcaccacatacgaagacgggggcgtgctgaccgctacccaggacaccagcctccaggacggctgcctcatctacaacgtcaagatcagaggggtgaacttcacatccaacggccctgtgatgcagaagaaaacactcggctgggaggccttcaccgagacgctgtaccccgctgacggcggcctggaaggcagaaacgacatggccctgaagctcgtgggcgggagccatctgatcgcaaacatcaagaccacatatagatccaagaaacccgctaagaacctcaagatgcctggcgtctactatgtggactacagactggaaagaatcaaggaggccaacaacgagacctacgtcgagcagcacgaggtggcagtggccagatactgcgacctccctagcaaactggggcacaagcttaatggatccggcgcaacaaacttctctctgctgaaacaagccggagatgtcgaagagaatcctggaccgatggccaagcctttgtctcaagaagaatccaccctcattgaaagagcaacggctacaatcaacagcatccccatctctgaagactacagcgtcgccagcgcagctctctctagcgacggccgcatcttcactggtgtcaatgtatatcattttactgggggaccttgtgcagaactcgtggtgctgggcactgctgctgctgcggcagctggcaacctgacttgtatcgtcgcgatcggaaatgagaacaggggcatcttgagcccctgcggacggtgccgacaggtgcttctcgatctgcatcctgggatcaaagccatagtgaaggacagtgatggacagccgacggcagttgggattcgtgaattgctgccctctggttatgtgtgggagggctaagATCGATAGATCCTAATCAACCTctggattacaaaatttgtgaaagattgactggtattcttaactatgttgctccttttacgctatgtggatacgctgctttaatgcctttgtatcatgctattgcttcccgtatggctttcattttctcctccttgtataaatcctggttgctgtctctttatgaggagttgtggcccgttgtcaggcaacgtggcgtggtgtgcactgtgtttgctgacgcaacccccactggttggggcattgccaccacctgtcagctcctttccgggactttcgctttccccctccctattgccacggcggaactcatcgccgcctgccttgcccgctgctggacaggggctcggctgttgggcactgacaattccgtggtgttgtcggggaaatcatcgtcctttccttggctgctcgcctgtgttgccacctggattctgcgcgggacgtccttctgctacgtcccttcggccctcaatccagcggaccttccttcccgcggcctgctgccggctctgcggcctcttccgcgtcttcgccttcgccctcagacgagtcggatctccctttgggccgcctccccgcctggtacctttaagaccaatgacttacaaggcagctgtagatcttagccactttttaaaagaaaaggggggactggaagggctaattcactcccaacgaagacaagatcacctgcaggacaggcgcgccctgctttttgcttgtactgggtctctctggttagaccagatctgagcctgggagctctctggctaactagggaacccactgcttaagcctcaataaagcttgccttgagtgcttcaagtagtgtgtgcccgtctgttgtgtgactctggtaactagagatccctcagacccttttagtcagtgtggaaaatctctagcacccgggcgattaaggaaagggctagatcattcttgaagacgaaagggcctcgtgatacgcctatttttataggttaatgtcatgataataatggtttcttagacgtcaggtggcacttttcggggaaatgtgcgcggaacccctatttgtttatttttctaaatacattcaaatatgtatccgctcatgagacaataaccctgataaatgcttcaataatattgaaaaaggaagagtatgagtattcaacatttccgtgtcgcccttattcccttttttgcggcattttgccttcctgtttttgctcacccagaaacgctggtgaaagtaaaagatgctgaagatcagttgggtgcacgagtgggttacatcgaactggatctcaacagcggtaagatccttgagagttttcgccccgaagaacgttttccaatgatgagcacttttaaagttctgctatgtggcgcggtattatcccgtgttgacgccgggcaagagcaactcggtcgccgcatacactattctcagaatgacttggttgagtactcaccagtcacagaaaagcatcttacggatggcatgacagtaagagaattatgcagtgctgccataaccatgagtgataacactgcggccaacttacttctgacaacgatcggaggaccgaaggagctaaccgcttttttgcacaacatgggggatcatgtaactcgccttgatcgttgggaaccggagctgaatgaagccataccaaacgacgagcgtgacaccacgatgcctgtagcaatggcaacaacgttgcgcaaactattaactggcgaactacttactctagcttcccggcaacaattaatagactggatggaggcggataaagttgcaggaccacttctgcgctcggcccttccggctggctggtttattgctgataaatctggagccggtgagcgtgggtctcgcggtatcattgcagcactggggccagatggtaagccctcccgtatcgtagttatctacacgacggggagtcaggcaactatggatgaacgaaatagacagatcgctgagataggtgcctcactgattaagcattggtaactgtcagaccaagtttactcatatatactttagattgatttaaaacttcatttttaatttaaaaggatctaggtgaagatcctttttgataatctcatgaccaaaatcccttaacgtgagttttcgttccactgagcgtcagaccccgtagaaaagatcaaaggatcttcttgagatcctttttttctgcgcgtaatctgctgcttgcaaacaaaaaaaccaccgctaccagcggtggtttgtttgccggatcaagagctaccaactctttttccgaaggtaactggcttcagcagagcgcagataccaaatactgttcttctagtgtagccgtagttaggccaccacttcaagaactctgtagcaccgcctacatacctcgctctgctaatcctgttaccagtggctgctgccagtggcgataagtcgtgtcttaccgggttggactcaagacgatagttaccggataaggcgcagcggtcgggctgaacggggggttcgtgcacacagcccagcttggagcgaacgacctacaccgaactgagatacctacagcgtgagctatgagaaagcgccacgcttcccgaagggagaaaggcggacaggtatccggtaagcggcagggtcggaacaggagagcgcacgagggagcttccagggggaaacgcctggtatctttatagtcctgtcgggtttcgccacctctgacttgagcgtcgatttttgtgatgctcgtcaggggggcggagcctatggaaaaacgccagcaacgcggcctttttacggttcctggccttttgctggccttttgctcacatgttctttcctgcgttatcccctgattctgtggataaccgtattaccgcctttgagtgagctgataccgctcgccgcagccgaacgaccgagcgcagcgagtcagtgagcgaggaagcggaagagcgcccaatacgcaaaccgcctctccccgcgcgttggccgattcattaatgcagcaagctcatggctgactaattttttttatttatgcagaggccgaggccgcctcggcctctgagctattccagaagtagtgaggaggcttttttggaggcctaggcttttgcaaaaagctccccgtggcacgacaggtttcccgactggaaagcgggcagtgagcgcaacgcaattaatgtgagttagctcactcattaggcaccccaggctttacactttatgcttccggctcgtatgttgtgtggaattgtgagcggataacaatttcacacaggaaacagctatgacatgattacgaatttcacaaataaagcatttttttcactgcattctagttgtggtttgtccaaactcatcaatgtatcttatcatgtctggatcaactggataactcaagctaaccaaaatcatcccaaacttcccaccccataccctattaccactgccaattacctgtggtttcatttactctaaacctgtgattcctctgaattattttcattttaaagaaattgtatttgttaaatatgtactacaaacttagtagt
