## Supplementary note 1 on orthologs for "Temporal dynamics of collateral RNA cleavage by LbuCas13a in human cells"

### Supplementary note on the collateral activity of different Cas13 orthologs

We initially compared the differed Cas13 orthologs in cells at 50 minutes after transfection, while targeting dEGFP (Supplementary figure 5B). However, from subsequent experiments we concluded that collateral RNA cleavage peaks at around 100 minutes after transfection (Figure 1A and C), and that targeting the 18S rRNA resulted in the highest collateral cleavage activity in all tested cell types (Figure 2). We wondered whether the other Cas13 orthologs were able to show detectable levels of collateral cleavage under these conditions. Therefore, we re-tested LwaCas13a, BzCas13b and RspCas13d, this time isolating total RNA after 100 minutes and targeting either GAPDH, dEGFP or 18S rRNA in HAP1 cells. Targeting GAPDH or dEGFP with LwaCas13a did not change the total RNA profile, however 18S rRNA targeting resulted in two additional peaks smaller in size than the 18S rRNA (Supplementary figure 2A). As 18S rRNA was the target RNA, this probably represents target RNA cleavage. Using BzCas13b resulted in a small extra peak between the 18S and 28S rRNA peak when targeting dEGFP or 18S rRNA (Supplementary figure 2B). Furthermore, RNAs smaller than 5S RNA and tRNA appeared to increase in abundance, which possibly represents the emergence of cleavage products. None of the samples using RspCas13d showed any alteration to the total RNA profile, suggesting that RspCas13d does not work well in human cells (Supplementary figure 2C). Thus, none of the other tested Cas13 orthologs were able to generate the dramatic RNA degradation pattern found when using LbuCas13a, although some collateral cleavage activity by LwaCas13a and BzCas13b cannot be completely ruled out.

Recently, it was reported that RfxCas13d exhibits collateral cleavage in eukaryotic cells (Shi *et al*, 2023; Li *et al*, 2023). Therefore, we explored how the *in vitro* and intracellular activity of RfxCas13d compared to the other orthologs tested in this study. First, we produced recombinant RfxCas13d protein (supplementary figure 1A) and assessed the collateral RNA cleavage activity of RfxCas13d *in vitro* with the RNaseAlert assay. RfxCas13d was not active in this assay for all three guide RNAs tested (Supplementary figure 16). It is possible that RfxCas13d, under these assay conditions, was not able to cleave the substrate provided by RNaseAlert. However, RfxCas13d has a similar cleavage preference as LbuCas13a, which seemingly has no issue cleaving the RNaseAlert reporter, both preferring a poly-U substrate (Brogan *et al*, 2021; Li *et al*, 2022; East-Seletsky *et al*, 2017). Alternatively, the RfxCas13d recombinant protein production might have had a very low active fraction. In any case, we cannot draw any conclusions about the *in vitro* activity of RfxCas13d. However, in another study LwaCas13a had higher activity than RfxCas13d *in vitro* (Li *et al*, 2022). This suggests that LbuCas13a has higher *in vitro* collateral RNA cleavage activity than RfxCas13d, as we find that LbuCas13a is much more active than LwaCas13a (Supplementary figure 1B).

Next we compared RfxCas13d and LbuCas13a side-by-side in HAP1-dEGFP cells. Although we were unable to produce enough RfxCas13d for iTOP delivery, we did have enough yield for RNP delivery by electroporation. RNA targeting with RfxCas13d did not alter the total RNA profile for any of the guide RNAs tested (targeting dEGFP, GAPDH, or 18s rRNA), while LbuCas13a showed the characteristic total RNA collateral cleavage pattern (Supplementary figure 3A). RfxCas13d showed a modest (not significant) increase in 5’ Cq values of the target RNA (Supplemental figure 5C), suggesting target RNA cleavage. However, there was no change in the Cq values of collateral RNA. Targeting with LbuCas13a resulted in increased 5’ Cq values for both the target and collateral RNA. In addition to RNP delivery by electroporation, we made a HAP1-dEGFP derived RfxCas13d expressing cell line. We did not find any evidence of collateral activity in this RfxCas13d protein expressing line (Supplementary figure 17). In conclusion, we could not find any evidence of collateral RNA cleavage activity by RfxCas13d. Other recent publications also report not finding indications of collateral RNA cleavage by RfxCas13d in cells (Tieu *et al*, 2024; Wessels *et al*, 2024). It is interesting to note that the recent reports of collateral cleavage by RfxCas13d in eukaryotic cell lines all use transient plasmid transfection, while above mentioned recent publications finding no collateral cleavage use lentiviral integration. This suggests that the intracellular (Rfx)Cas13 concentration is an important factor in whether collateral RNA cleavage occurs.

Altogether, these results highlight how situational the collateral RNA cleavage activity of Cas13 orthologs seems to be, depending on many factors such as delivery method, guide RNA, target RNA and cell line. This might explain the conflicting reports of collateral cleavage for orthologs such as LwaCas13a and RfxCas13d. Our work suggests that LbuCas13a may be the most reliable ortholog for the induction of collateral RNA cleavage in human cells, as we were able to find collateral cleavage in all tested cell lines, using different RNP delivery methods, and targeting different highly expressed transcripts.
