## Supplementary figures and images for "Temporal dynamics of collateral RNA cleavage by LbuCas13a in human cells"

### Supplementary figure 1

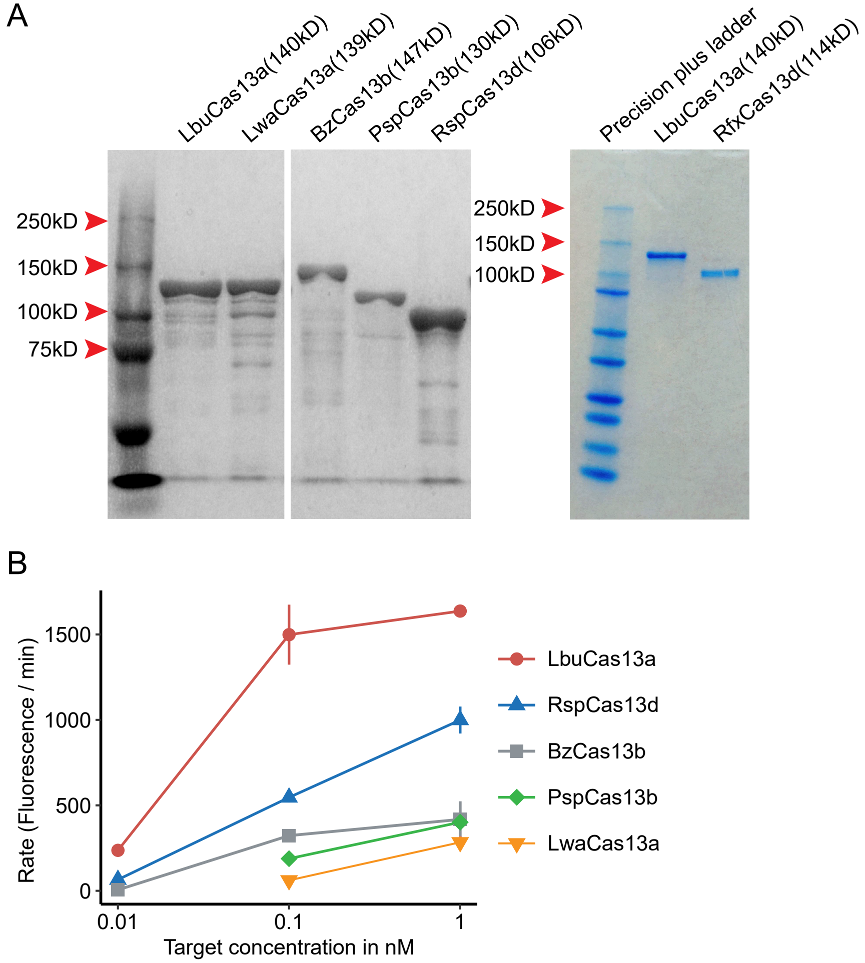


J. F. Bot et al. Supplementary figure 1

### Supplementary figure 2

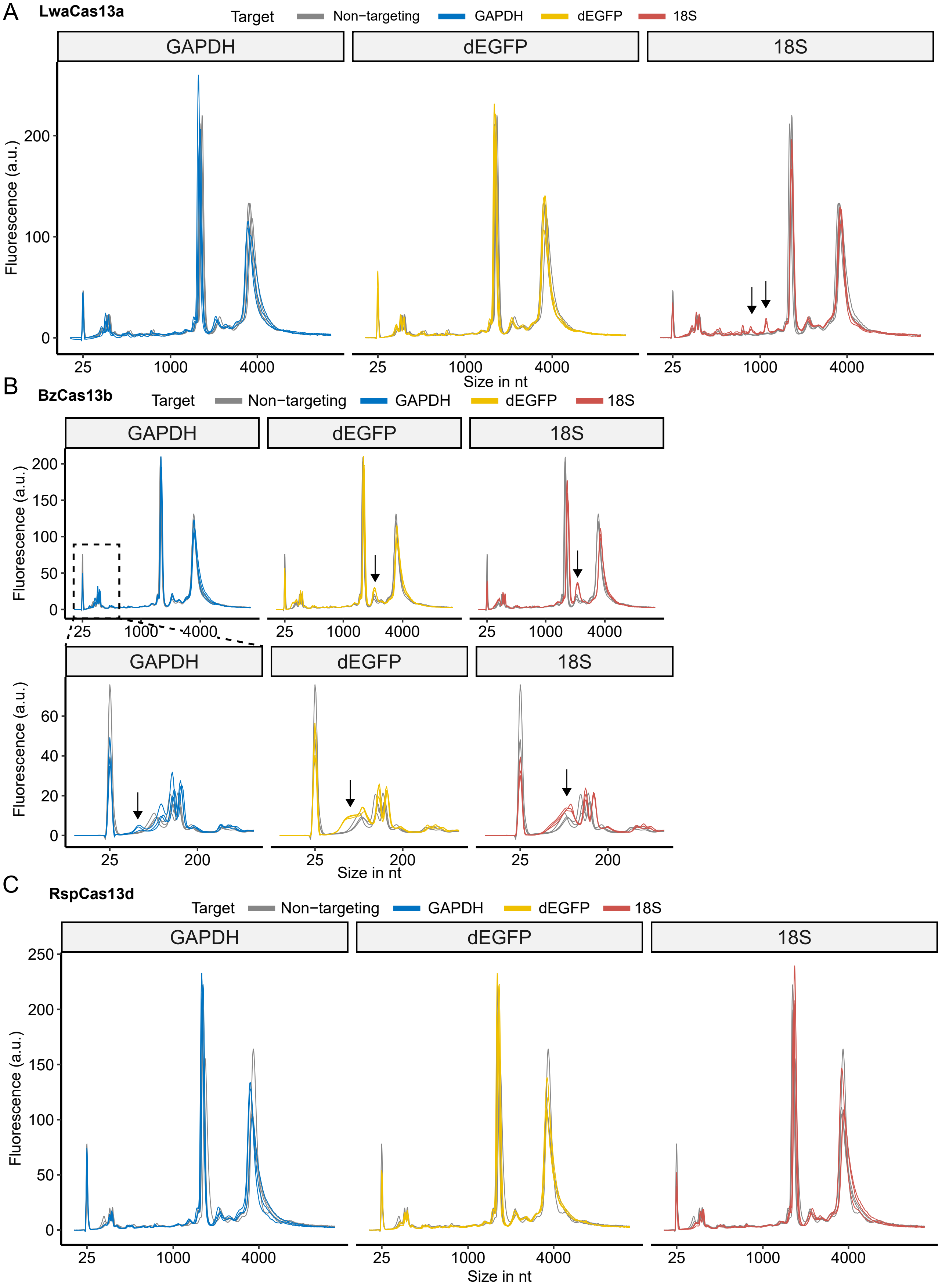


J. F. Bot et al. Supplementary figure 2

### Supplementary figure 3

A

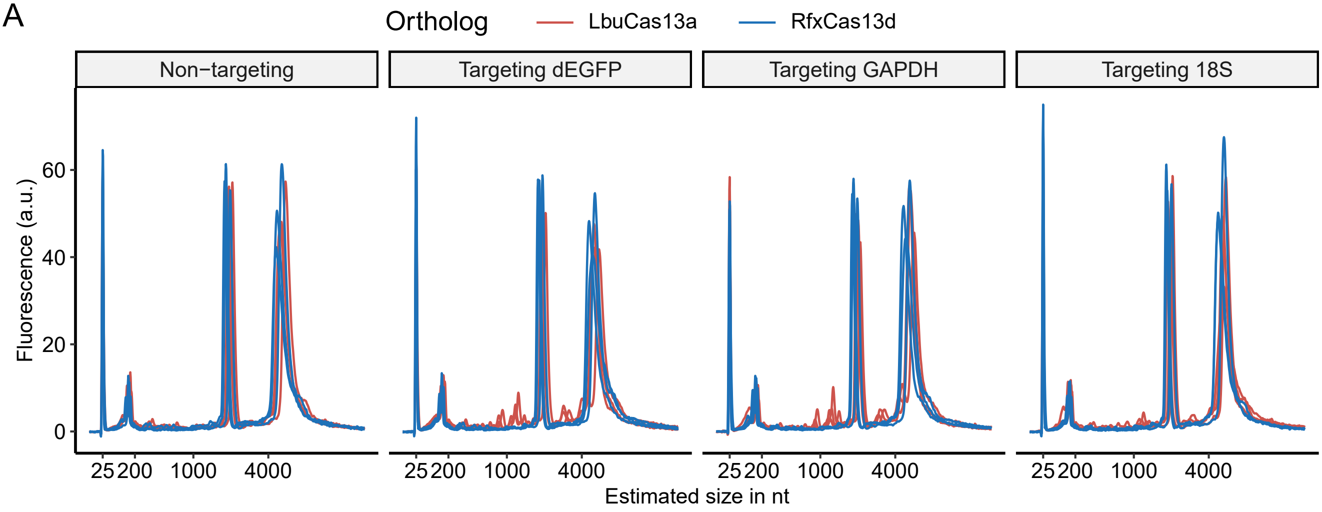


J. F. Bot et al. Supplementary figure 3

### Supplementary figure 4

A

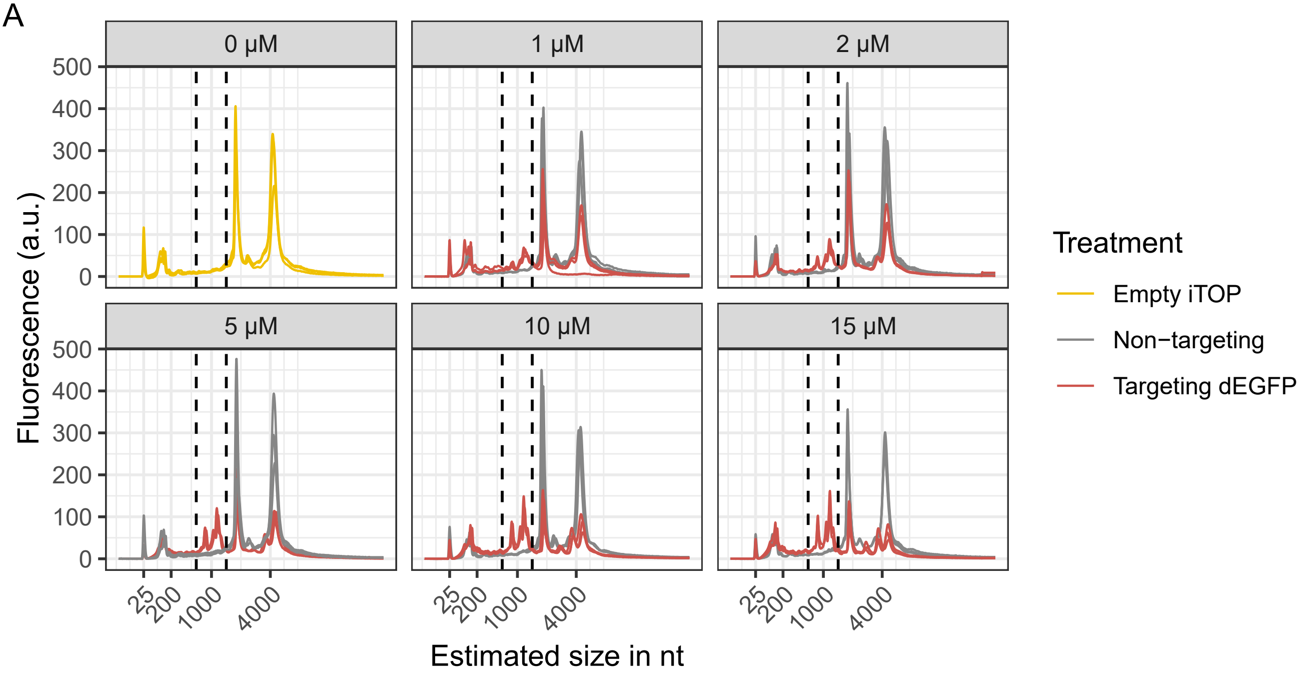


B

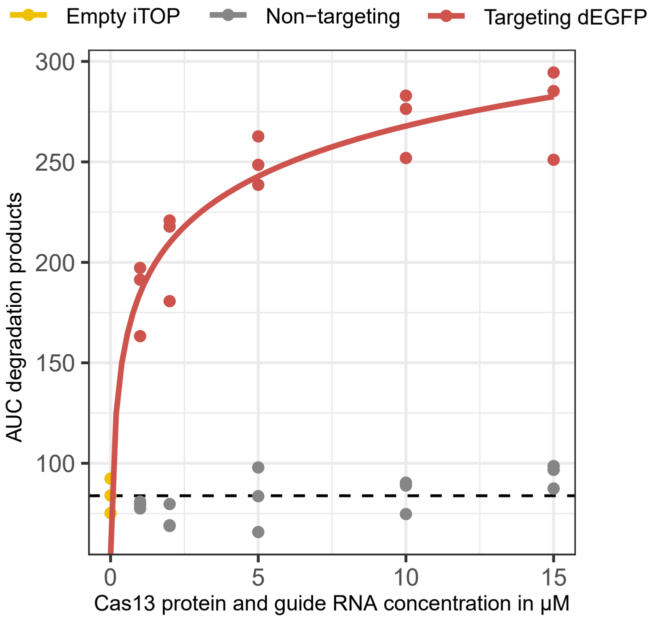


J. F. Bot et al. Supplementary figure 4

### Supplementary figure 5

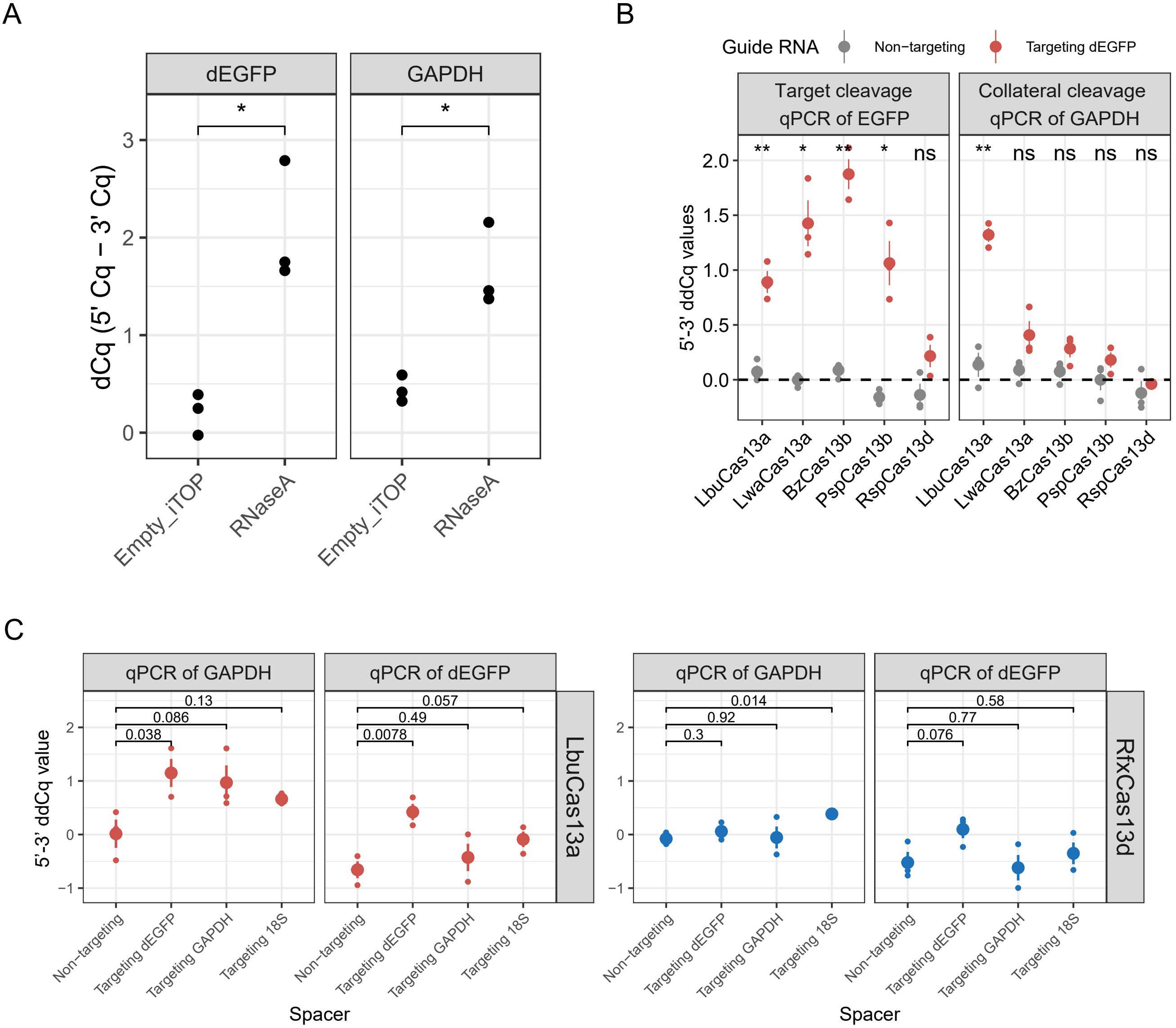

### Supplementary figure 6

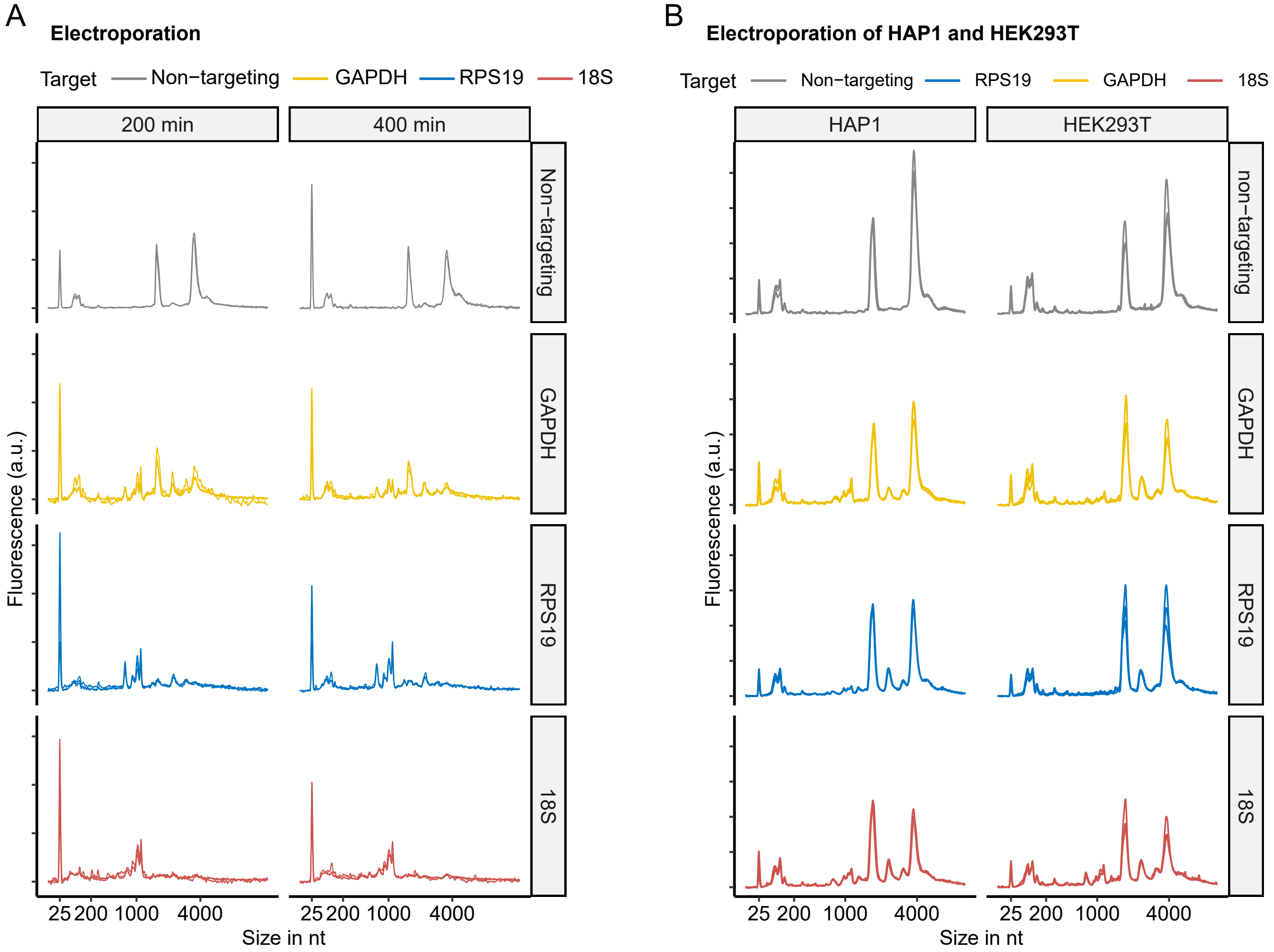


J. F. Bot et al. Supplementary figure 6

### Supplementary figure 7

A

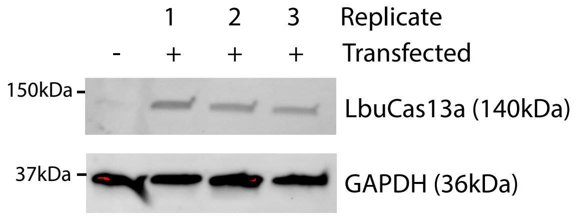


B

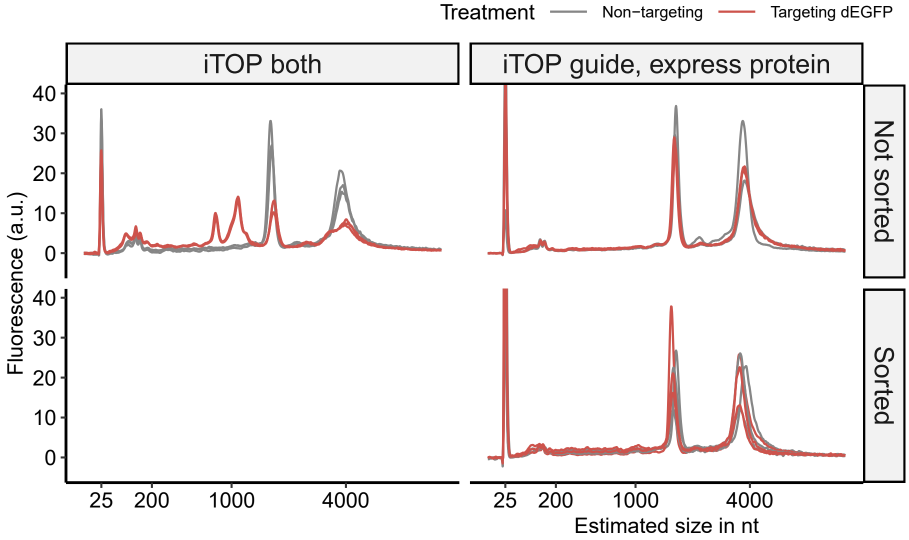


J. F. Bot et al. Supplementary figure 7

### Supplementary figure 8

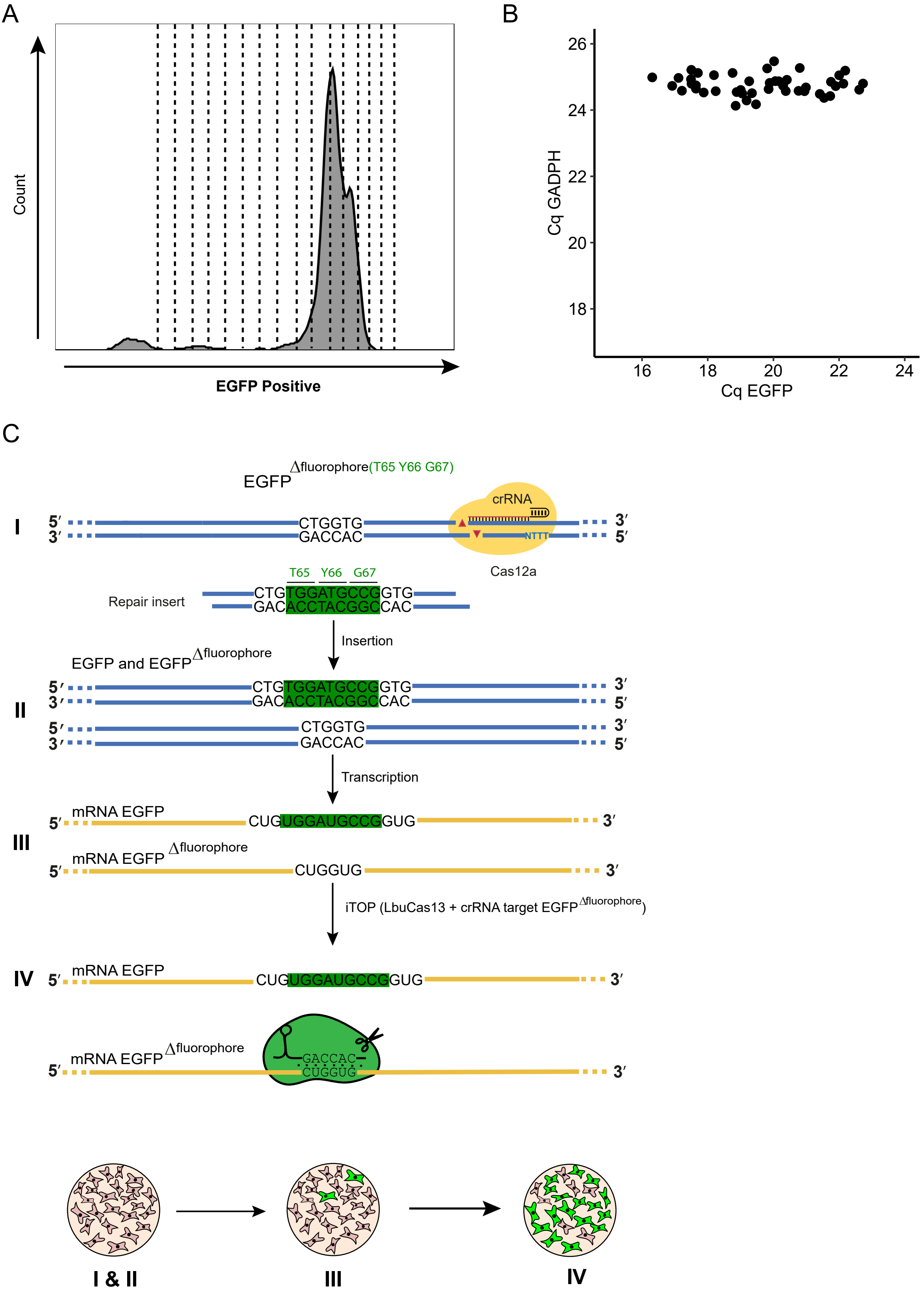


J. F. Bot et al. Supplementary figure 8

### Supplementary figure 9

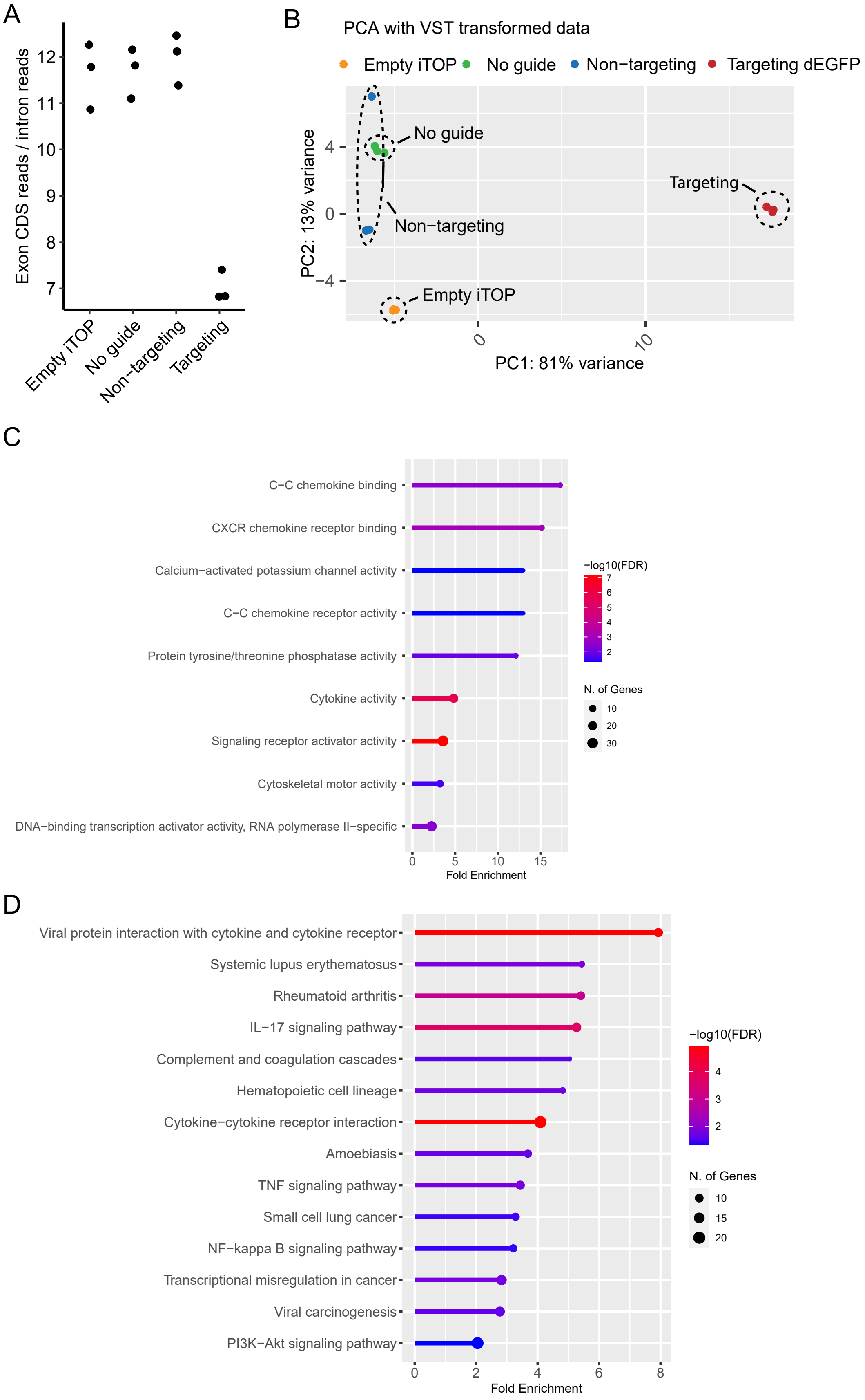


J. F. Bot et al. Supplementary figure 9

### Supplementary figure 10

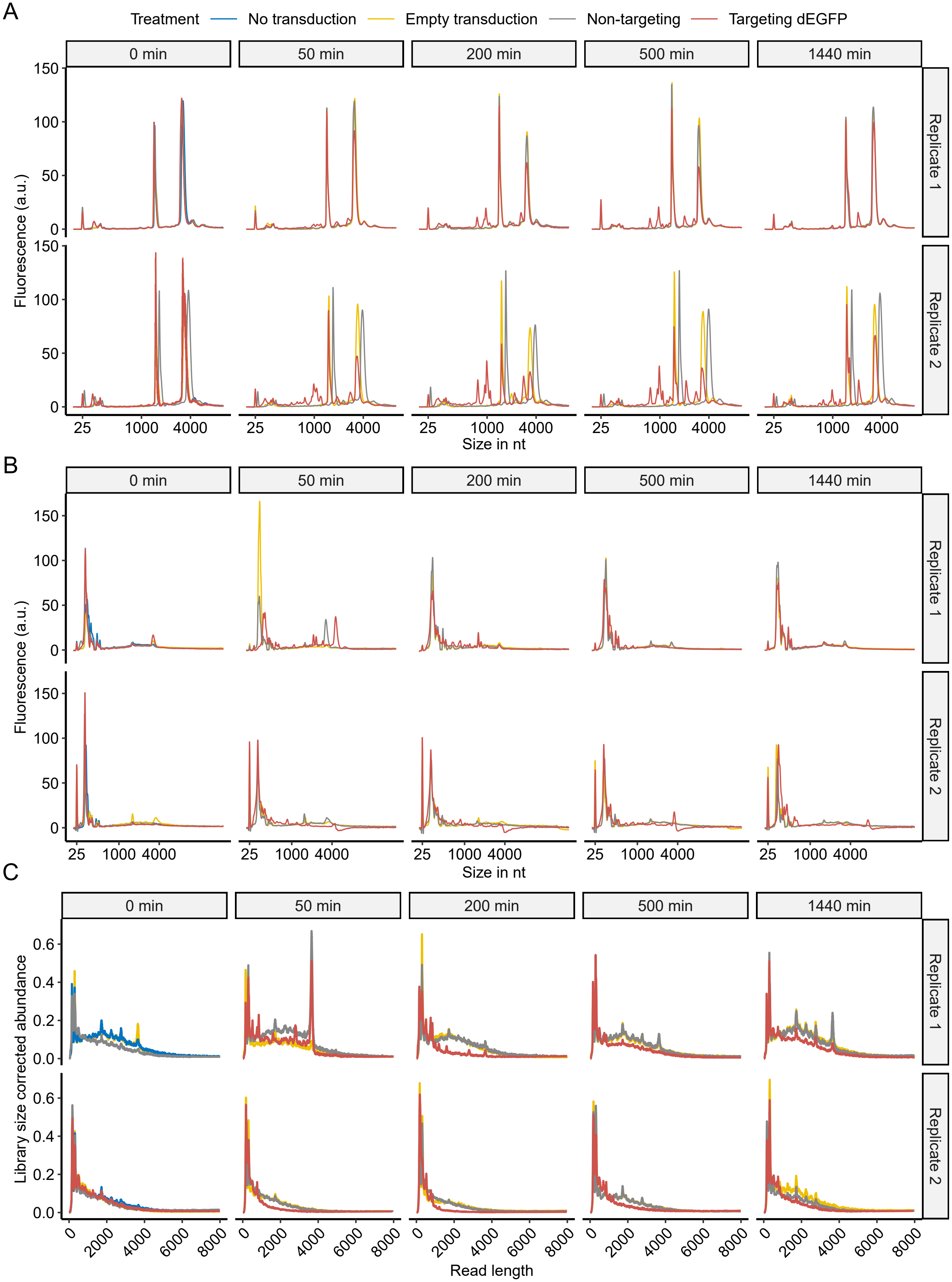


J. F. Bot et al. Supplementary figure 10

### Supplementary figure 11

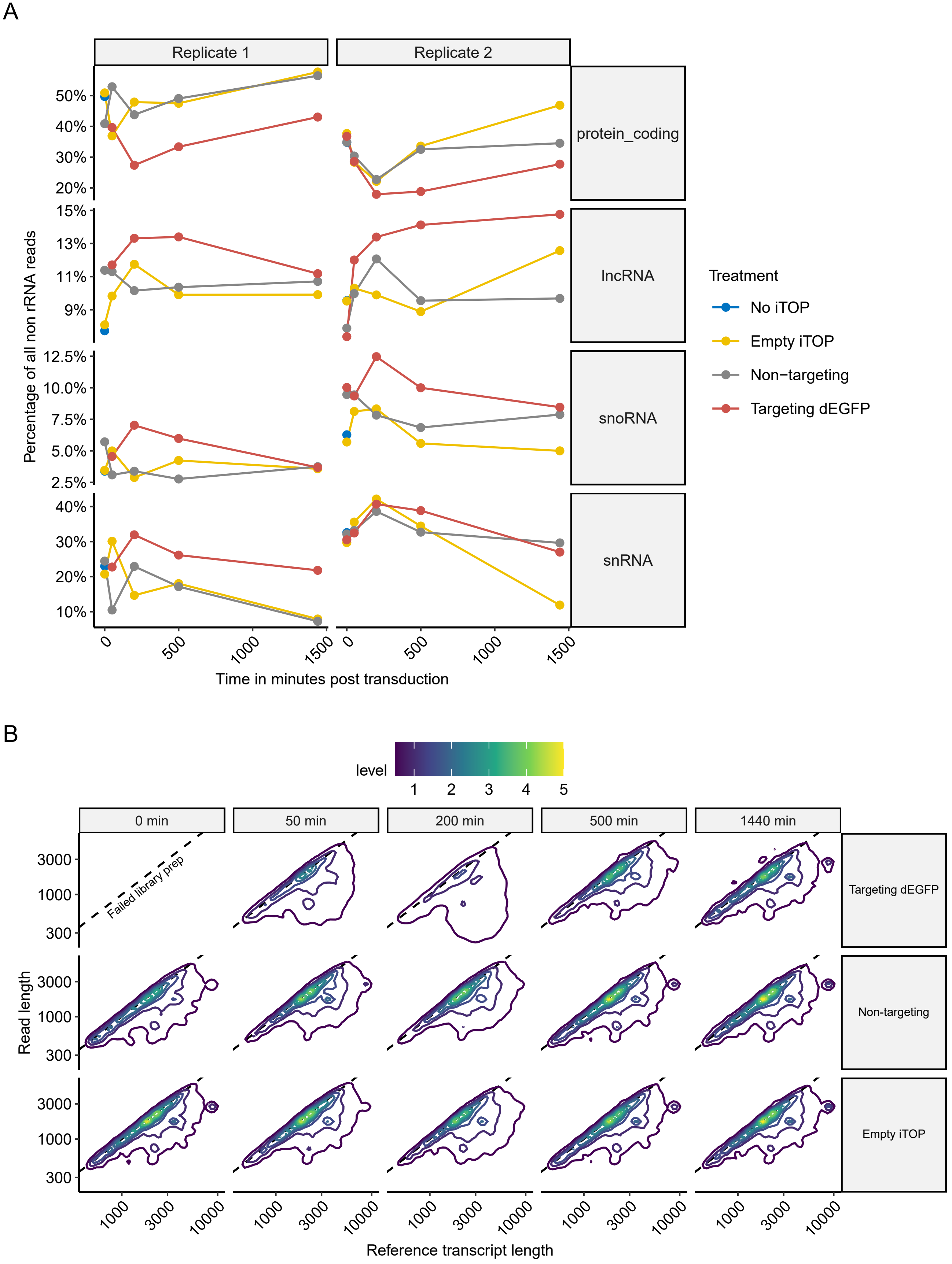


J. F. Bot et al. Supplementary figure 11

### Supplementary figure 12

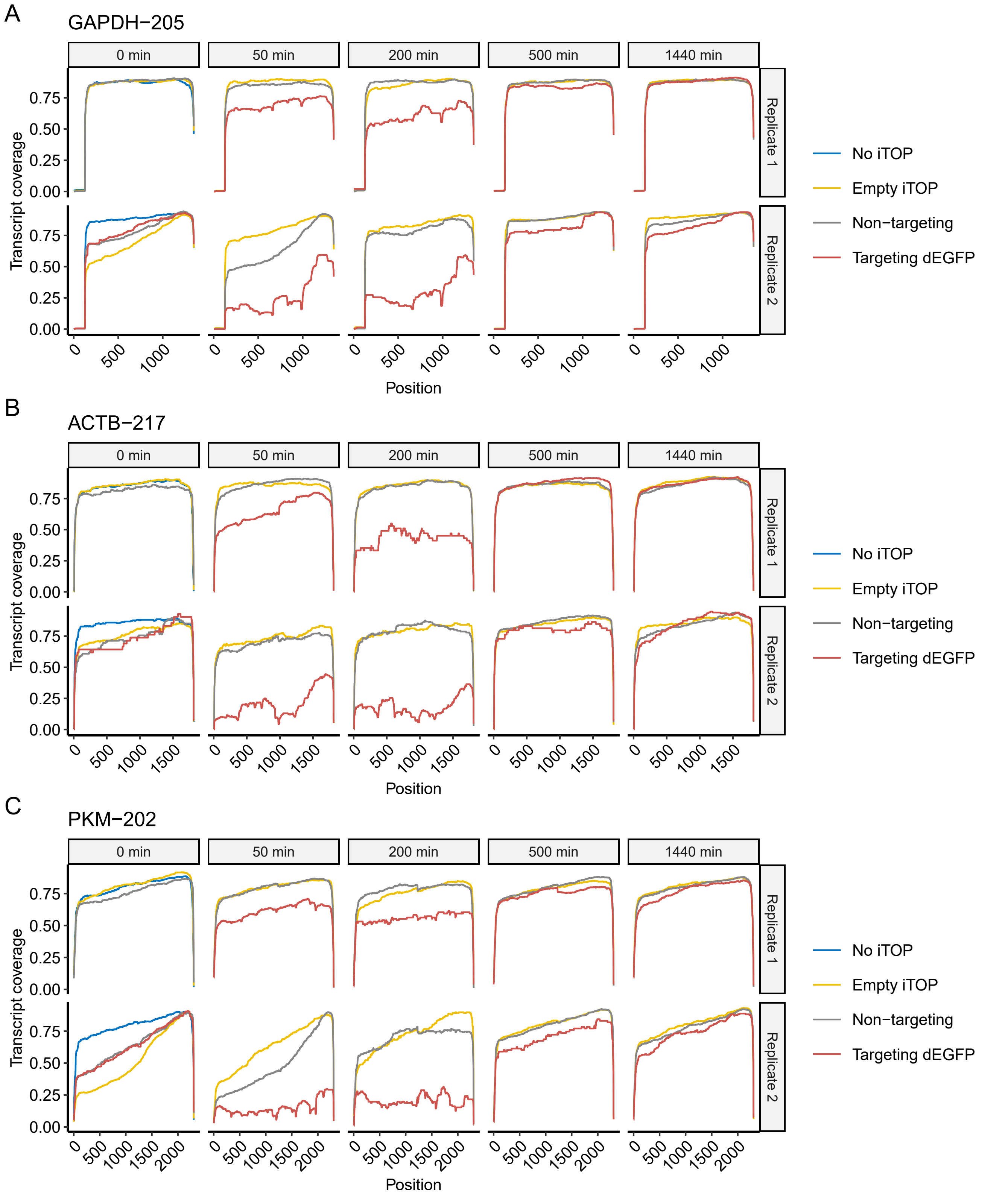


J. F. Bot et al. Supplementary figure 12

### Supplementary figure 13

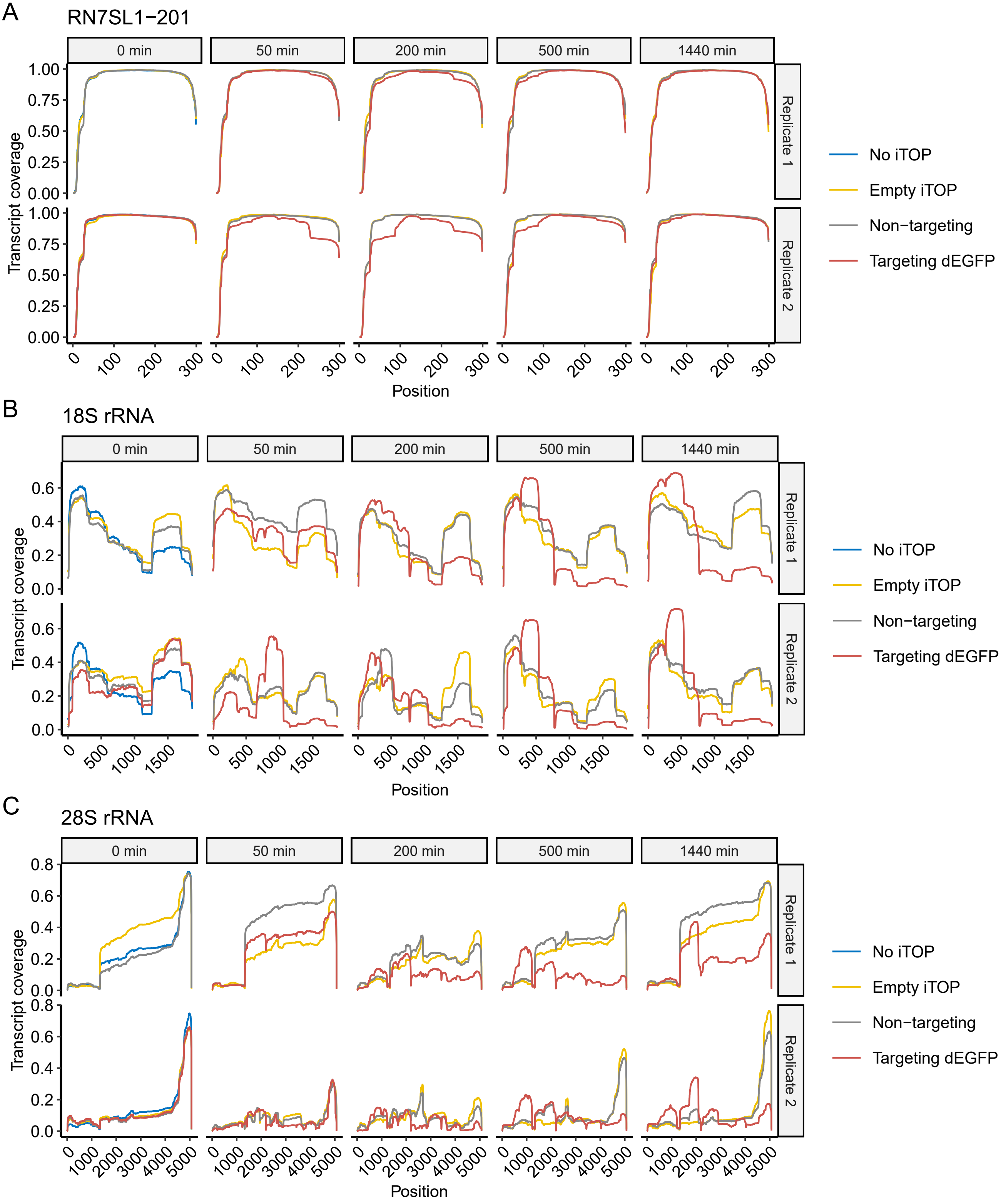


J. F. Bot et al. Supplementary figure 13

### Supplementary figure 14

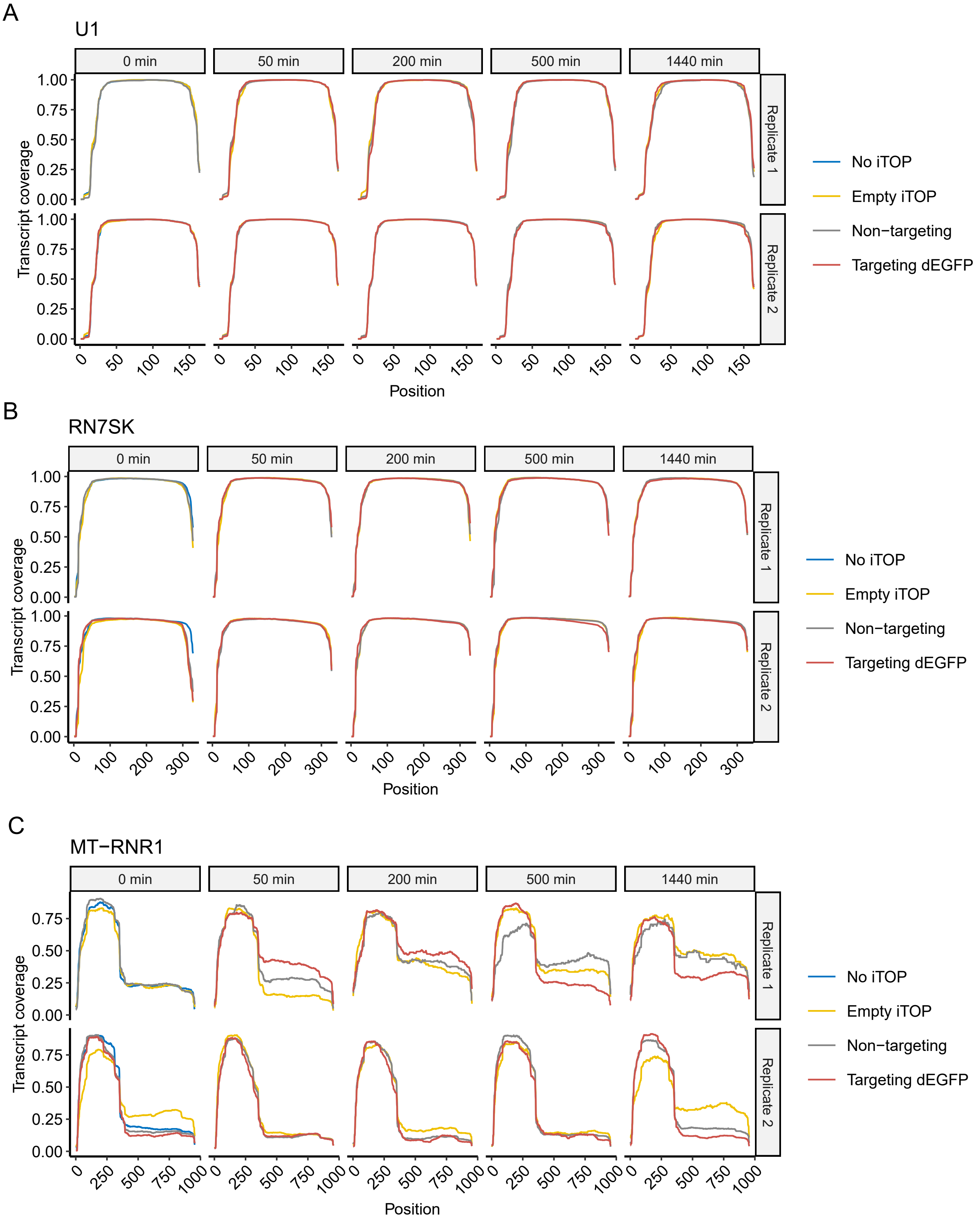


J. F. Bot et al. Supplementary figure 14

### Supplementary figure 15

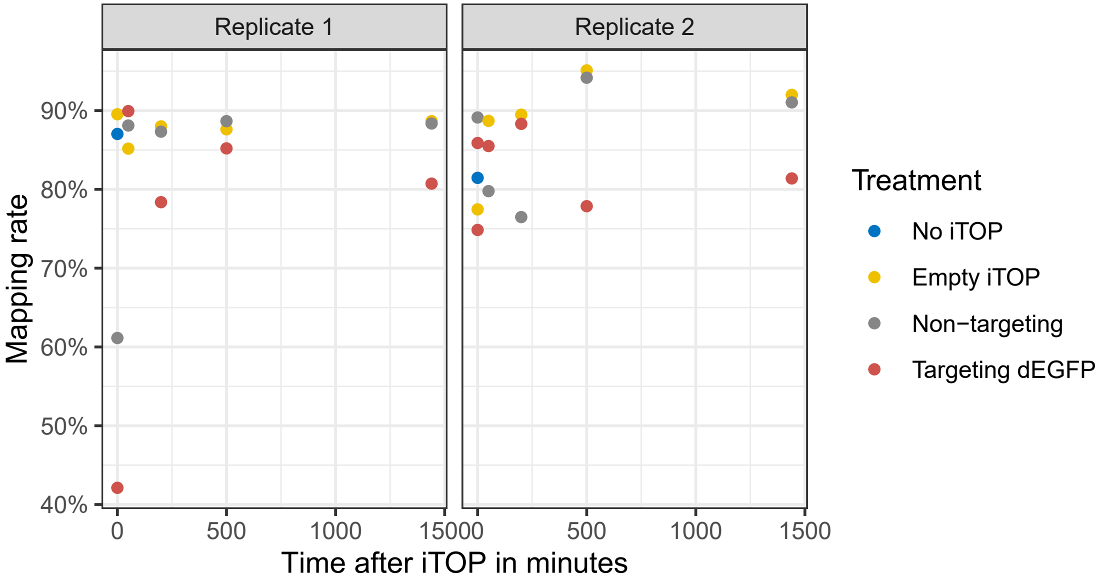

### Supplementary figure 16

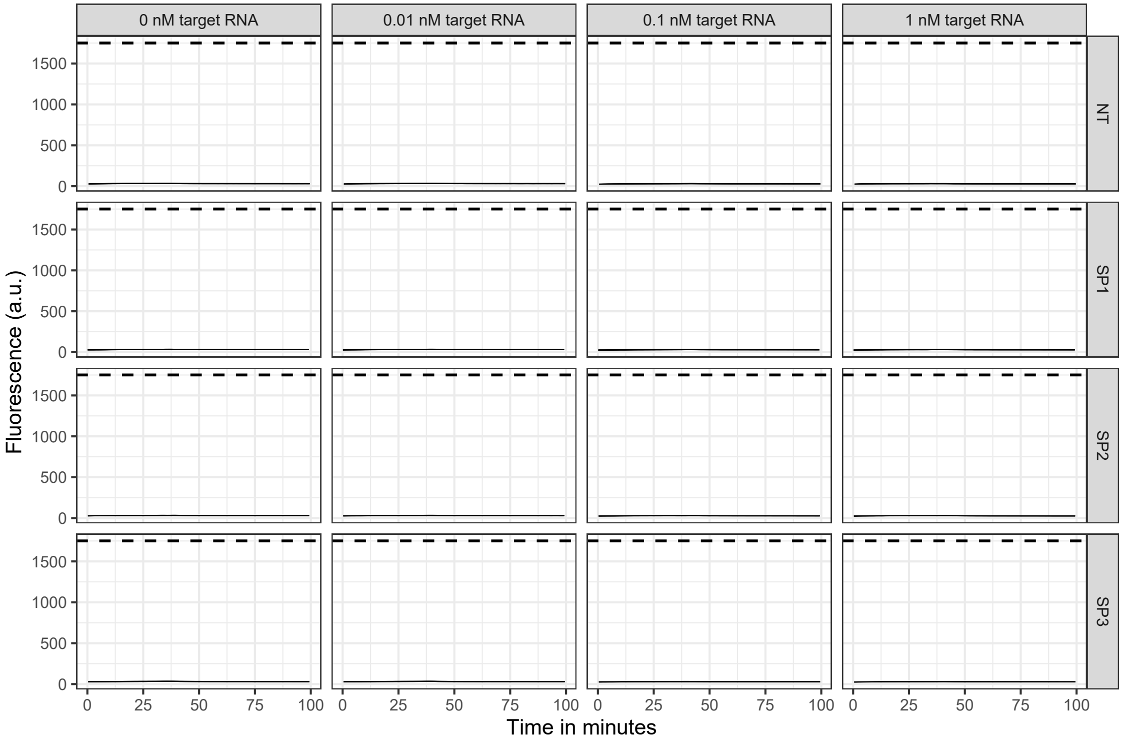

### Supplementary figure 17

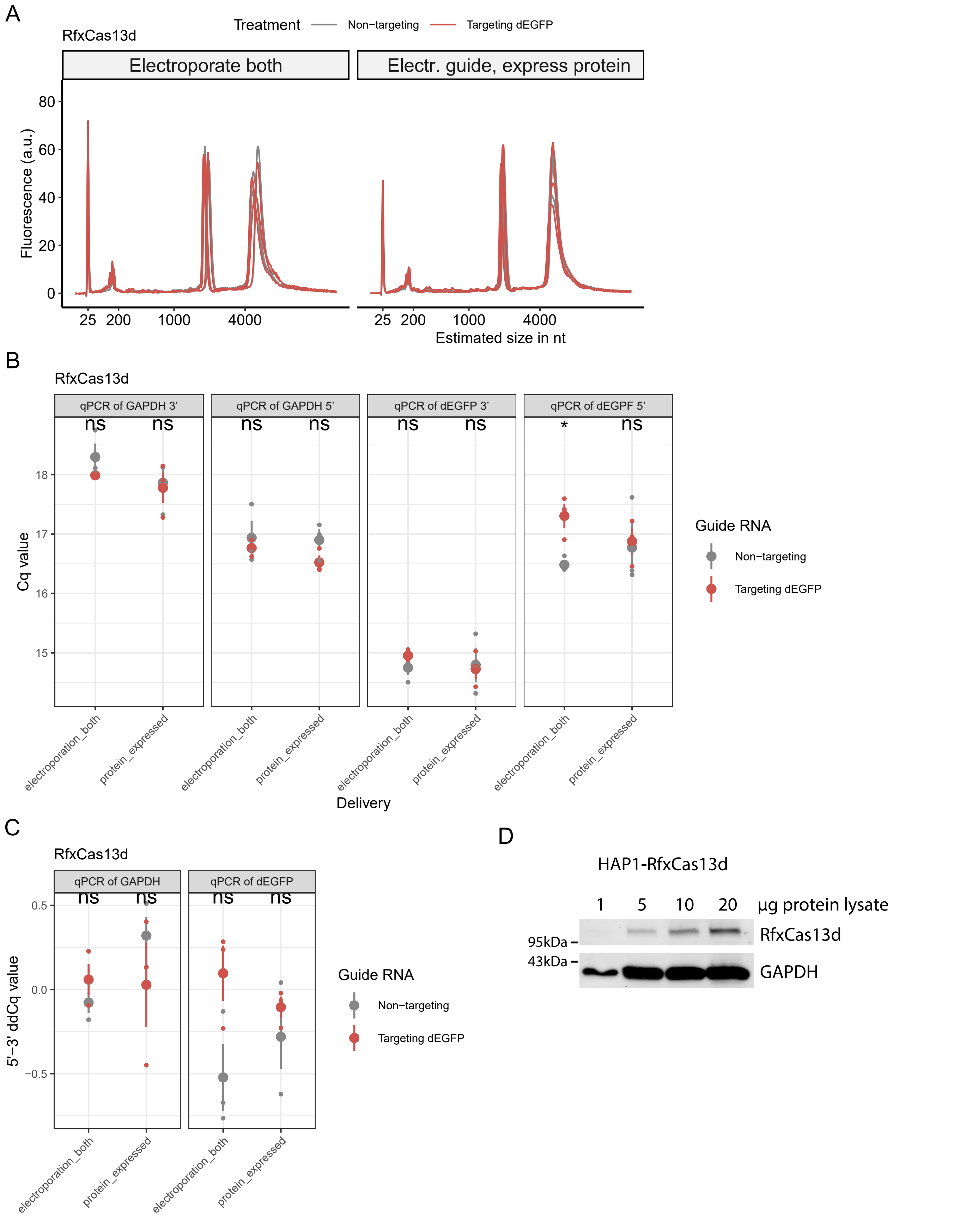


J. F. Bot et al. Supplementary figure 17
